# *In vitro* activity of bedaquiline and imipenem against actively growing, nutrient-starved, and intracellular *Mycobacterium abscessus*

**DOI:** 10.1101/2021.08.06.455495

**Authors:** Olumide Martins, Jin Lee, Amit Kaushik, Nicole C. Ammerman, Kelly E. Dooley, Eric L. Nuermberger

## Abstract

*Mycobacterium abscessus* lung disease is difficult to treat due to intrinsic drug resistance and the persistence of drug-tolerant bacteria. Currently, the standard of care is a multi-drug regimen with at least 3 active drugs, preferably including a β-lactam (imipenem or cefoxitin). These regimens are lengthy, toxic, and have limited efficacy. The search for more efficacious regimens led us to evaluate bedaquiline, a diarylquinoline licensed for treatment of multidrug-resistant tuberculosis. We performed *in vitro* time-kill experiments to evaluate the activity of bedaquiline alone and in combination with the first-line drug imipenem against *M. abscessus* under various conditions. Against actively growing bacteria, bedaquiline was largely bacteriostatic and antagonized the bactericidal activity of imipenem. Contrarily, against nutrient-starved persisters, bedaquiline was bactericidal, while imipenem was not, and bedaquiline drove the activity of the combination. In an intracellular infection model, bedaquiline and imipenem had additive bactericidal effects. Correlations between ATP levels and the bactericidal activity of imipenem and its antagonism by bedaquiline were observed. Interestingly, the presence of Tween 80 in the media affected the activity of both drugs, enhancing the activity of imipenem and reducing that of bedaquiline. Overall, these results show that bedaquiline and imipenem interact differently depending on culture conditions. Previously reported antagonistic effects of bedaquiline on imipenem were limited to conditions with actively multiplying bacteria and/or the presence of Tween 80, whereas the combination was additive or indifferent against nutrient-starved and intracellular *M. abscessus*, where promising bactericidal activity of the combination suggests it may have a role in future treatment regimens.

## Introduction

Lung disease caused by *Mycobacterium abscessus* infection is difficult to treat. Currently, the recommended treatment for *M. abscessus* lung infections is a multi-drug regimen comprised of at least 3 drugs with *in vitro* activity and preferably including imipenem or cefoxitin (1). This regimen, which can be administered for months to years, results in cure in only about 50% of patients and is plagued by problems with drug toxicity and poor tolerability (2). Thus, there is an urgent need to improve treatment of *M. abscessus* lung infections. One approach to identify new treatment options is to evaluate drugs that are effective for other mycobacterial infections, with bedaquiline being a good candidate for repurposing against *M. abscessus*.

Traditional treatment regimens for multidrug-resistant tuberculosis (MDR-TB) were 18-24 months in duration and associated with about 50% cure rates (4), a situation comparable to the current standard of care for *M. abscessus* lung disease. Bedaquiline -- a diarylquinoline approved for treatment of MDR-TB-- is a key component of a new, 6-month MDR-TB regimen (bedaquiline-pretomanid-linezolid or “BPaL”) with a demonstrated 90% cure rate (3). Using mouse models, each individual drug in BPaL was shown to contribute to bacterial killing of the regimen; and bedaquiline specifically was found to contribute significant treatment-shortening activity (5). As several studies have demonstrated that bedaquiline has activity against *M. abscessus*, including previous work by our group showing that bedaquiline was highly bactericidal against non-replicating *M. abscessus* populations (6, 7), we considered that bedaquiline has potential for both improving and shortening treatment of *M. abscessus* lung disease.

Understanding how bedaquiline combines with current first-line drugs is an important step in characterizing bedaquiline’s potential as part of a shorter, curative regimen for *M. abscessus* lung disease. To that end, the objective of this study was to evaluate the activity of bedaquiline-imipenem combinations against *M. abscessus.* Interestingly, Lindman and Dick recently reported that bedaquiline antagonized imipenem’s *in vitro* bactericidal activity against actively growing *M. abscessus* (8). However, Le Moigne *et al.* did not observe antagonism between these drugs in a mouse model of *M. abscessus* lung disease (9). To achieve our objectives, and in light of these recent findings, we conducted a series of *in vitro* studies focused on evaluating the activity of bedaquiline-imipenem combinations across different *M. abscessus* populations, namely actively growing, nutrient-starved non-replicating, and intracellular bacteria. Additionally, we generated resistant mutants to address the impact of bedaquiline resistance on the combined drug activity across these different conditions. This systematic evaluation has produced novel datasets that highlight how experimental conditions can impact drug activity and also provides insight into how bedaquiline and imipenem may be used together for treatment of *M. abscessus* lung infections.

## Results

### Selection and characterization of bedaquiline-resistant *M. abscessus* isolates

We selected and characterized three unique isolates with decreased *in vitro* susceptibility to clofazimine in the *M. abscessus* strain ATCC 19977 (wild-type [WT]) background (**Table S1**). Each of these isolates contained a mutation in *MAB_2299c*, a gene encoding a TetR-family transcriptional repressor of MmpS-MmpL efflux pump systems that confers cross-resistance to clofazimine and bedaquiline when inactivated (10). Indeed, each of our *MAB_2299c* mutant isolates had reduced susceptibility to bedaquiline. The two isolates with *MAB_2299c* mutations that were analyzed (OM4 and OM7) for concurrent mutations in *atpE*, which encodes the ATP synthase subunit targeted by bedaquiline (11), had no such mutations. None of the isolates had mutations in *MAB_4384*, another gene reported to be associated with bedaquiline resistance in *M. abscessus* (12). In cation-adjusted Mueller-Hinton broth (CAMHB), the minimum inhibitory concentration (MIC) of bedaquiline increased 4- or 8-fold to 0.25-0.5 µg/mL against the *MAB_2299c* mutants compared to the wild-type parent strain (MIC 0.0625 µg/mL). Although neither the Clinical & Laboratory Standards Institute (CLSI) nor the manufacturer have defined testing standards for bedaquiline against *M. abscessus* (13, 14), MICs in this range were previously characterized as “bedaquiline-resistant” for *M. abscessus* isolates (10, 15). Therefore, we refer to our *MAB_2299c* mutants as bedaquiline-resistant strains. Isolate OM7, which contained a 150 bp deletion starting at position -28 upstream of the *MAB_2299c* start codon (**Fig. S1**), was utilized in experiments described in this report.

### Assessment of bedaquiline and imipenem/avibactam against *M. abscessus* WT and OM7 mutant in nutrient-rich conditions with Tween 80

We first evaluated the activity of bedaquiline and imipenem alone against actively multiplying *M. abscessus* ATCC 19977 WT parent and OM7 mutant populations in CAMHB supplemented with 0.05% (vol/vol) Tween 80, a surfactant included to reduce the impact of mycobacterial clumping on quantification of colony-forming units (CFUs)(16, 17). Bedaquiline alone exerted only bacteriostatic activity, limiting growth at concentrations ≥0.0625 and ≥2 µg/mL against WT and OM7, respectively (**Fig. 1A-B**; see **Table S2** for all CFU data). In contrast, imipenem alone limited growth at 1 µg/mL and had strong bactericidal activity at concentrations ≥2 µg/mL against both strains after 3 days of drug exposure (**Fig. 1C-D**). We also evaluated the activity of imipenem in combination with avibactam, a β-lactamase inhibitor that enhances the susceptibility of *M. abscessus* to β-lactams, including imipenem (18, 19). In combination with avibactam at 2 µg/mL, the imipenem concentration needed to inhibit growth decreased 2-fold to 1 µg/mL for both strains (**Fig. 1E-F**).

**Fig. 1.**
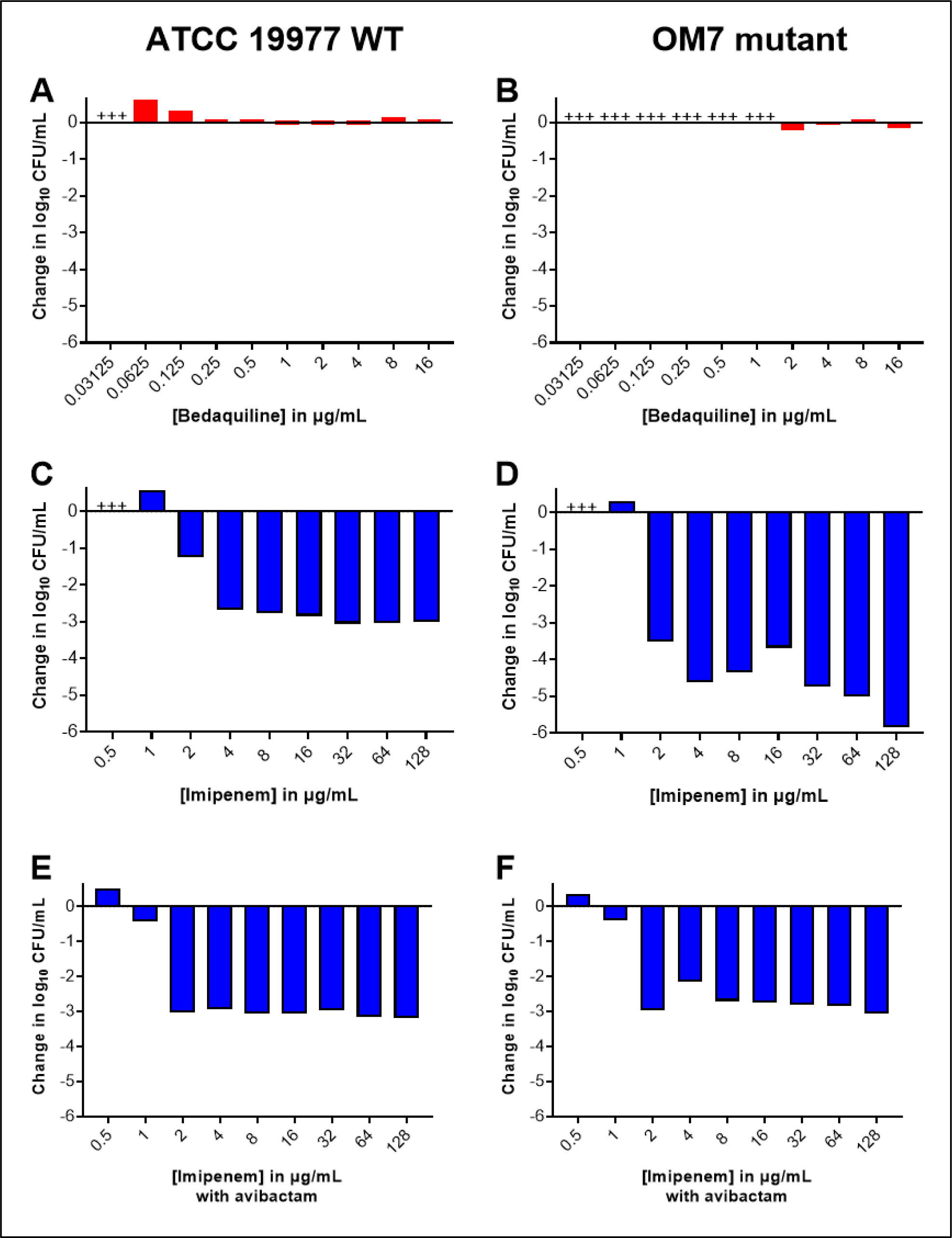
Activity of bedaquiline (A-B), imipenem (C-D), and imipenem with avibactam (E-F) against *M. abscessus* ATCC 19977 WT (A, C, E) and OM7 mutant (B, D, F) in CAMHB with 0.05% Tween 80. The change in log_10_ CFU/mL after 3 days of drug exposure relative to Day 0 is presented for each drug/strain set. Avibactam concentration in panels E-F was 2 µg/mL. +++ indicates bacterial overgrowth and clumping that precluded CFU determination. Avibactam alone at concentrations from 0.5 to 64 µg/mL had no anti-*M. abscessus* activity; bacteria grew/clumped at all avibactam concentrations, precluding CFU determination. The Day 0 bacterial concentrations (in log_10_ CFU/mL) for each panel were as follows: A) 5.96; B) 6.11; C) 5.85; D) 5.86; E) 5.85; F) 5.86. CFU values are provided in **Table S2**.

Next, we evaluated the activity of bedaquiline and imipenem/avibactam combinations against actively growing *M. abscessus* WT and OM7 mutant strains. As previously observed, bedaquiline alone at 0.5 and 1 µg/mL prevented growth of the WT strain and permitted growth of the OM7 mutant, and imipenem/avibactam combinations of 2/2 and 4/2 µg/mL were bactericidal against both bacterial strains (**Fig. 2**; **Table S3**). However, the addition of bedaquiline to the imipenem/avibactam combinations abolished this bactericidal activity.

**Fig. 2.**
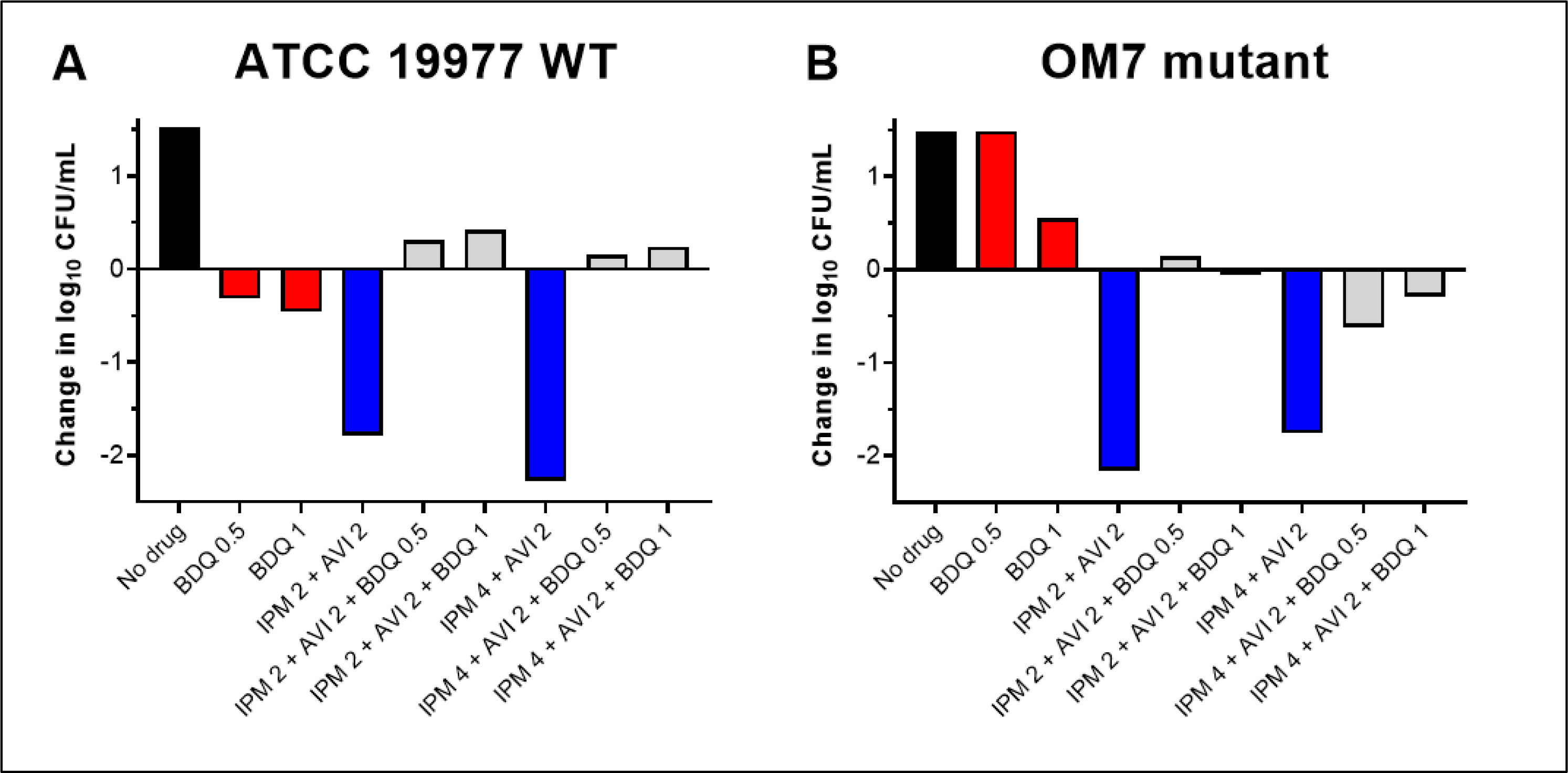
Activity of bedaquiline (BDQ) and imipenem (IPM)/avibactam (AVI) combinations against *M. abscessus* ATCC 19977 WT (A) and OM7 mutant (B) strains in CAMHB with 0.05% Tween 80. The change in log_10_ CFU/mL after 3 days of drug exposure relative to Day 0 is presented for each drug/strain set. Black bars represent the no drug control; red bars indicate BDQ only samples; blue bars represent IPM/AVI only samples, and gray bars indicate IPM/AVI plus BDQ samples. The number after each drug abbreviation represents the concentration in µg/mL. The Day 0 bacterial concentrations (in log_10_ CFU/mL) were 5.57 and 5.76 for WT and OM7 mutant strains, respectively. CFU values are provided in **Table S3**.

Against both strains, the activity of bedaquiline alone appeared concentration-dependent, but this relationship was less clear when combined with imipenem/avibactam. Therefore, we next examined the concentration-ranging activity of bedaquiline when added to imipenem/avibactam combinations against *M. abscessus* WT and OM7. Against both strains, bedaquiline/imipenem/avibactam combinations were more bactericidal when bedaquiline was included at concentrations ≤0.0156 µg/mL, and this bactericidal activity was bedaquiline concentration-dependent (**Fig. S2**; **Table S4**), with bactericidal activity declining with increasing bedaquiline concentration. Overall, the magnitude of killing associated with bedaquiline/imipenem/avibactam combinations was greater against the OM7 mutant than against the WT strain.

### **Assessment of bedaquiline combined with other cell wall synthesis inhibitors** (and non cell wall synthesis inhibitors) **against *M. abscessus* WT in nutrient-rich conditions with Tween 80**

Our data indicated that bedaquiline antagonized the bactericidal activity of imipenem/avibactam against both WT parent and OM7 mutant strains and suggested that this antagonism might only occur at concentrations at which bedaquiline alone was able to limit bacterial growth. Other groups have reported antagonism between bedaquiline and cell wall synthesis inhibitors, including imipenem as well as non-β-lactam inhibitors, against *M. abscessus* and other mycobacteria (8, 20-22), but these previous studies did not include bedaquiline at sub-inhibitory concentrations. Therefore, we examined the concentration-ranging impact of bedaquiline on the bactericidal activity of two additional cell wall synthesis inhibitors: the β-lactam meropenem and N-4S-methylcyclohexyl-4,6-dimethyl-1H-indole-2-carboxamide, a non-β-lactam MmpL3 inhibitor with demonstrated *in vitro* activity against *M. abscessus* (23). We also evaluated the impact of bedaquiline on the bactericidal activity of clarithromycin, a protein synthesis inhibitor. Meropenem, the MmpL3 inhibitor, and clarithromycin were used at concentrations expected to have activity against *M. abscessus* WT: 16, 2, and 4 µg/mL, respectively (6, 23, 24), and meropenem was used without avibactam to allow for more direct comparisons between the different groups. Because these agents are relatively more stable in aqueous media than imipenem, this experiment was extended out to 8 days. Again, bedaquiline alone had limited activity against actively growing WT bacteria, resulting in bacteriostasis at 0.125 µg/mL and a weak bactericidal effect at concentrations of 1-2 µg/mL (**Fig. 3A**; see **Table S5** for all CFU data from this figure). When combined with meropenem, these bedaquiline concentrations antagonized the bactericidal activity of meropenem over 8 days, while the lower bedaquiline concentrations of 0.0039 and 0.0156 µg/mL that were largely inactive on their own were not antagonistic but rather contributed additional killing compared to meropenem alone at Day 8 (**Fig. 3B**). When combined with the MmpL3 inhibitor, bedaquiline at concentrations ≥0.0156 µg/mL abrogated the bactericidal activity compared to the MmpL3 inhibitor alone, while bedaquiline added at 0.0039 µg/mL remained highly bactericidal, although to a lesser magnitude than the MmpL3 inhibitor alone (**Fig. 3C**). In contrast, bedaquiline did not diminish the bactericidal activity of clarithromycin and added activity at concentrations ≥0.125 µg/mL at Day 8 (**Fig. 3D**).

**Fig. 3.**
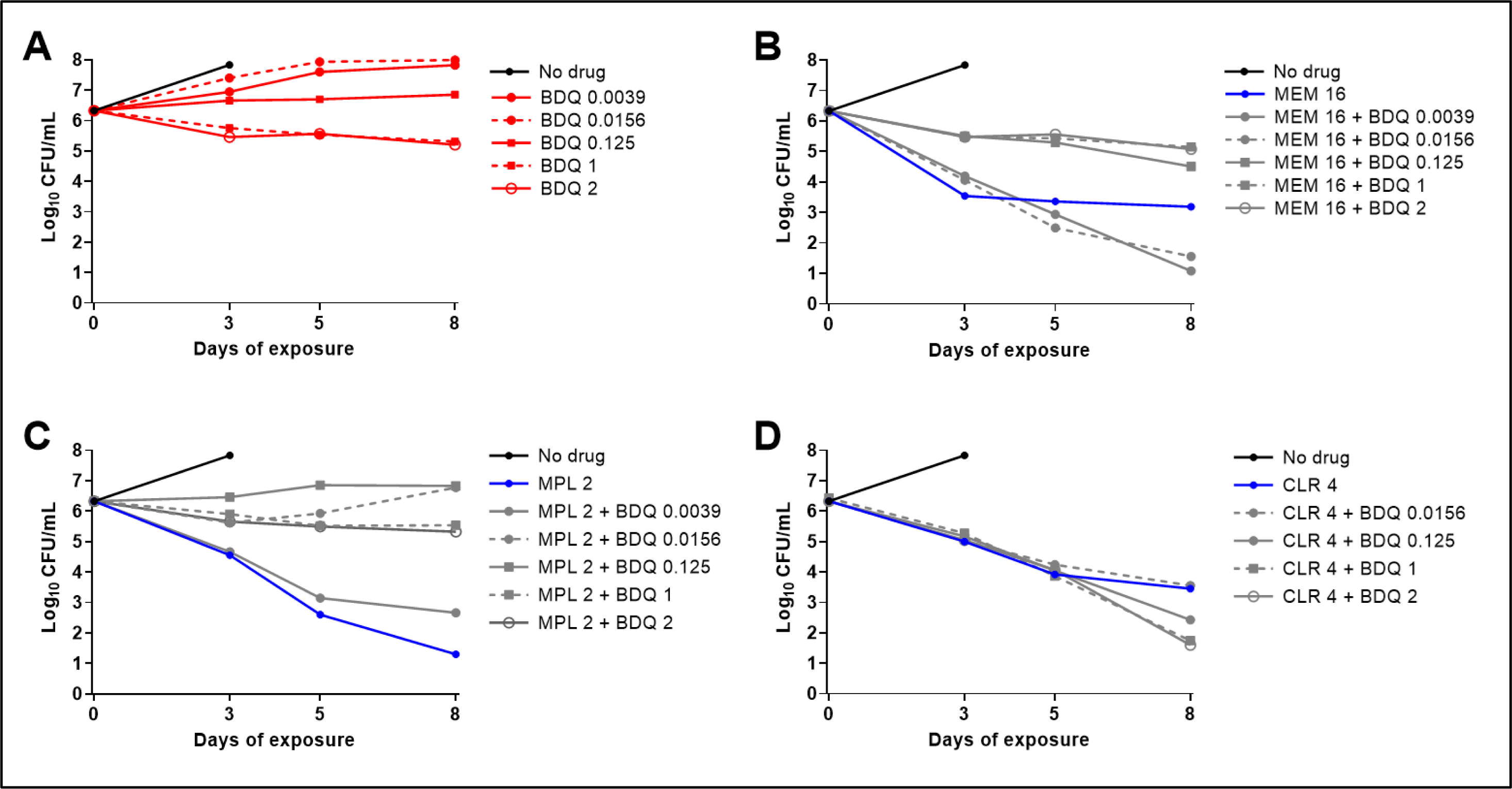
Activity of bedaquiline (BDQ) alone (A), BDQ plus meropenem (MEM) (B), BDQ plus MmpL3 inhibitor (MPL) (C), and BDQ plus clarithromycin (CLR) (D) against *M. abscessus* ATCC 19977 WT in CAMHB with 0.05% Tween 80. The number after each drug abbreviation represents the concentration in µg/mL. MPL is N-4S-methylcyclohexyl-4,6-dimethyl-1H-indole-2-carboxamide. The bacteria in the no drug control overgrew and clumped after Day 3, precluding CFU quantification. CFU values are provided in **Table S5**.

### Correlation of ATP levels with bedaquiline/imipenem bactericidal activity against *M. abscessus* WT in nutrient-rich conditions with Tween 80

Previous work by Lindman and Dick demonstrated that the bactericidal activity of imipenem correlated with increased *M. abscessus* ATP production, which was also abrogated with co-exposure to bedaquiline (8). Therefore, we next evaluated the relative bacterial ATP levels associated with bedaquiline and imipenem against actively growing WT bacteria in CAMHB with Tween 80. To reduce study variables, avibactam was omitted. In both biological replicates, we observed that imipenem’s potent bactericidal activity was unchanged or even increased when bedaquiline was added at concentrations of 0.0039 or 0.0078 µg/mL, yet adding bedaquiline at concentrations ≥0.0625 µg/mL nearly eliminated imipenem’s bactericidal activity (**Fig. 4A,B**; **Table S6**). The relative levels of bacterial ATP, measured as relative light units (RLU) adjusted by CFU count, correlated with the bactericidal effect; ATP levels increased when bacteria were exposed to bactericidal imipenem/bedaquiline concentrations, and ATP levels decreased as the bedaquiline concentration increased (**Fig. 4C,D**). RLU data not adjusted by CFU counts are presented in **Fig. S3**.

**Fig. 4.**
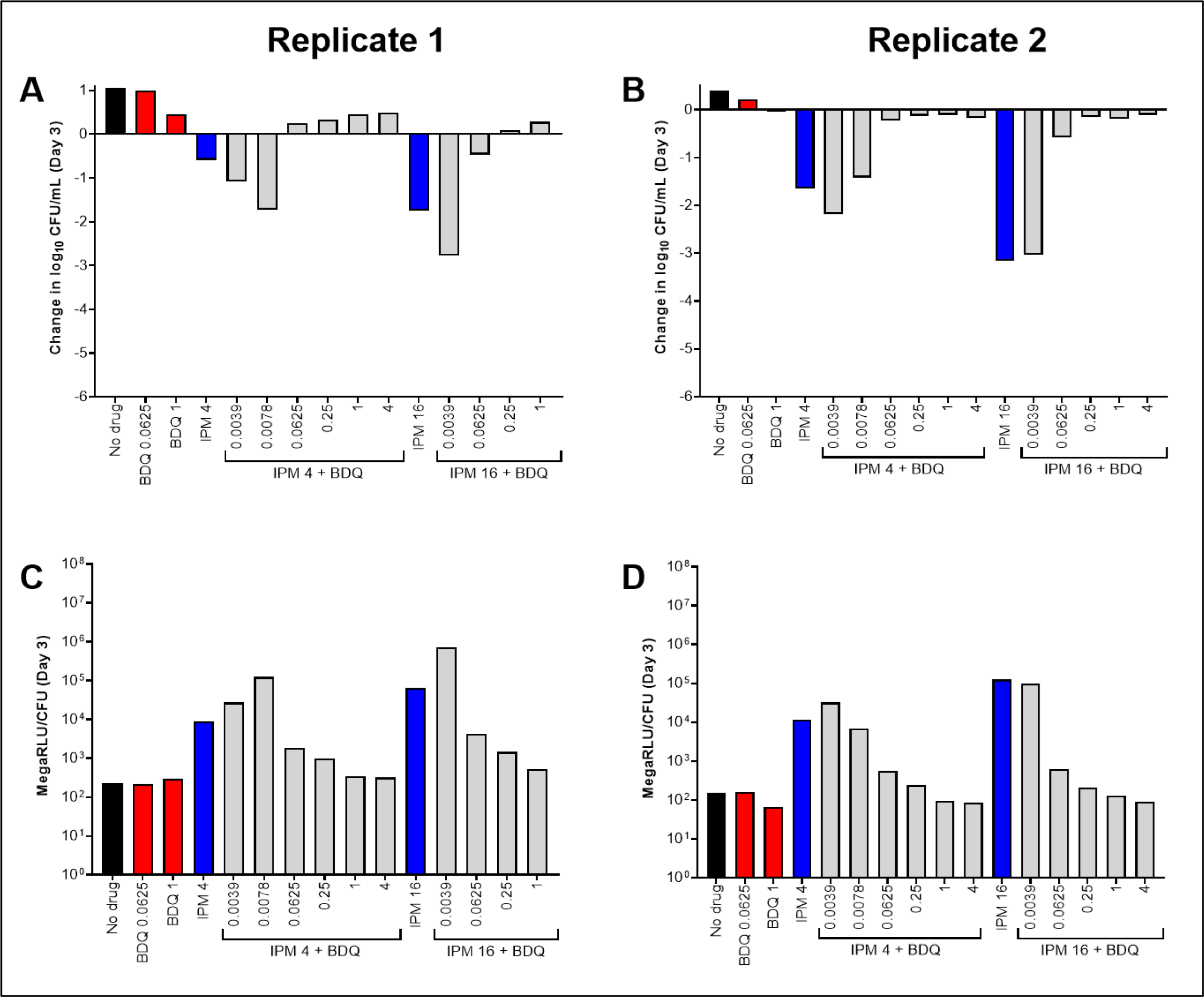
Activity of bedaquiline (BDQ) and imipenem (IPM) against *M. abscessus* ATCC 19977 WT (A,B) and relative bacterial ATP levels (C,D), in CAMHB with 0.05% Tween 80. The change in log_10_ CFU/mL after 3 days of drug exposure relative to Day 0 is presented for each biological replicate in panels A and B. CFU-adjusted ATP levels, indicated by relative light units (RLU)/CFU from samples after 3 days of drug exposure, are presented for each biological replicate in panels C and D. Black bars represent the no drug control; red bars indicate BDQ only samples; blue bars represent IPM only samples, and gray bars indicate IPM plus BDQ samples. The number after each drug abbreviation represents the concentration in µg/mL (for gray bars the BDQ concentration in µg/mL is presented under each bar). RLU/mL for all samples (not adjusted for CFUs) are presented in **Fig. S3**. The Day 0 bacterial concentration for panels A and B was 6.25 and 7.03 log_10_ CFU/mL, respectively. CFU and RLU values are provided in **Table S6**.

### Assessment of bedaquiline and imipenem activity against *M. abscessus* ATCC 19977 WT and OM7 in nutrient-rich conditions without Tween 80

Thus far, the surfactant Tween 80 was present in all experimental conditions. As noted earlier, this was done in an effort to reduce bacterial clumping. However, as bacterial clumping was not always prevented (**Fig. 1**; **Fig. 3**; **Table S3**; **Table S4**), and as surfactants such as Tween 80 can influence cell wall permeability and drug susceptibility (25, 26), we also evaluated the activity of bedaquiline and imipenem against *M. abscessus* in CAMHB without Tween 80. After 3 days in these assay conditions, bedaquiline completely inhibited the growth of WT bacteria starting at 0.0312 µg/mL and exhibited bactericidal activity (reducing CFU by ∼0.5-0.9 log_10_ CFU/mL) at concentrations ≥0.0625 µg/mL (**Fig. 5A**; see all CFU data in **Table S7**). Bedaquiline only limited growth of the OM7 mutant at concentrations ≥0.5 µg/mL (**Fig. 5B**). Therefore, against both WT and OM7 populations, bedaquiline was more active (on a µg/mL basis) in CAMHB without Tween 80 than that with Tween 80 (**Fig. 1A,B**). In contrast, the activity of imipenem was reduced when Tween 80 was omitted from the media. After 3 days of exposure, imipenem inhibited growth of WT and OM7 at 16 and 4 µg/mL, respectively (**Fig. 5C,D**). Although imipenem was tested up to 256 µg/mL, the magnitude of killing never exceeded 2 log_10_ CFU/mL against either strain, while in CAMHB with 0.05% Tween 80, CFU reductions exceeded 2 log_10_ CFU/mL at 2-4 µg/mL imipenem (**Fig. 1C,D**).

**Fig. 5.**
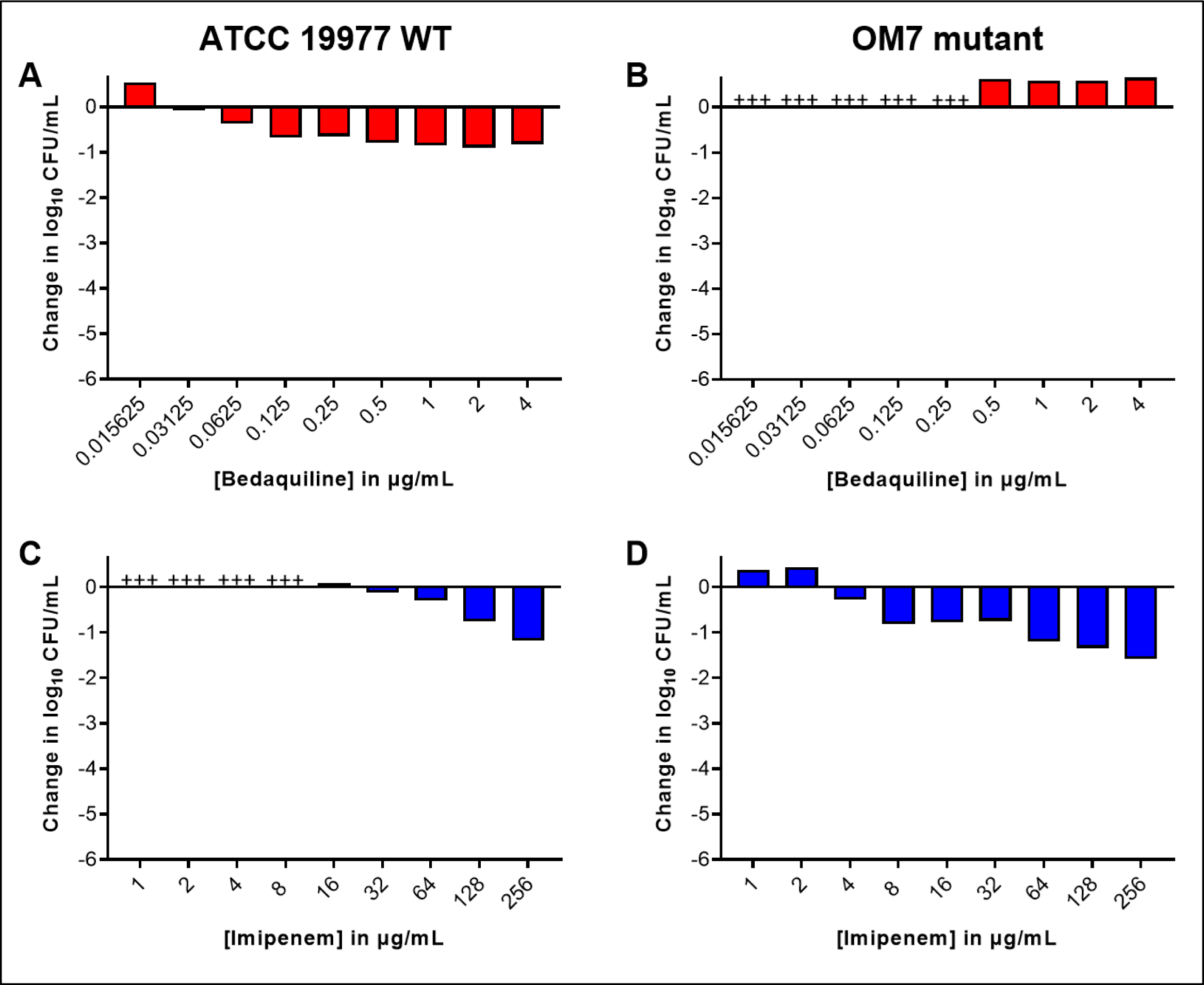
Activity of bedaquiline (A,B) and imipenem (C,D) against *M. abscessus* ATCC 19977 WT (A,C) and OM7 mutant (B,D) in CAMHB without Tween 80. The change in log_10_ CFU/mL after 3 days of drug exposure relative to Day 0 is presented for each drug/strain set. +++ indicates bacterial growth and clumping that precluded CFU determination. The Day 0 bacterial concentrations (in log_10_ CFU/mL) for each panel were as follows: A) 5.90; B) 5.08; C) 5.91; D) 6.20. CFU values are provided in **Table S7.**

We next assessed the activity of imipenem/bedaquiline combinations in CAMHB without Tween 80. Again, we observed that, relative to activity in Tween 80-containing media, bedaquiline alone had increased killing and imipenem alone had decreased killing in the absence of Tween 80 (**Fig. 6A,B** (replicate 1); **Fig. S4A** (replicate 2); see all CFU data in **Tables S8-9**). For both WT and OM7 strains, the addition of bedaquiline decreased imipenem’s bactericidal activity, and lower concentrations of bedaquiline combined with imipenem did not increase bactericidal activity, as was observed in Tween 80-containing media (**Fig. 4**; **Fig. S2**). In these assay conditions, relative bacterial ATP levels also correlated with bacterial killing (**Fig. 6C,D**; **Fig. S4B**). ATP levels, not adjusted by CFU counts, indicate that ATP levels in the samples that overgrew and clumped (precluding CFU quantification) were similar to the ATP levels of the no drug control samples for both WT and OM7 populations (**Fig. S4C**; **Fig. S5**; see all RLU data in **Tables S8-S9**).

**Fig. 6.**
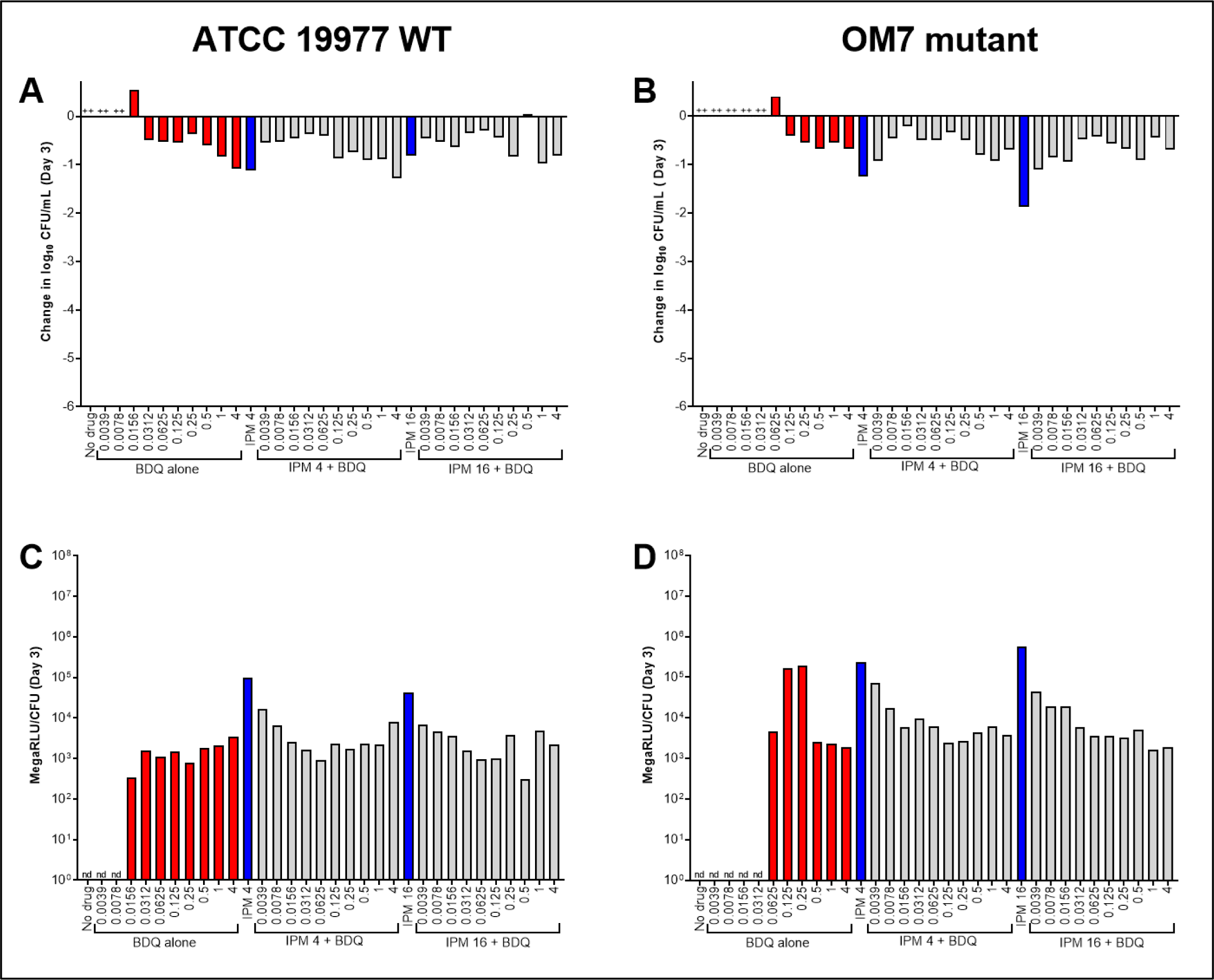
Activity of bedaquiline (BDQ) and imipenem (IPM) against *M. abscessus* ATCC 19977 WT (A) and OM7 mutant (B), with relative bacterial ATP levels (C,D), in CAMHB without Tween 80. The changes in log_10_ CFU/mL after 3 days of drug exposure relative to Day 0 are presented in panels A and B; CFU-adjusted ATP levels, indicated by relative light units (RLU)/CFU from samples after 3 days of drug exposure, are presented in panels C and D. Red bars indicate BDQ only samples; blue bars represent IPM only samples, and gray bars indicate IPM + BDQ samples. The number after the IPM abbreviation represents the concentration in µg/mL; for red and gray bars, the BDQ concentration in µg/mL is under each bar. ++ indicates that the bacterial overgrew and clumped, precluding CFU quantification. nd indicates not determined (due to lack of CFU count data). RLU/mL for all samples (not adjusted for CFUs) are presented in **Fig. S5**. The Day 0 bacterial concentration for panels A and B was 5.62 and 5.70 log_10_ CFU/mL, respectively. CFU and RLU values are provided in **Table S8**.

### Evaluation of bedaquiline and imipenem activity against nutrient-starved *M. abscessus* ATCC 19977 WT and OM7 in PBS without Tween 80

Regardless of the presence or absence of Tween 80, imipenem alone was clearly more active than BDQ alone against actively growing *M. abscessus* strains in nutrient-rich media, a finding that aligns with previously published work (6-8). However, we previously reported the converse against nutrient-starved *M. abscessus*; namely, that bedaquiline is highly bactericidal while imipenem has no observable activity (6). Therefore, we next evaluated the bactericidal activity of imipenem/bedaquiline combinations against *M. abscessus* WT and OM7 populations that had been nutrient-starved in PBS for 14 days prior to drug exposure. After nutrient starvation, drug activity assays were performed in PBS without Tween 80. As expected, bedaquiline alone had potent, concentration-dependent bactericidal activity against nutrient-starved *M. abscessus* WT (**Fig. 7A**, Day 3; **Fig. S6A**, Day 7). Bedaquiline also exerted concentration-dependent killing against the nutrient-starved OM7 mutant (**Fig. 7B**; **Fig. S6B**), although with a much lower magnitude of killing compared to the WT parent strain (see all CFU data in **Table S10**). Imipenem alone at 4 or 16 µg/mL had no bactericidal activity against either strain, and the bactericidal activity of bedaquiline was unchanged by the addition of imipenem. In these nutrient starvation conditions, bacterial ATP levels adjusted by CFU counts correlated with bactericidal activity for both strains (**Fig. 7C,D**; **Figs. S6C,D**). For both WT and OM7, the total ATP levels, not adjusted by CFUs, decreased with increasing bedaquiline concentration (**Fig. S7**; **Fig. S8**), similar to what was observed in CAMHB without Tween 80 (**Fig. S5**), although with overall lower bacterial ATP levels in nutrient starvation conditions.

**Fig. 7.**
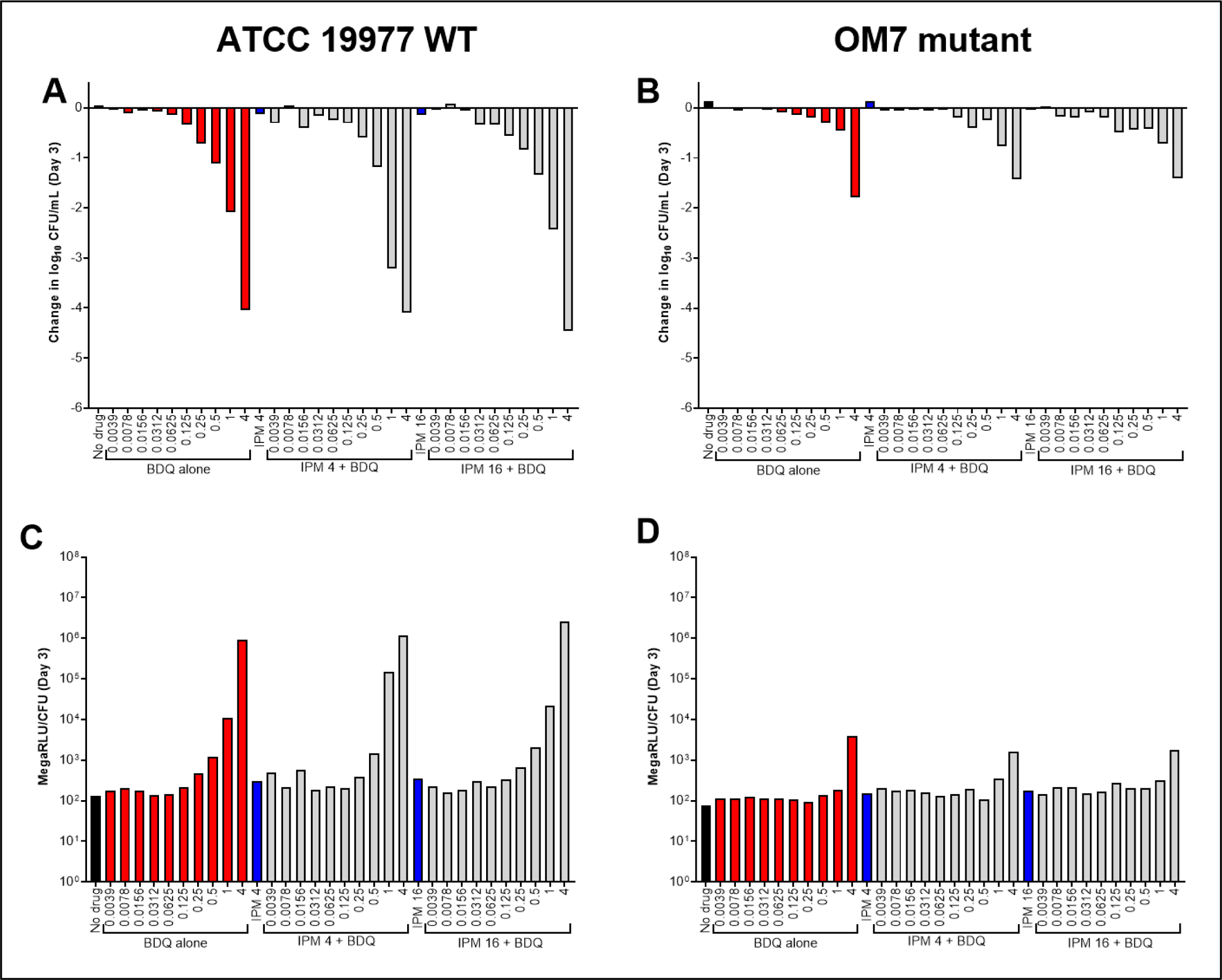
Activity of bedaquiline (BDQ) and imipenem (IPM) against nutrient-starved *M. abscessus* ATCC 19977 WT (A) and OM7 mutant (B), with relative bacterial ATP levels (C,D), in PBS without Tween 80. Bacteria were nutrient-starved in PBS for 14 days prior to drug exposure. The changes in log_10_ CFU/mL after 3 days of drug exposure relative to Day 0 are presented in panels A and B; CFU-adjusted ATP levels, indicated by relative light units (RLU)/CFU from samples after 3 days of drug exposure, are presented in panels C and D. Red bars indicate BDQ only samples; blue bars represent IPM only samples, and gray bars indicate IPM + BDQ samples. The number after the IPM abbreviation represents the concentration in µg/mL; for red and gray bars, the BDQ concentration in µg/mL is under each bar. Data after 7 days of drug exposure are presented in **Fig. S7**. RLU/mL (not adjusted for CFUs) for WT and OM7 are presented in **Fig. S8** and **Fig. S9**, respectively. The Day 0 bacterial concentration for panels A and B was 6.25 and 6.29 log_10_ CFU/mL, respectively. CFU and RLU values are provided in **Table S9**.

### Comparison of bedaquiline and imipenem activity against nutrient-starved *M. abscessus* ATCC 19977 WT in PBS with or without Tween 80

Because the presence of Tween 80 positively impacted the activity of imipenem and negatively impacted the activity of bedaquiline against actively growing *M. abscessus*, we also evaluated the impact of Tween 80 on the bactericidal activity of these drugs, alone and together, against nutrient-starved bacteria in PBS with 0.05% Tween 80. For logistical reasons, bacteria were nutrient starved in PBS for 20 days prior to drug exposure in this experiment, and bacteria were exposed to drugs for up to 4 days in Tween-containing PBS. Although bedaquiline alone still exerted concentration-dependent bactericidal activity in these conditions (**Fig. 8**; **Fig. S9**; see all CFU data in **Table S11**), the magnitude of the killing was greatly reduced compared to its activity in PBS with Tween 80. In PBS with Tween 80, bedaquiline alone at 1 and 4 µg/mL reduced CFU counts by approximately 1 and 1.5 log_10_ CFU/mL, respectively, after 4 days of exposure, while bedaquiline alone at 1 and 4 µg/mL reduced them by approximately 2 and 4 log_10_ CFU/mL, respectively, in 3 days against nutrient-starved WT in PBS without Tween 80 (**Fig. 7A**). Conversely, imipenem alone reduced CFU counts by >3 log_10_ CFU/mL against nutrient-starved bacteria in PBS with Tween 80, which was similar to the magnitude of killing observed in CAMHB with Tween 80 (**Fig. 1C**). When imipenem and bedaquiline were combined, bactericidal activity was reduced compared to imipenem alone; however, the combined killing of imipenem/bedaquiline at higher concentrations of bedaquiline aligned with the bactericidal activity of bedaquiline alone. A head-to-head-comparison in PBS with or without Tween against WT bacteria that were nutrient-starved for 14 days prior to drug exposure confirmed these findings (**Fig. S10**; **Table S12**).

**Fig. 8.**
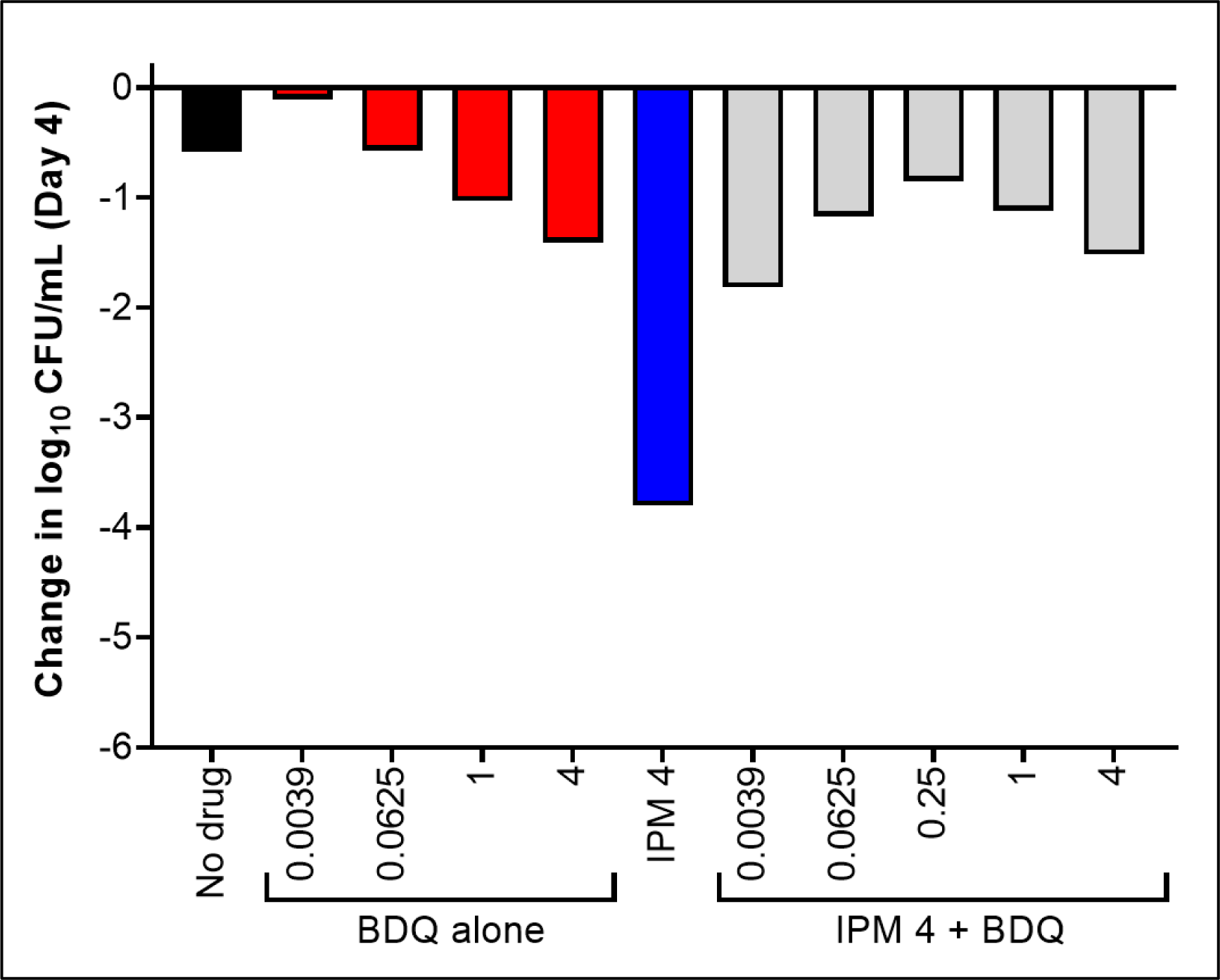
Activity of bedaquiline (BDQ) and imipenem (IPM) against nutrient-starved *M. abscessus* ATCC 19977 WT in PBS with 0.05% Tween 80. Bacteria were nutrient-starved in PBS for 20 days prior to drug exposure. The changes in log_10_ CFU/mL after 4 days of drug exposure relative to Day 0 are presented. Red bars indicate BDQ only samples; blue bars represent IPM only samples, and gray bars indicate IPM + BDQ samples. The number after the IPM abbreviation represents the concentration in µg/mL; for red and gray bars, the BDQ concentration in µg/mL is under each bar. Data after 1 day of drug exposure, as well as data from a biological replicate, are presented in **Fig. S13**. The Day 0 bacterial concentration was 6.07 log_10_ CFU/mL. CFU values are provided in **Table S11**.

### Investigation of bedaquiline and imipenem activity against intracellular *M. abscessus* ATCC 19977 WT and OM7 mutant

Our *in vitro* results highlighted the impact that experimental conditions can have on the assessment of drug activity against *M. abscessus*. In order to better understand which assay conditions, if any, may be predictive of activity in models of higher biological complexity, we next evaluated the activity of bedaquiline and imipenem against intracellular *M. abscessus* WT during infection of THP-1 monocytic cells. Exposure to bedaquiline alone at concentrations ranging from 0.25 to 16 µg/mL resulted in concentration-dependent activity, ranging from growth-limiting to bacteriostatic to bactericidal, against intracellular *M. abscessus* (**Fig. S11A-C**; see all CFU data in **Table S13**). In contrast, concentration-ranging activity was not clearly observed with imipenem at concentrations ranging from 16 to 64 µg/mL; imipenem at 16-32 µg/mL had similar activity (static or cidal, depending on the replicate) against intracellular *M. abscessus*, while imipenem at 8 µg/mL was consistently less active and only limited intracellular bacterial growth relative to the no drug control (**Fig. S11D-F**). The combination of bedaquiline at 0.5 or 1 µg/mL with imipenem at 32-64 µg/mL exerted bactericidal activity, killing ∼1-2 log_10_ intracellular CFU/mL over 5 days (**Fig. 9**). The 2-drug combinations demonstrated equivalent or better activity against intracellular *M. abscessus* compared to either drug alone, and antagonism was not observed in these assay conditions.

**Fig. 9.**
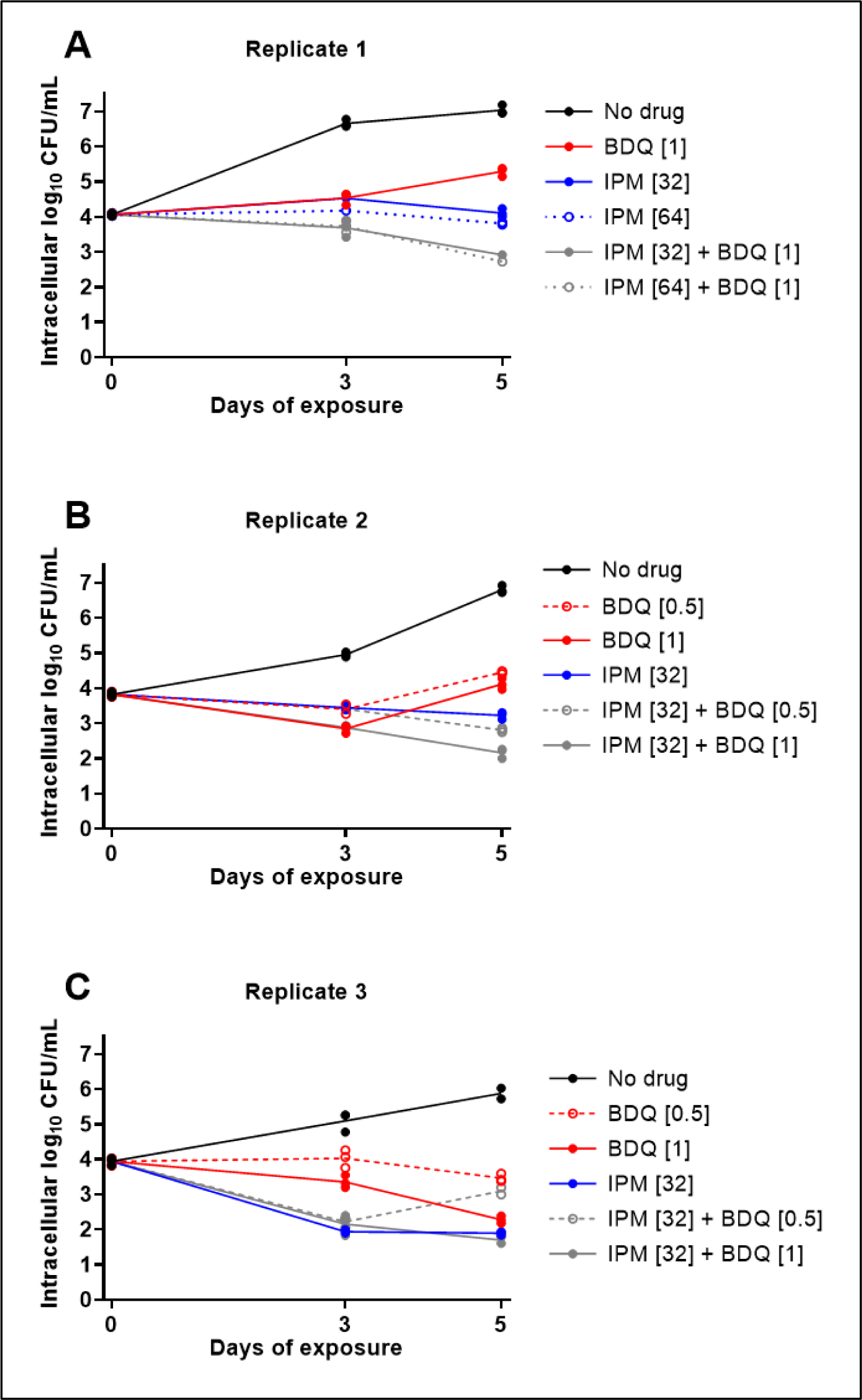
Activity of bedaquiline (BDQ) and imipenem (IPM) combinations against intracellular (THP-1 cells) *M. abscessus* ATCC 19977 WT. Data are presented for 3 biological replicates: replicate 1, panel A; replicate 2, panel B; replicate 3, panel C. Each data point represents a technical replicate, and the connecting lines pass through the mean values. The number in brackets after each drug abbreviation represents the concentration in µg/mL. All CFU values, including data from additional BDQ+IPM combinations and a fourth biological replicate, are provided in **Table S13**.

## Discussion

In this series of experiments, we systematically evaluated the *in vitro* activity of bedaquiline and imipenem against the ATCC 19977 wild type strain of *M. abscessus* and an isogenic *MAB_2299c* mutant in different culture conditions: actively growing bacteria in nutrient-rich media; net non-replicating bacteria under nutrient starvation in PBS; and actively growing intracellular bacteria in THP-1 cells. Overall, we found that the activity of bedaquiline and imipenem, either alone or in combination, was significantly influenced by assay conditions, highlighting current knowledge gaps and limiting broad generalizations with respect to translating *in vitro* data into predictions of *in vivo* efficacy.

One consistent finding was that bedaquiline alone had limited or no bactericidal activity against actively multiplying *M. abscessus* in nutrient-rich media, a finding that aligns with previous reports (6-8, 11). However, even within this limited activity, we observed that the antibacterial effects of bedaquiline were greater (on a µg/mL basis) against bacteria in CAMHB without the surfactant Tween 80 than in CAMHB supplemented with 0.05% Tween 80. We also consistently observed that bedaquiline had strong bactericidal activity against nutrient-starved *M. abscessus* in the absence of Tween 80, in agreement with our previous report (6), but the magnitude of killing decreased in the presence of Tween 80. Bedaquiline’s activity against intracellular *M. abscessus*, which was mainly bacteriostatic with limited bacterial killing, aligned best with the activity in CAMHB, with the magnitude of effect (based on the µg/mL bedaquiline added to assay media) falling somewhere between the activity observed in CAMHB with and without Tween 80.

The activity of imipenem was affected by Tween 80 to an even greater extent. In CAMHB without Tween 80, imipenem alone had limited bactericidal activity, a finding that aligns with previous reports (6, 27). In contrast, imipenem alone was highly bactericidal in CAMHB with Tween 80. In agreement with our previous study (6), imipenem alone had no bactericidal activity against nutrient-starved bacteria in the absence of Tween 80; but, in the presence of Tween 80, imipenem had striking bactericidal activity against nutrient-starved *M. abscessus*, with killing of a similar magnitude as in CAMHB with Tween 80. The largely bacteriostatic or limited bactericidal activity of imipenem against intracellular bacteria, a finding also reported by others (28, 29), aligned best with the activity in CAMHB without Tween 80.

The activity of imipenem-bedaquiline combinations was also highly dependent on assay conditions. Against actively multiplying *M. abscessus* in CAMHB with Tween 80, the potent bactericidal activity of imipenem was enhanced when bedaquiline was added at low concentrations, ≤0.0078 µg/mL. However, as the concentration of bedaquiline increased above 0.0078 µg/mL, the addition of bedaquiline decreased the bactericidal activity of imipenem until it was no longer evident, although the activity of the imipenem-bedaquiline combination was better than the activity of bedaquiline alone. Lindman and Dick previously reported that inhibitory concentrations of bedaquiline antagonize the *in vitro* bactericidal activity of imipenem against actively multiplying *M. abscessus*; however, the media used in their assay was Middlebrook 7H9 broth (BD Difco) supplemented with 0.5% albumin, 0.2% glucose, 0.085% sodium chloride, 0.0003% catalase, 0.2% glycerol, and 0.05% Tween 80 (verified by written correspondence) (8). In CAMHB without Tween 80, activity of the two-drug combination was driven by the activity of bedaquiline. Any antagonism of imipenem by bedaquiline could not be ascertained, as the activity of imipenem alone was more limited in the absence of Tween. Similarly, the bactericidal activity of imipenem-bedaquiline combinations against nutrient-starved *M. abscessus* in the absence of Tween also appeared to be driven by the activity of bedaquiline. However, in the presence of Tween 80, the addition of bedaquiline drastically antagonized the bactericidal activity of imipenem against nutrient-starved bacteria. Additive activity was observed between imipenem and bedaquiline against intracellular bacteria.

Consideration of the clinical relevance of assay conditions is always important when evaluating antibacterial drugs and regimens *in vitro*. Unfortunately, for treatment of *M. abscessus* lung infections, there are no established or validated *in vitro* assays known to predict *in vivo* activity, and the situation is further complicated by the lack of data regarding clinical treatment outcomes. However, inclusion of Tween 80 in drug activity assays may lead to conclusions that are clinically misleading. Tween is a non-ionic surfactant used as an emulsifier in drugs and food. It has been known for decades that Tween 80 can impact drug activity against bacteria, and for agents acting on the cell wall, such as imipenem, Tween 80 may increase susceptibility by increasing membrane permeability (25, 26). Interestingly, Tween 80 had the opposite impact on bedaquiline activity against *M. abscessus*, decreasing bedaquiline’s activity in CAMHB and in nutrient starvation assays. Lounis *et al.* reported similar findings for bedaquiline against *M. tuberculosis*, and suggested that Tween 80 may be directly interacting with bedaquiline, thus limiting the amount of free drug in solution (30). We initially included Tween 80 in our assay to reduce bacterial clumping, without appreciating how drastic its impact would be on assay outcomes. In this study, the most biologically complex system used was the intracellular infection assay. Overall, the drug activity against intracellular bacteria seemed to be best predicted by activity in CAMHB without Tween 80, providing further weight to the data from *in vitro* assays performed without Tween 80.

The impact of Tween 80 on nutrient-starved drug activity assays is further complicated by the presence of oleic acid, a breakdown product of Tween 80 in aqueous solutions and hydrolysis by mycobacterial enzymes (17, 31-33). Oleic acid can serve as a nutrient source for mycobacteria. Due to its known ability to enhance the *in vitro* growth of *M. tuberculosis* (17, 31), it is routinely used as a supplement in liquid and solid media for isolating and cultivating mycobacteria. Including Tween 80 in the nutrient-starvation samples *during* drug exposure likely introduced the nutrient oleic acid into the PBS. Although we did not observe net bacterial growth in these conditions, it is possible that the presence of oleic acid impacted bacterial metabolism such that the bacteria became more susceptible to killing by imipenem. This concept is further supported by data from Berube *et al.*, who reported that imipenem alone had no bactericidal activity against nutrient-starved *M. abscessus* in PBS with 0.05% tyloxapol, a surfactant that does not provide a nutrient source for the bacteria (34). In this study, we did not have an intracellular or other more biologically complex model of net non-replicating *M. abscessus* to aid in our translation of drug activity in the presence or absence of Tween 80.

The *in vivo* activity of imipenem, bedaquiline, and/or imipenem-bedaquiline have been evaluated in different mouse models of *M. abscessus* lung infection. Story-Roller *et al.* reported that imipenem (100 mg/kg twice daily) had modest *in vivo* bactericidal activity against an actively multiplying *M. abscessus* lung infection in a C3HeB/FeJ mouse model with dexamethasone-induced immunosuppression, with imipenem killing just over 2 log_10_ CFU/lung over 4 weeks of treatment (35). Le Moigne *et al.* also evaluated imipenem activity in the lungs of *M. abscessus* -infected C3HeB/FeJ mice, but without inducing immune suppression, and thus in their model, the bacterial burden in untreated mice decreased over time. The lung bacterial burden in mice that received 2 weeks of imipenem alone (100 mg/kg twice daily) was no different from the lung burden in untreated mice (9). Le Moigne *et al.* also evaluated the activity of bedaquiline and bedaquiline-imipenem combinations in this mouse model. After 2 weeks of treatment, bedaquiline alone (30 mg/kg once daily) contributed modest bactericidal activity resulting in an approximately 1 log_10_ CFU/lung reduction compared to the decline observed in untreated mice, and the activity of imipenem-bedaquiline was like that of bedaquiline alone, indicating that the activity of the 2-drug combination was driven more by bedaquiline’s activity than imipenem’s activity (9). Lerat *et al.* also evaluated bedaquiline alone (25 mg/kg) in an athymic nude mouse model of *M. abscessus* infection in which the lung bacterial burden in untreated mice decreased over time. After 1 month of treatment, there was no difference in lung bacterial burden between untreated and bedaquiline-treated mice but, after 2 months of treatment, there was a modest decline in lung CFU counts in bedaquiline-treated mice (36). In contrast, Obregón-Henao *et al.* reported that dosing bedaquiline alone (30 mg/kg once daily for 9 days) resulted in strong bactericidal activity against *M. abscessus* in the lungs of GKO^-/-^ mice, in which the bacterial counts decreased in untreated mice, and also in the lungs of SCID mice, in which the lung bacterial burden increased in untreated mice (37). The mouse models utilized by Story-Roller *et al*., Le Moigne *et al.*, and Lerat *et al.* used *M. abscessus* strain ATCC 19977, while Obregón-Henao *et al.* used a different strain in their studies.

How well do the imipenem and bedaquiline activity profiles from our different *in vitro* and intracellular assays align with the observed *in vivo* activity in these mouse models of *M. abscessus* lung infection? Direct comparison between the *in vitro* and *in vivo* studies is confounded by several factors. The duration of our *in vitro* studies, which ranged from 3-7 days, was much shorter than most of the treatment durations in mouse models. In addition, understanding the drug exposures in mice is more complicated than in *in vitro* assays. The imipenem dose administered to mice in the studies described, 100 mg/kg, has been shown to result a maximum plasma concentration of about 85 µg/mL, but the drug is rapidly metabolized, with a plasma half-life of about 18 minutes, which leads to the requirement for multiple doses per day to achieve suitable exposures (38). Further, the imipenem concentrations that we utilized for our *in vitro* combination studies do not fluctuate over time in the manner that they do in mice. Overall, if the goal is to have the range of concentrations tested be representative of exposures in mice, and the imipenem activity data in the mouse models of actively multiplying and declining *M. abscessus* lung infections would have been best predicted by the *in vitro* activity observed in assays without Tween 80 and by the intracellular assays.

For bedaquiline, comparison between *in vitro* and *in vivo* findings is limited by the nature of bedaquiline metabolism in mice. Compared to humans, mice more rapidly convert bedaquiline into its M2 metabolite, *N*-desmethyl bedaquiline, which is the dominant species in mouse plasma and lungs (39). Without understanding the activity of the M2 metabolite against *M. abscessus*, it is impossible to compare *in vitro* activity to the activity observed in bedaquiline-treated mice. However, as bedaquiline treatment resulted in bactericidal activity in several mouse models (9, 37), it is reasonable to hypothesize that the M2 metabolite does indeed have activity against *M. abscessus.* It is known that the M2 metabolite contributes approximately 50% of the *in vivo* activity of bedaquiline in mice in infected with *M. tuberculosis* (39). Keeping all of this in mind, and assuming average plasma concentrations in mice of around 0.2 and 1 µg/mL of BDQ and M2, respectively (39), the nature of the bedaquiline and imipenem-bedaquiline activity in the mouse model reported by Le Moigne *et al.* seems to align best with *in vitro* activity observed in the nutrient starvation assay without Tween 80.

Another aspect of the present study was the assessment of bacterial ATP levels in association with imipenem and bedaquiline exposure and bacterial cell death. Similar to Lindman and Dick (8), we found that exposure to bactericidal concentrations of imipenem in CAMHB with Tween 80 was associated with an increase in bacterial ATP levels. This relationship also correlated with the activity of imipenem-bedaquiline combinations in these assay conditions. As observed by Lindman and Dick, exposure to inhibitory concentrations of bedaquiline alone was associated with a modest decline in ATP levels, consistent with bedaquiline’s mechanism of action, and ATP levels decreased in association with the observed antagonism of imipenem’s bactericidal effects when these inhibitory concentrations of bedaquiline were added to imipenem. However, we also expanded on the observations of Lindman and Dick by evaluating the addition of lower, subinhibitory concentrations of bedaquiline to imipenem, which proved to increase, rather than antagonize, the bactericidal effects of imipenem in association with increases in ATP levels. These concentration-dependent effects of bedaquiline when combined with imipenem may shed further light on the relationship between intrabacterial ATP levels and killing by imipenem and other β-lactams described by Lindman and Dick. That the bacteria may respond to sub-inhibitory concentrations of bedaquiline by, at least transiently, increasing ATP production and that this may augment a similar response to imipenem exposure and result in additive effects on both ATP levels and killing further supports their conclusion that increased ATP levels are at least a surrogate for bactericidal effects of imipenem and may be part of the causal pathway. Interestingly, in CAMHB without Tween 80, a condition in which imipenem was not as bactericidal, bacterial ATP levels in imipenem-exposed *M. abscessus* were not higher than those in the no drug control samples (**Figs. S4**, **S5**). Just as the bactericidal activity of imipenem-bedaquiline combinations seemed to be driven by bedaquiline in this condition, ATP levels in bacteria exposed to imipenem-bedaquiline decreased as the bedaquiline concentration increased, and overall the ATP levels in bacteria exposed to imipenem-bedaquiline were not different than in bacteria exposed to bedaquiline alone. A similar relationship was observed during drug exposure in nutrient starvation conditions without Tween 80. Because bedaquiline was bactericidal in these conditions, when adjusting the RLU by CFU counts, it may appear that ATP levels are increasing as the bedaquiline concentration increases (**Fig. 7**). However, the non-CFU-adjusted RLU data indicate that total ATP levels decreased with increasing BDQ concentration (**Figs. S7**, **S8**).

Again, we must ask, what is the biological and/or clinical relevance of these data? Overall, we lean towards the conclusion that assay conditions without Tween 80 are more relevant to drug activity in intracellular and mouse infection models. Lindman and Dick reported that the bactericidal activity of imipenem and cefoxitin against actively multiplying *M. abscessus* was associated with bacterial ATP bursts, and that the addition of bedaquiline antagonized this activity; however, the assay media used was not clearly stated (8). Shetty and Dick demonstrated similar bacterial ATP bursts in *Mycobacterium bovis* BCG exposed to cell wall synthesis inhibitors, including isoniazid, and that both bactericidal activity and ATP levels were decreased when the samples were co-exposed to bedaquiline or other agents affecting ATP production (21); and Zeng *et al.* confirmed these findings (20). These assays were conducted in Middlebrook 7H9 broth supplemented with 0.05% Tween 80 and DTA medium supplemented with 0.02% Tween 80. There are intriguing biological processes occurring in mycobacterial cells exposed to cell wall synthesis inhibitors and agents that target ATP production, and further dissection of the impact of media conditions on these relationships is needed to understand their impact during *in vivo* treatment.

The overarching purpose of our work is to provide data which will help inform the design of improved treatment regimens for patients with *M. abscessus* lung disease. A key issue we sought to address in this work was whether imipenem and bedaquiline could be used together in a treatment regimen. The data from our *in vitro* (without Tween) and intracellular assays, as well as from the *in vivo* combination study reported by Le Moigne *et al.* (9), indicate that the activity of imipenem-bedaquiline combinations may be driven largely by the activity of bedaquiline, without marked antagonism of imipenem. However, even if antagonism of imipenem by bedaquiline does have clinical relevance, it may not necessarily rule out using these drugs together. For example, in tuberculosis treatment, the activity of isoniazid, which is highly bactericidal against actively multiplying *M. tuberculosis*, is antagonized both in mouse models and in humans, by pyrazinamide, a drug more active against non-replicating bacteria; however, regimens including this combination of drugs have been used successfully for decades (40-43). Furthermore, if we consider that imipenem has a much shorter half-life than bedaquiline and reaches steady state much sooner than bedaquiline, imipenem could exert its activity against actively replicating bacilli without interference or may be augmented by sub-inhibitory concentrations of bedaquiline during the initial stages of treatment. Finally, our results show that mutational disruption of *MAB 2299c* reduces bacterial susceptibility to bedaquiline across multiple assay conditions but may increase susceptibility to imipenem. If bedaquiline proves to be a drug with treatment-shortening potential in *M. abscessus* infections, imipenem may be a particularly effective companion agent for preventing emergence of bedaquiline resistance. Ultimately, an imipenem-bedaquiline combination would be part of a larger multi-drug regimen, and other optimized companion agents may further enhance the utility of imipenem-bedaquiline for treatment of *M. abscessus* lung disease.

## Materials and Methods

### Bacteria

*M. abscessus* subsp. *abscessus* strain ATCC 19977 was obtained from the American Type Culture Collection (ATCC) and used in all experiments.

### Media

Bacterial cultures were initiated in standard growth media: Middlebrook 7H9 broth supplemented with 10% (v/v) Middlebrook OADC supplement, 0.1% (v/v) glycerol, and 0.05% (v/v) Tween 80. Drug activity assays in nutrient-rich media were performed using either Middlebrook 7H9 broth with 10% (v/v) OADC and 0.1% (v/v) glycerol but without Tween; or CAMHB with or without 0.05% Tween 80. Polystyrene petri-dishes (100 mm × 15 mm) containing 20 mL 7H11 agar supplemented with 10% (v/v) OADC and 0.1% (v/v) glycerol were used to determine CFU counts. Difco BBL Mueller Hinton II broth (cation-adjusted) powder (i.e., CAMHB powder), Difco Middlebrook 7H9 broth powder, Difco Mycobacteria 7H11 agar powder, and BBL Middlebrook OADC enrichment were manufactured by Becton, Dickinson and Company. Glycerol and Tween 80 were purchased from Fisher Scientific.

### Drugs

Imipenem powder was purchased from Biosynth Carbosynth. Bedaquiline and N-4S-methylcyclohexyl-4,6-dimethyl-1H-indole-2-carboxamide were provided by the TB Alliance. Clofazimine, meropenem and clarithromycin were purchased from Sigma. Drugs were dissolved in either PBS (imipenem and meropenem) or dimethyl sulfoxide (bedaquiline, N-4S-methylcyclohexyl-4,6-dimethyl-1H-indole-2-carboxamide, clofazimine, and clarithromycin), and drug solutions were filter-sterilized.

### Selection and characterization of *MAB_2299c* mutants

Frozen stock of *M. abscessus* ATCC 19977 WT was cultured in standard growth media until an optical density at 600 nm (OD_600_) >1.5 was achieved. Ten-fold dilutions (in PBS) of the bacterial suspension were cultured on 7H11 agar containing the following concentrations of clofazimine: 8µg/mL, 16µg/mL, 32µg/mL, and 64µg/mL. Individual colonies growing on 7H11 agar containing 8 µg/mL clofazimine were selected, expanded, and stored in growth media at -80°C; these isolates were also subjected to a second round of selection on agar containing clofazimine at 32µg/mL and 64µg/mL. Individual colonies were selected, expanded, and stored in growth media. The MICs of both clofazimine and bedaquiline were determined for selected isolates using the broth microdilution method in round-bottom, polystyrene 96-well plates as previously described (44), with assays conducted in CAMHB and Middlebrook 7H9 with 10% (v/v) OADC and 0.1% (v/v) glycerol, without Tween 80 in either media. The concentration ranges tested were clofazimine 128-0.125 µg/mL and bedaquiline 4-0.0038 µg/mL in CAMHB and 16-0.125 µg/mL in 7H9. MIC was defined as the lowest concentration without visible growth. Isolates with increased MICs at least 4 times higher than the WT parent strain were evaluated by PCR and sequencing for mutations in the following genes: *MAB_2299c*, *MAB_4384*, and *atpE* (*MAB_1448*). Genomic DNA was prepared from each isolate as previously described (45). Primer sequences and expected amplicon sizes were as follows: *MAB_2299c* forward: 5’ CGC GTT TCA TCA GGA TCT TT 3’, reverse: 5’ CCT ACG TGG ATG CCA AGG 3’, 862bp; *MAB-_4384* forward: 5’ GGC AGG GTC AGC AGA AAT 3’, reverse: 5’ ATG TTG TGT GCG GGG TCT 3’, 840 bp; *atpE* forward: 5’ TGG ACG AGG ACC ATC ACT AA 3’, reverse: 5’ GAC GGC AGA AGC GAC AC 3’, 375 bp. Purified amplicons were analyzed by Sanger sequencing at GENEWIZ. Sequencing results were compared against the *M. abscessus* ATCC 19977 published genome, NCBI accession number NC_010397.

### Drug activity assays in nutrient-rich media

Frozen stocks of *M. abscessus* were cultured in standard growth media until the bacterial suspension reached OD_600_ ∼1 (approximately 10^7^-10^8^ CFU/mL), at which point the assays were initiated (“Day 0”). Assays were conducted with the indicated media in a total volume of 2.5 mL in 14 mL, round-bottom, polystyrene tubes as previously described (6).

### Drug activity assays in nutrient starvation conditions

Frozen stocks of *M. abscessus* were cultured in growth media from frozen stock until the bacterial suspension reached OD_600_ ∼1. The bacterial suspension was subsequently spun (1900 rcf for 10 minutes), washed, and resuspended three times to the original volume in PBS with 0.05% (v/v) Tween 80, as previously described(6). After the third wash, the bacteria were resuspended in PBS with 0.05% (v/v) Tween 80 and were incubated at 37°C for the indicated duration (14 or 20 days). Assays were initiated after the indicated duration of nutrient starvation; the bacterial suspensions were diluted in PBS with or without Tween 80 to an OD_600_ of 0.6 on Day 0 of the assay. Drugs stocks were added to achieve the indicated concentrations. Assays were conducted in a total volume of 2.5 mL in 14 mL, round-bottom, polystyrene tubes as previously described (6).

### Relative ATP measurements

The Promega BacTiter-Glo™ Microbial Cell Viability Assay was used to measure the relative ATP in assay samples. At each indicated timepoint, 20 µL of assay sample was removed and incubated with 20µL of the BacTiter-Glo™ reagent in clear 1.5 mL snap-cap tubes at room temperature for 5 minutes. After the incubation, RLU was recorded for each sample using Turner TD-20/20 luminometer.

### Intracellular infections and drug activity assays

Frozen stocks of *M. abscessus* were cultured in standard growth media. After reaching OD_600_ ∼1, the culture was spun at 1100 rpm for 7 minutes to remove any cellular debris or dead cells. For infection, the bacterial suspension was subsequently diluted to reach an OD_600_ of 0.0005 in Roswell Park Memorial Institute (RPMI) medium with L-glutamine, supplemented with 10% (v/v) heat-inactivated fetal bovine serum (FBS) to generate a bacterial suspension of ∼5x10^4^ cells/mL. THP-1 cells (obtained from ATCC, product number TIB-202) were grown in supplemented RPMI medium to a density of 10^5^ cells/mL, aliquoted into 24-well polystyrene plates (tissue culture-treated, 1mL cell suspension per well), and activated by incubation with 50nM phorbol 12-myristate 13-acetate for 24 hours at 37°C. After activation, the adherent THP-1 cells were washed with PBS and then infected with 1mL of the adjusted bacterial suspension at OD_600_ 0.05, 0.005, 0.0005 to achieve an MOI of 10:1, 1:1, and 1:10 (bacteria:THP-1 cells), respectively. All drug activity assays were conducted at an MOI of 1:10. After 3 hours of infection at 37°C, each well was washed three times with 1mL PBS to remove extracellular bacteria. After the final wash, 1 mL RPMI media containing bedaquiline or imipenem at the indicated concentrations (or no drug) was added to the appropriate wells. Drug-containing media was replaced daily. At each indicated timepoint, each well was washed three times with PBS, and then the THP-1 cells were lysed with sterile deionized water, and lysates were collected for CFU determination. In each biological replicate, three wells were used for each condition (*i.e.*, triplicate technical replicates). All THP-1 cells were incubated at 37°C and 5% CO_2_.

### Quantitative cultures, CFU counting, and analysis

In all assays, CFU counts were determined by culturing serial 10-fold dilutions of bacterial suspensions on 7H11 agar. CFU counts were not determined for samples where the bacteria had overgrown, clumped, and fallen out of suspension. Serial dilutions were prepared in PBS, and 0.5 mL of each dilution was cultured on 7H11 agar. After all liquid was absorbed into the agar, the plates were sealed in plastic bags and incubated at 37°C for 5-7 days before reading and recording CFU counts. The dilution that yielded CFU counts between 10-120 and closest to 50 was used to determine CFU/mL. The CFU/mL value (*x*) was log transformed as log_10_ (*x* + 1) prior to analysis. All analyses were conducted using GraphPad Prism version 9.

## Acknowledgements.

This work was funded by the National Institutes of Health (R21-AI137814 to ELN), the Cystic Fibrosis Foundation (ELN), and by Genentech (research fellowship grant to OM). KED is supported by National Institutes of Health (K24AI150349).

## SUPPLEMENTAL MATERIAL

**Fig. S1.**
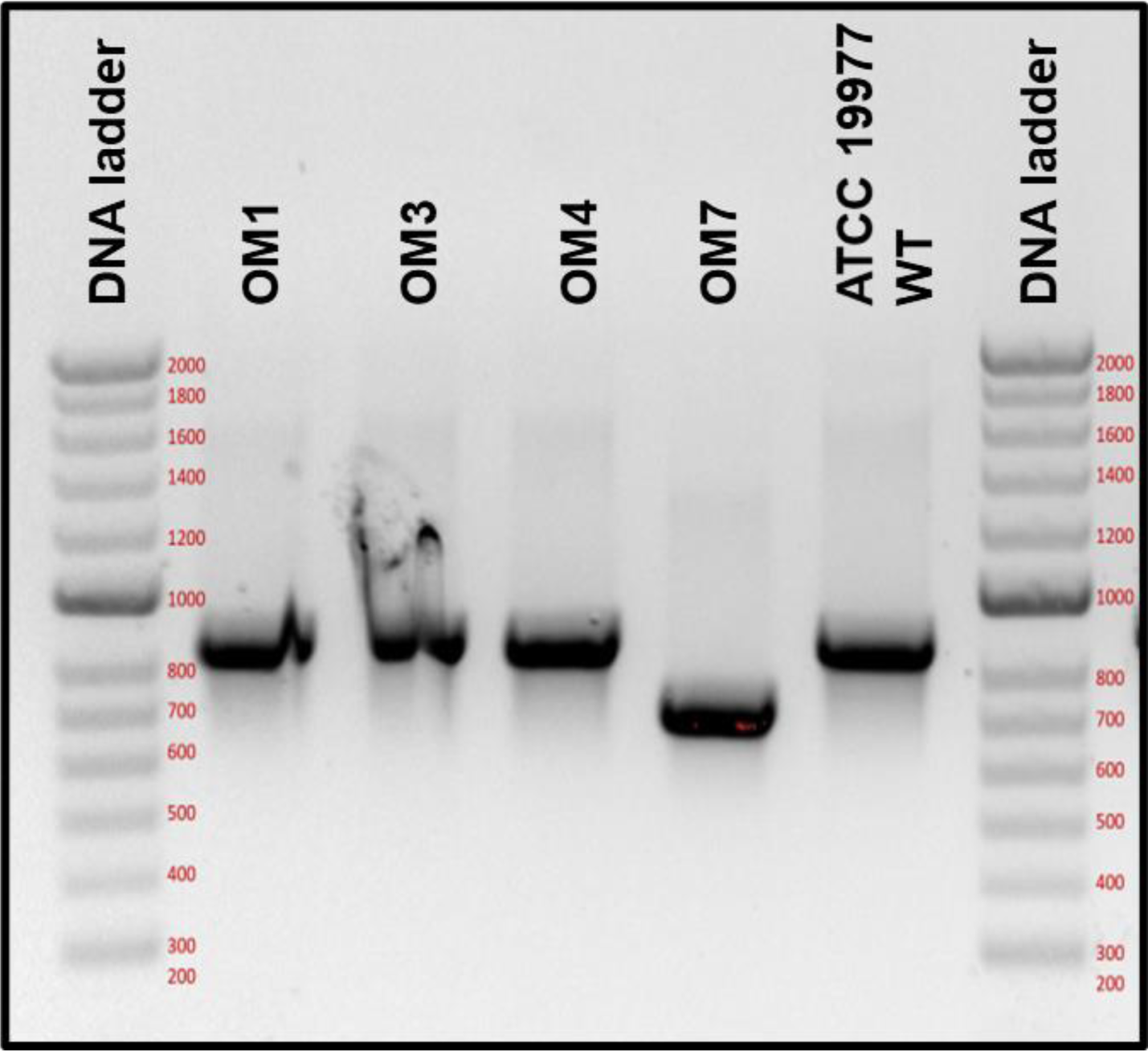
*MAB_2299c* PCR amplicons from *M. abscessus* isolates with reduced susceptibility to clofazimine and bedaquiline. The expected amplicon size for the *M. abscessus* ATCC 19977 WT parent strain was 796 bp.

**Fig. S2.**
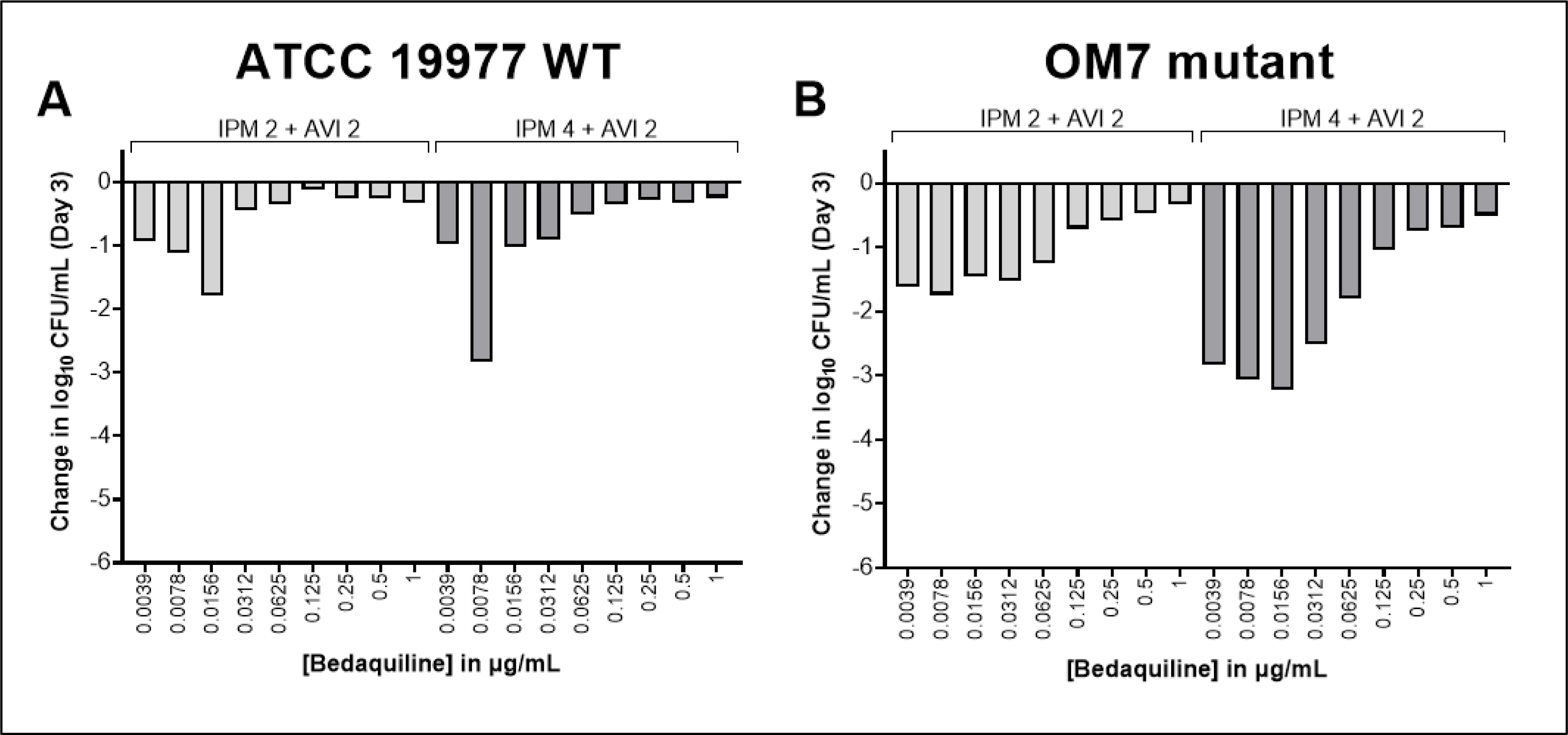
Concentration-ranging bactericidal activity of bedaquiline with imipenem (IPM)/avibactam (AVI) combinations against *M. abscessus* ATCC 19977 WT parent (A) and OM7 mutant (B) strains. The change in log10 CFU/mL after 3 days of drug exposure relative to Day 0 is presented for each drug/strain set. Assay medium was CAMHB with 0.05% Tween 80. The number after each IPM and AVI abbreviation represents the concentration µg/mL. The Day 0 bacterial concentrations were 6.13 and 6.22 log10 CFU/mL for WT and OM7 mutant strains, respectively. CFU values are provided in **Table S4**.

**Fig. S3.**
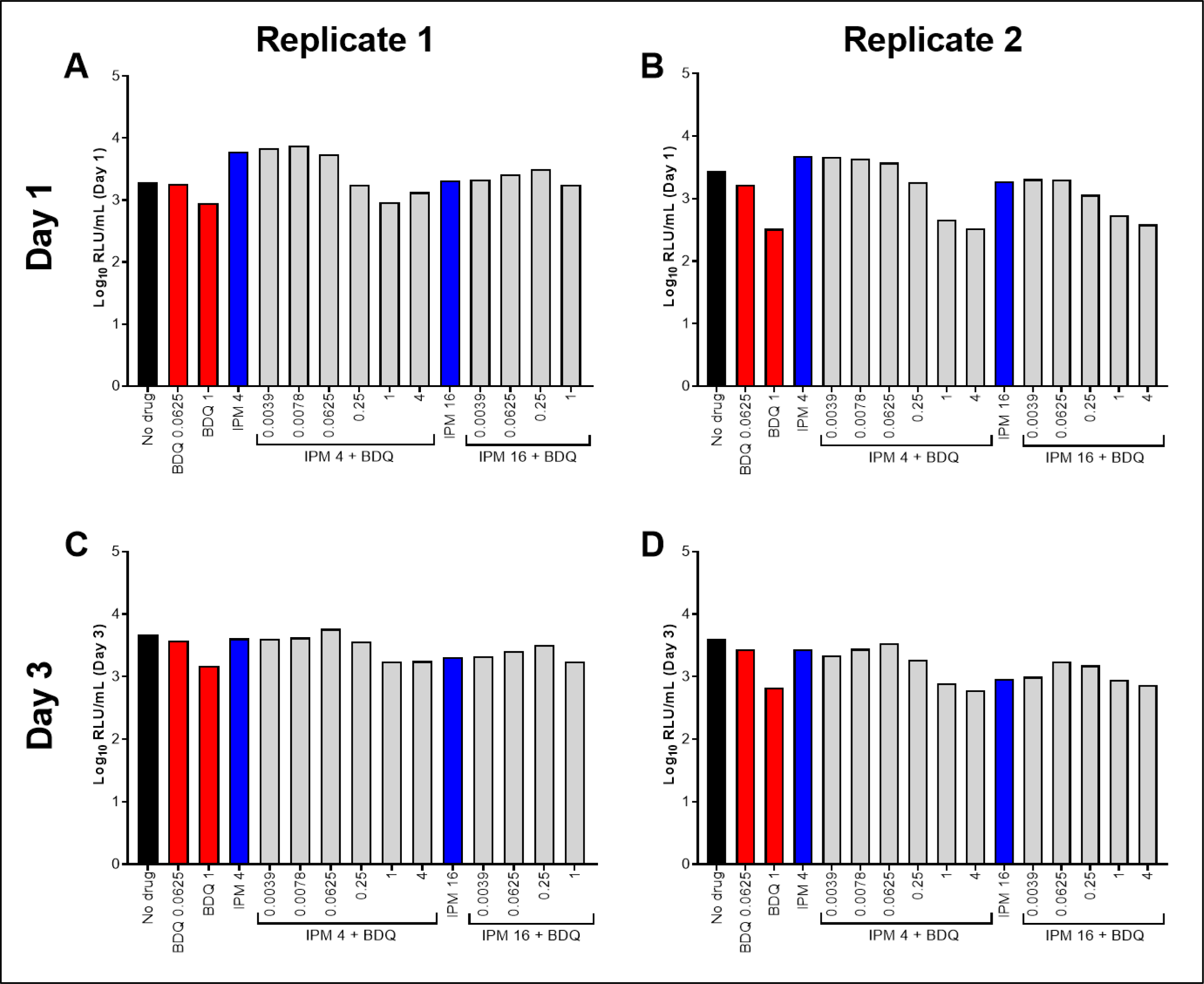
Relative ATP levels, not adjusted by CFU counts, associated with exposure of *M. abscessus* ATCC 19977 WT to bedaquiline (BDQ) and imipenem (IPM) for 1 day (A,B) or 3 days (C,D) in CAMHB with 0.05% Tween 80. Replicate 1 (A,C) is the experiment presented in Fig. 4A,**C**. Replicate 2 (B,D) is the experiment presented in Fig. 4B**,D**. ATP levels are indicated by relative light units (RLU)/CFU from samples. Black bars represent the no drug control; red bars indicate BDQ only samples; blue bars represent IPM only samples, and gray bars indicate IPM plus BDQ samples. The number after each drug abbreviation represents the concentration in µg/mL (for gray bars the BDQ concentration in µg/mL is presented under each bar). RLU values are provided in **Table S6**.

**Fig. S4.**
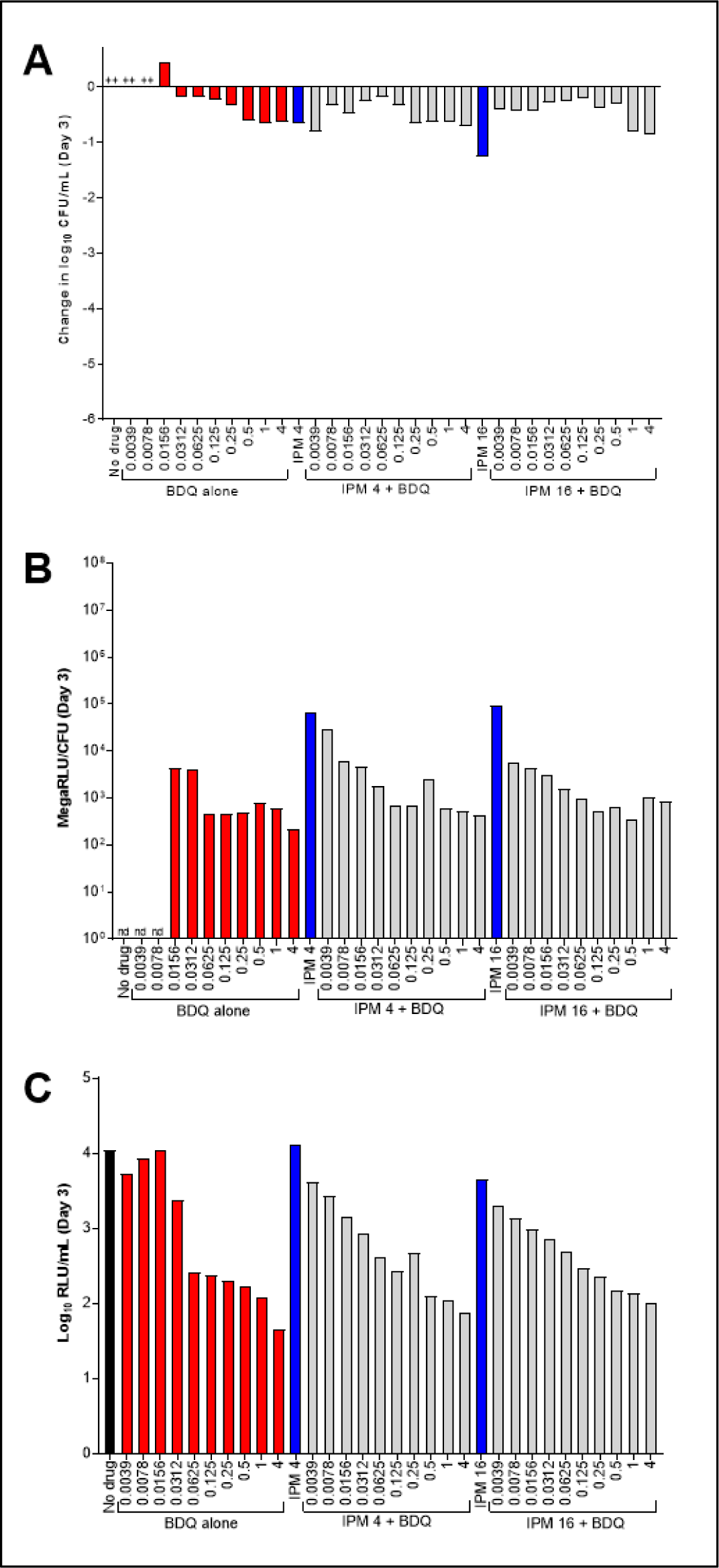
Activity of bedaquiline (BDQ) and imipenem (IPM) against *M. abscessus* ATCC 19977 WT (A), with CFU-adjusted (B) and unadjusted (C) relative bacterial ATP levels, in CAMHB without Tween 80. These data are from a second biological replicate of the experiment in Fig. 6. The changes in log10 CFU/mL after 3 days of drug exposure relative to Day 0 are presented in panel A. CFU-adjusted ATP levels, indicated by relative light units (RLU)/CFU from samples after 3 days of drug exposure, are presented in panel B; relative ATP levels, not adjusted by CFU counts, are presented in panel C. Black bars indicate the no drug control; red bars indicate BDQ only samples; blue bars represent IPM only samples, and gray bars indicate IPM + BDQ samples. The number after the IPM abbreviation represents the concentration in µg/mL; for red and gray bars, the BDQ concentration in µg/mL is under each bar. The Day 0 bacterial concentration for panel A was 5.95 log10 CFU/mL. CFU and RLU values are provided in **Table S9**.

**Fig. S5.**
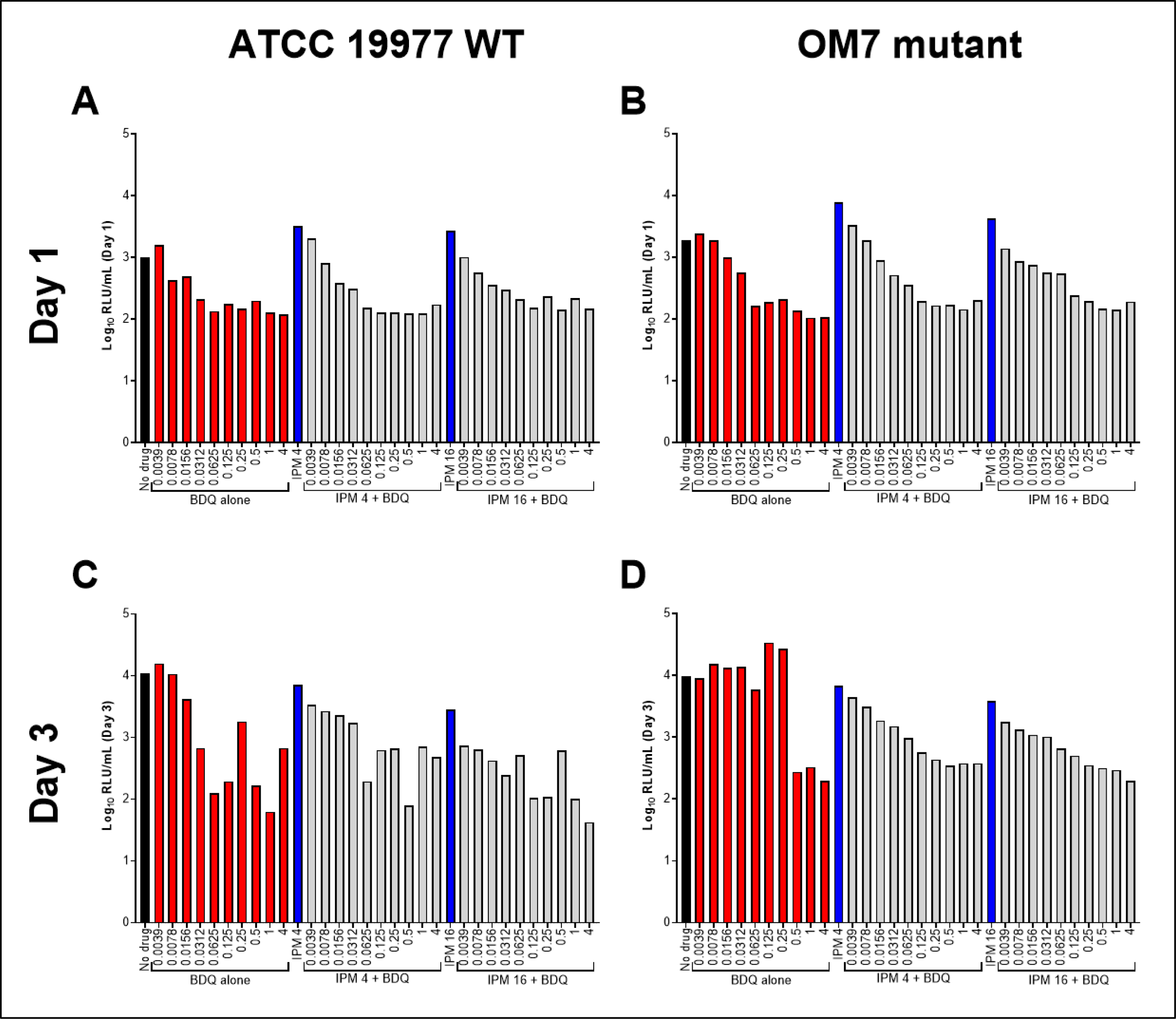
Relative ATP levels, not adjusted by CFU counts, associated with exposure of *M. abscessus* ATCC 19977 WT (A,C) and OM7 mutant (B,D) to bedaquiline (BDQ) and imipenem (IPM) for 1 day (A,B) or 3 days (C,D) in CAMHB without Tween 80. ATP levels are indicated by relative light units (RLU)/CFU from samples. Black bars represent the no drug control; red bars indicate BDQ only samples; blue bars represent IPM only samples, and gray bars indicate IPM plus BDQ samples. The number after each drug abbreviation represents the concentration in µg/mL (for gray bars the BDQ concentration in µg/mL is presented under each bar). RLU values are provided in **Table S8**.

**Fig. S6.**
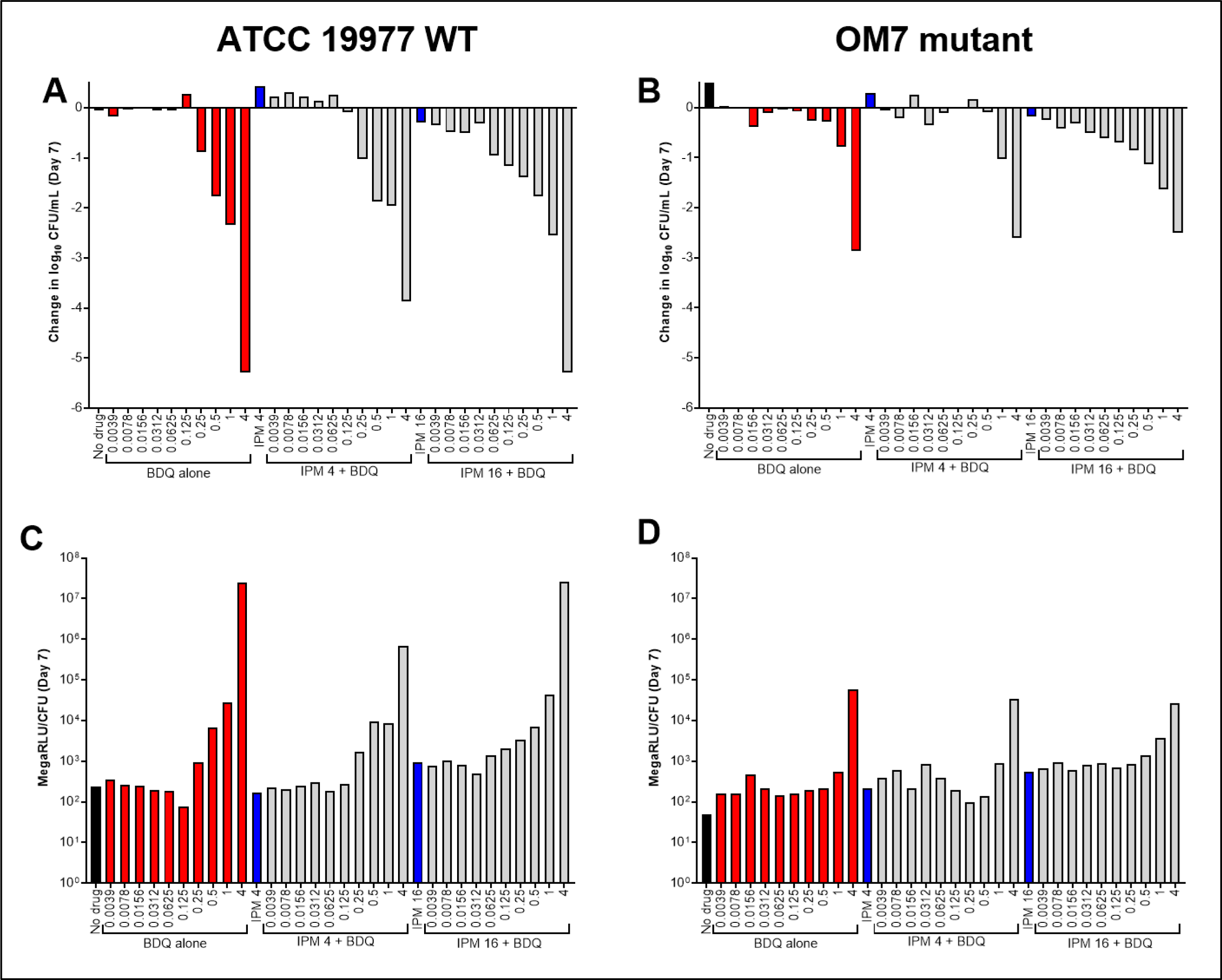
Activity of bedaquiline (BDQ) and imipenem (IPM) against nutrient-starved *M. abscessus* ATCC 19977 WT (A) and OM7 mutant (B), with relative bacterial ATP levels (C,D), in PBS without Tween 80. Bacteria were nutrient-starved in PBS for 14 days prior to drug exposure. The changes in log10 CFU/mL after 7 days of drug exposure relative to Day 0 are presented in panels A and B; CFU-adjusted ATP levels, indicated by relative fluorescence units (RLU)/CFU from samples, after 7 days of drug exposure are presented in panels C and D. Black bars indicate the no drug control; red bars indicate BDQ only samples; blue bars represent IPM only samples, and gray bars indicate IPM + BDQ samples. The number after the IPM abbreviation represents the concentration in µg/mL; for red and gray bars, the BDQ concentration in µg/mL is under each bar. Data after 3 days of drug exposure are presented in Fig. 7. RLU/mL (not adjusted for CFUs) for WT and OM7 are presented in **Fig. S7** and **Fig. S8**, respectively. The Day 0 bacterial concentration for panels A and B was 6.25 and 6.29 log10 CFU/mL, respectively. CFU and RLU values are provided in **Table S10**.

**Fig. S7.**
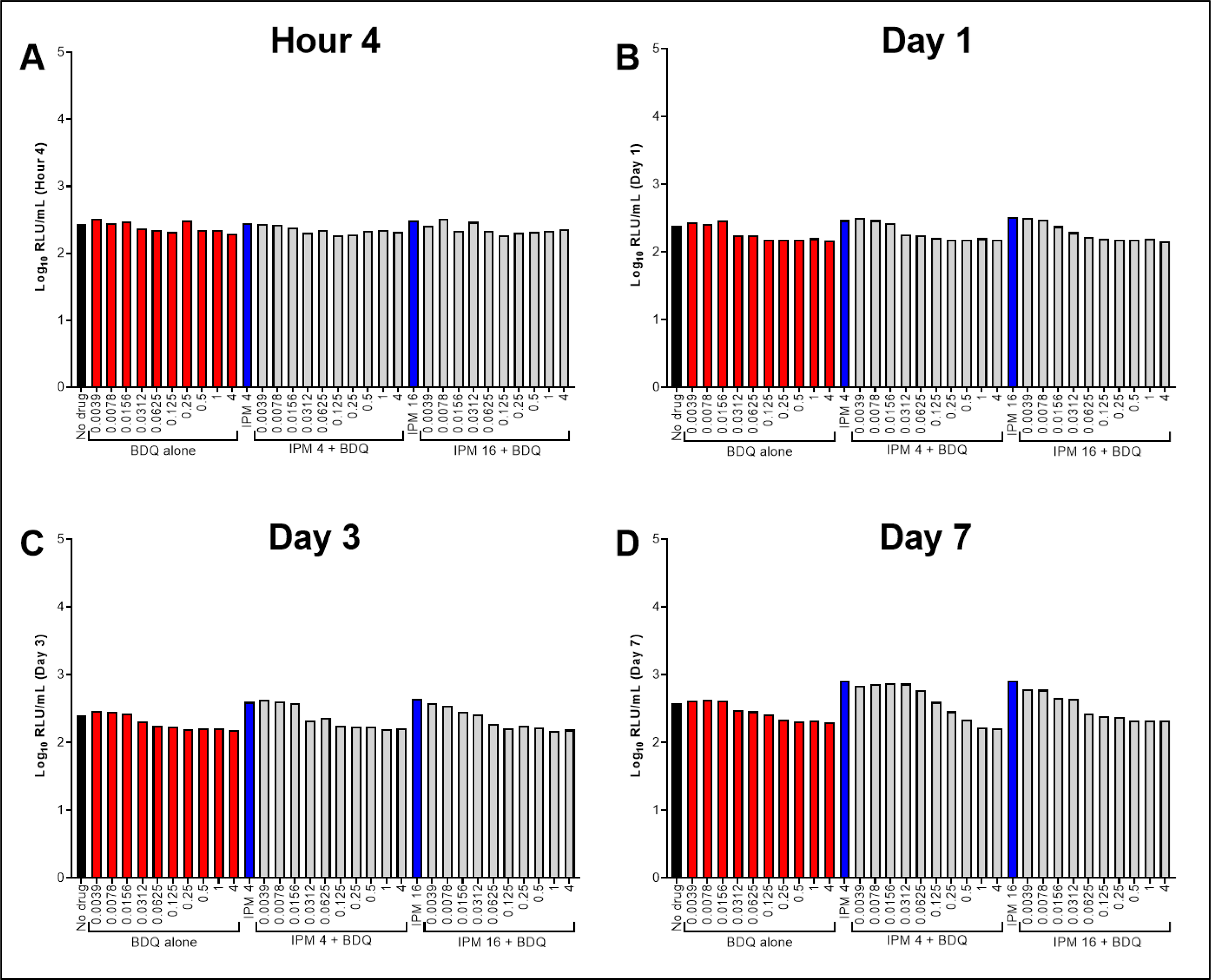
Relative ATP levels, not adjusted by CFU counts, associated with exposure of nutrient-starved *M. abscessus* ATCC 19977 WT to bedaquiline (BDQ) and imipenem (IPM) for 4 hours (A), 1 day (B), 3 days (C), or 7 days (D) in PBS without Tween 80. Bacteria were nutrient-starved in PBS for 14 days prior to drug exposure. ATP levels are indicated by relative light units (RLU)/CFU from samples. Black bars represent the no drug control; red bars indicate BDQ only samples; blue bars represent IPM only samples, and gray bars indicate IPM plus BDQ samples. The number after each drug abbreviation represents the concentration in µg/mL (for gray bars the BDQ concentration in µg/mL is presented under each bar). RLU values are provided in **Table S10**.

**Fig. S8.**
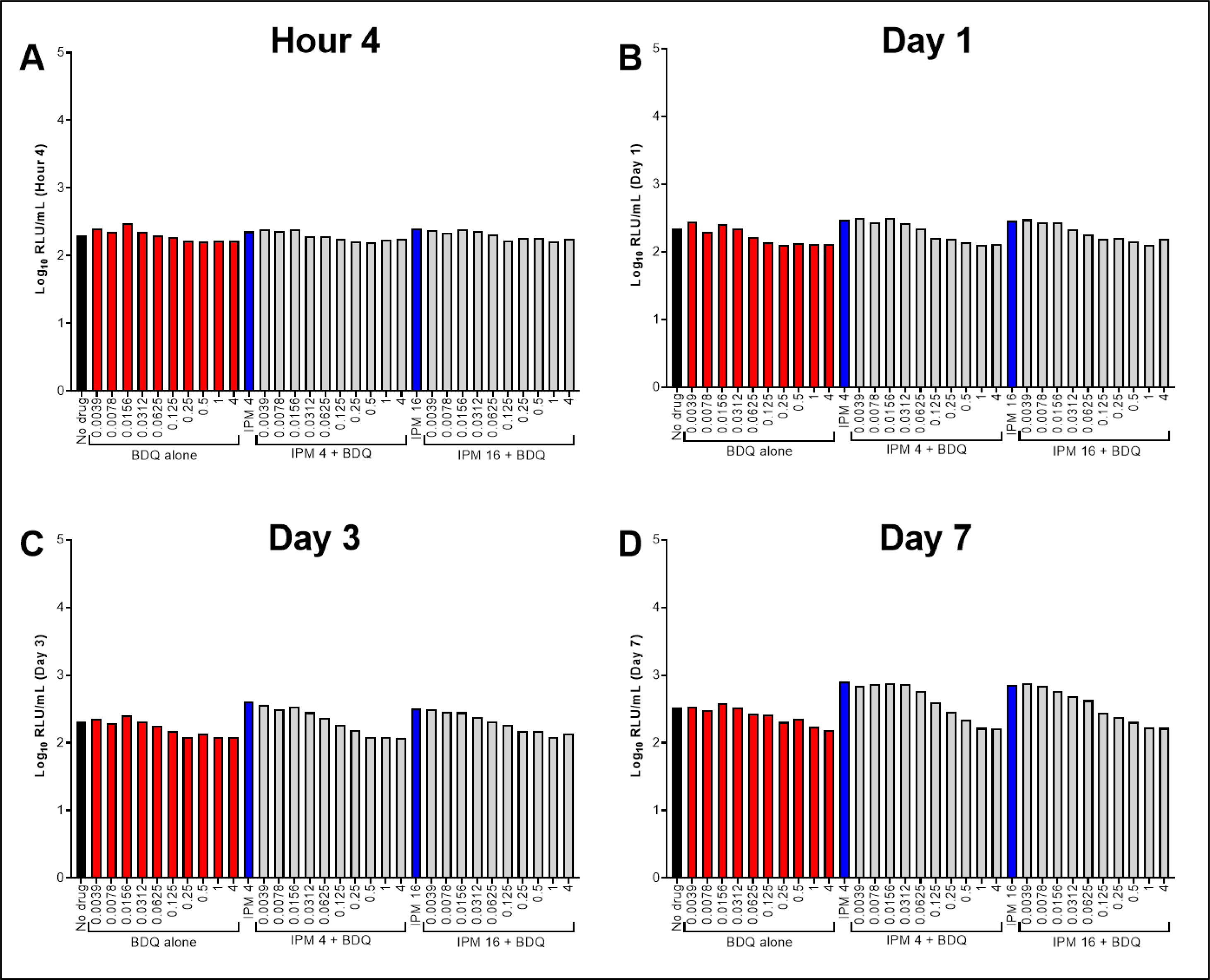
Relative ATP levels, not adjusted by CFU counts, associated with exposure of nutrient-starved *M. abscessus* OM7 mutant to bedaquiline (BDQ) and imipenem (IPM) for 4 hours (A), 1 day (B), 3 days (C), or 7 days (D) in PBS without Tween 80. Bacteria were nutrient-starved in PBS for 14 days prior to drug exposure. ATP levels are indicated by relative fluorescence units (RLU)/CFU from samples. Black bars represent the no drug control; red bars indicate BDQ only samples; blue bars represent IPM only samples, and gray bars indicate IPM plus BDQ samples. The number after each drug abbreviation represents the concentration in µg/mL (for gray bars the BDQ concentration in µg/mL is presented under each bar). RLU values are provided in **Table S10**.

**Fig. S9.**
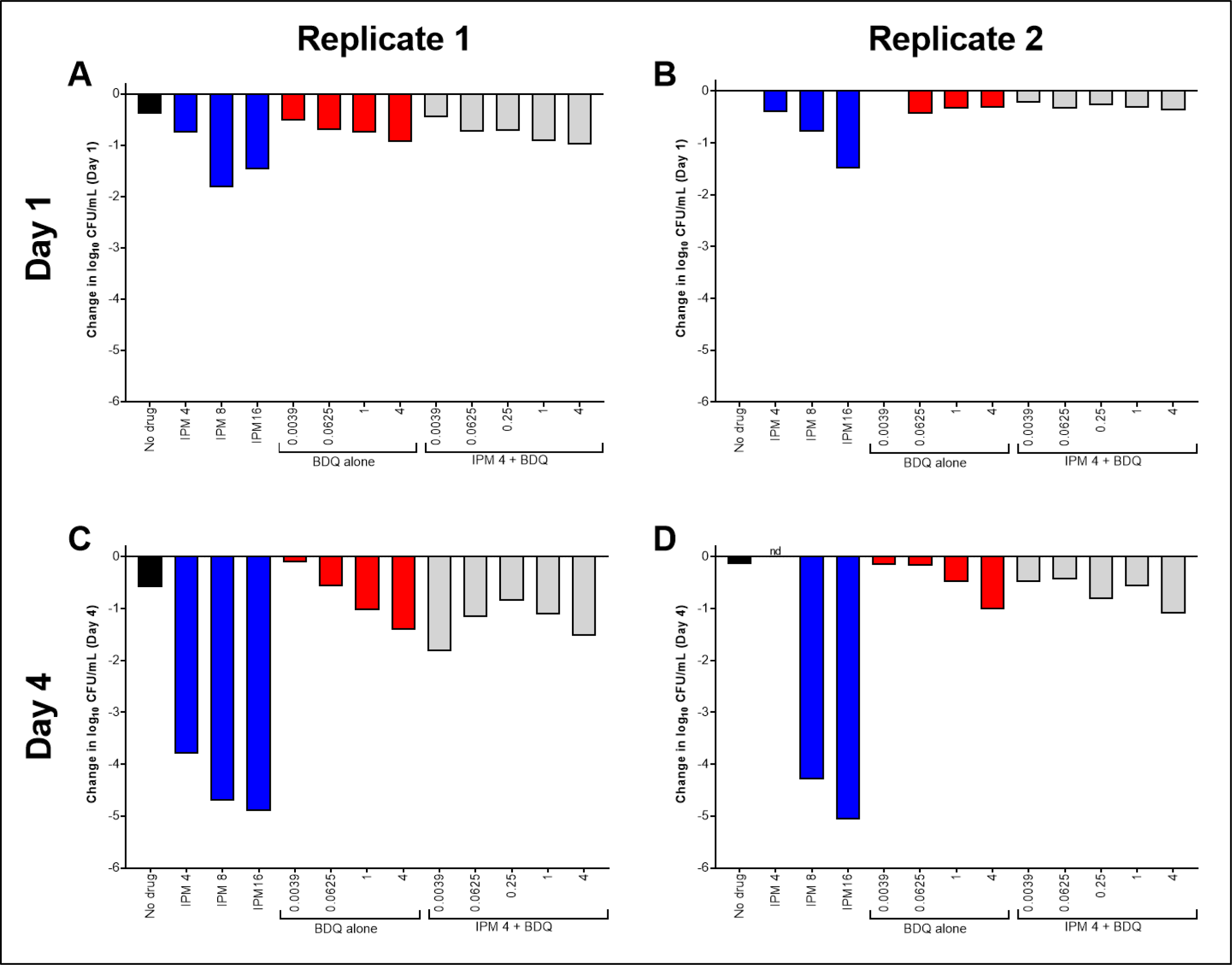
Activity of bedaquiline (BDQ) and imipenem (IPM) against nutrient-starved *M. abscessus* ATCC 19977 WT in PBS with 0.05% Tween 80. Bacteria were nutrient-starved in PBS for 20 days prior to drug exposure. Data are presented for two biological replicates (replicate 1 in panels A and C; and replicate 2 in panels B and D). The changes in log_10_ CFU/mL after 1 and 4 days of drug exposure relative to Day 0 are presented in panels A/B and C/D, respectively. Black bars indicate the no drug control; red bars indicate BDQ only samples; blue bars represent IPM only samples, and gray bars indicate IPM + BDQ samples. The number after the IPM abbreviation represents the concentration in µg/mL; for red and gray bars, the BDQ concentration in µg/mL is under each bar. Partial data from replicate 1 after 4 days of drug exposure are presented in Fig. 8. The Day 0 bacterial concentration for replicate 1 and 2 was 6.07 and 6.38 log_10_ CFU/mL, respectively. CFU values are provided in **Table S11**.

**Fig. S10.**
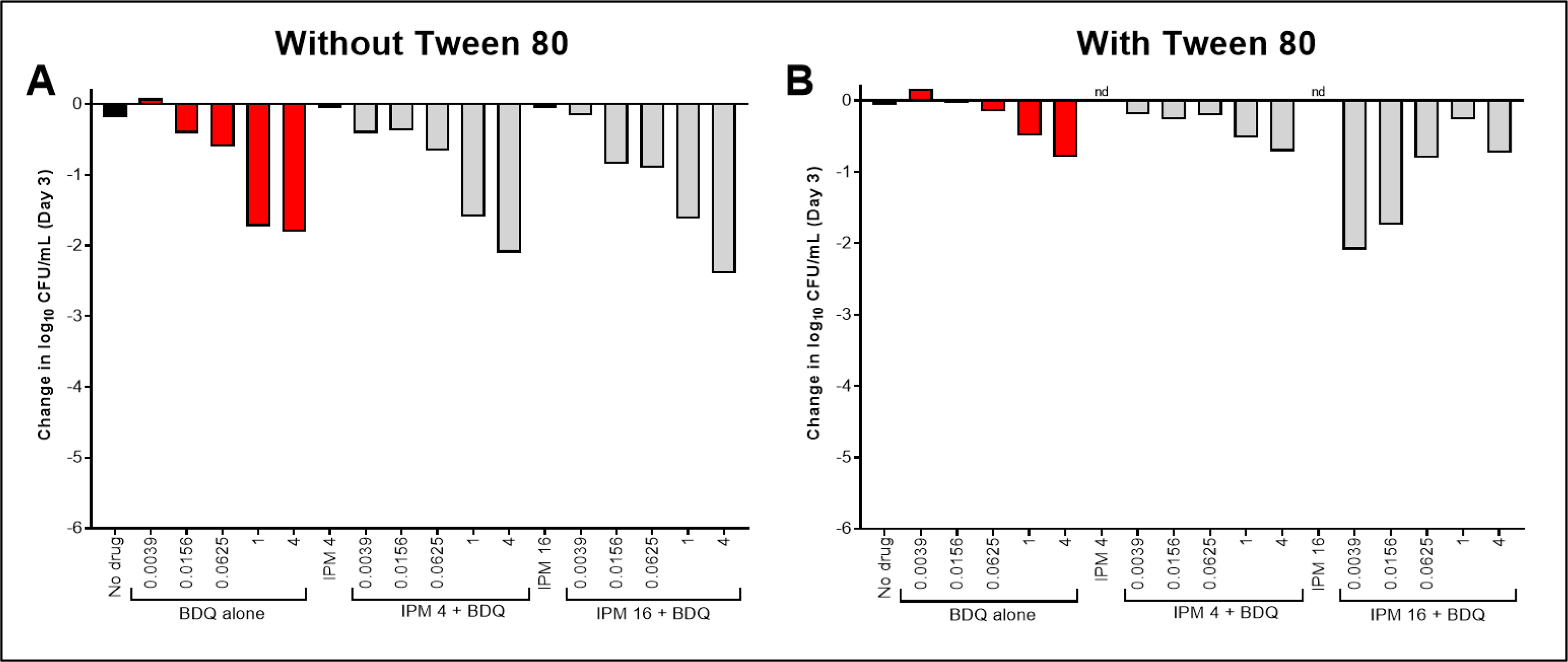
Activity of bedaquiline (BDQ) and imipenem (IPM) against nutrient-starved *M. abscessus* ATCC 19977 WT in PBS without Tween 80 (A) and with 0.05% Tween 80 (B). Bacteria were nutrient-starved in PBS for 20 days prior to drug exposure. Data are presented for two biological replicates (replicate 1 in panels A and C; and replicate 2 in panels B and D). The changes in log_10_ CFU/mL after 3 days of drug exposure relative to Day 0 are presented. Black bars indicate the no drug control; red bars indicate BDQ only samples; blue bars represent IPM only samples, and gray bars indicate IPM + BDQ samples. The number after the IPM abbreviation represents the concentration in µg/mL; for red and gray bars, the BDQ concentration in µg/mL is under each bar. The Day 0 bacterial concentration for panel A and B was 6.48 and 6.51 log10 CFU/mL, respectively. nd, not determined (samples lost). CFU values are provided in **Table S12**.

**Fig. S11.**
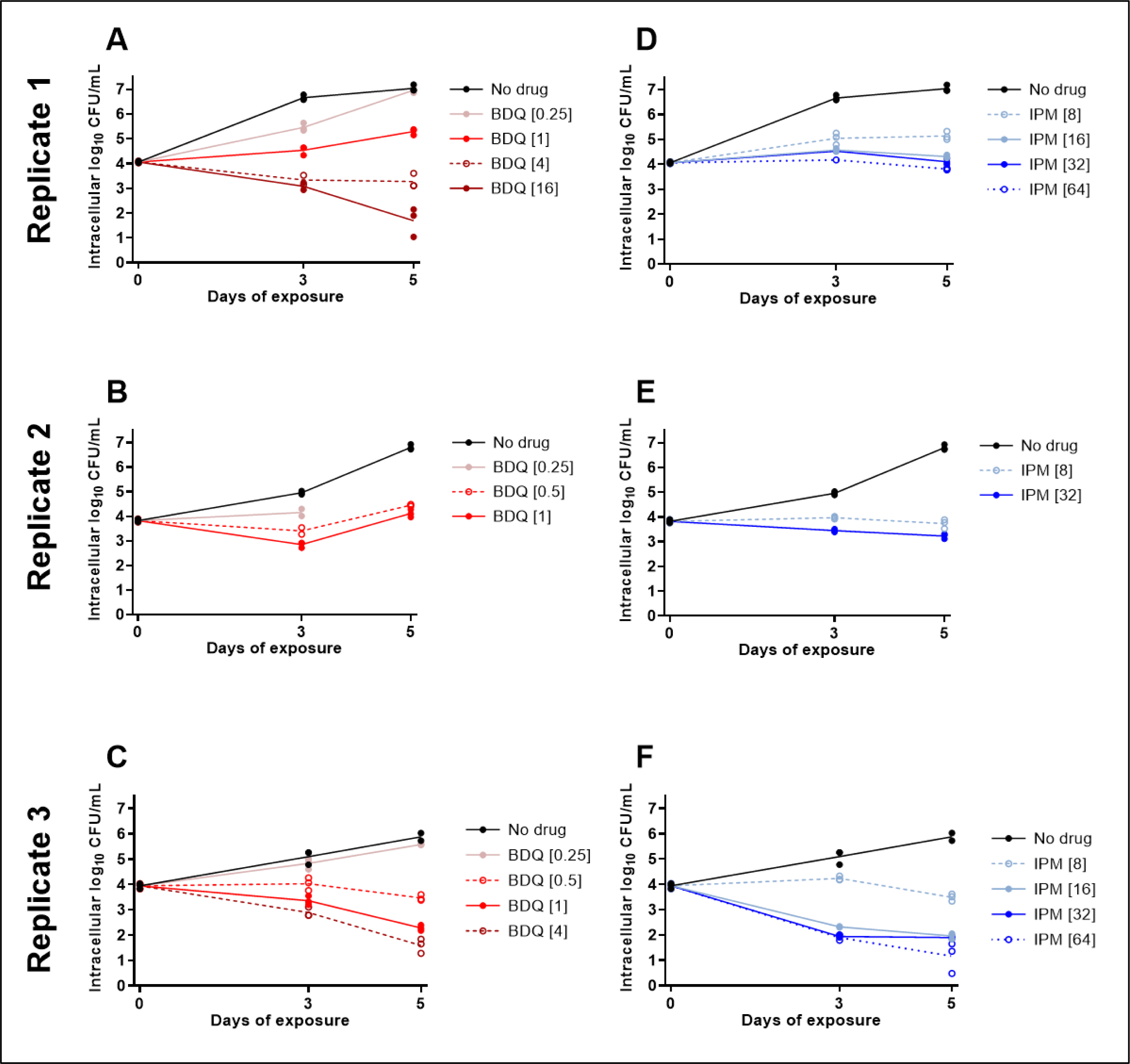
Concentration-ranging activity of bedaquiline (BDQ) (A-C) and imipenem (IPM) (D-F) alone against intra-cellular (THP-1 cells) *M. abscessus* ATCC 19977 WT. Data are presented for 3 biological replicates: replicate 1, panels A,D; replicate 2, panels B,E; replicate 3, panels C,F. Each data point represents a technical replicate, and the connecting lines pass through the mean values. The [number] after each drug abbreviation represents the concentration in µg/mL. All CFU values are provided in **Table S13**.

**Table S1.** Bedaquiline-resistant M. abscessus laboratory isolates.

| *M. abscessus*<br>isolate | Mor-<br>pho-<br>type | CFZ MIC (µg/ml) |  | BDQ MIC (µg/ml) |  | *MAB_2299c*<br>mutation | *MAB_2299c*<br>AA change | *MAB_4384*<br>mutation |
| --- | --- | --- | --- | --- | --- | --- | --- | --- |
|  |  | CAMHB | 7H9 | CAMHB | 7H9 |  |  |  |
| ATCC 19977<br>WT | WT | 0.25 | 8 | 0.0625 | 1 | None | None | None |
| OM1 | WT | 4 | 16 | 0.25 | 4 | 19C>T | Q7I | None |
| OM3 | WT | 2 | 16 | 0.5 | 4 | 19C>T | Q7I | None |
| OM4 | Rough | nd | 16 | nd | 4 | 566G>del | Frameshift | None |
| OM7 | WT | 2 | 16 | 0.5 | 4 | -18_122del | N-terminal<br>truncation | None |

WT, wild type. CFZ, clofazimine. BDQ, bedaquiline. MIC: minimum inhibitory concentration, determined by broth microdilution. CAMHB: cation-adjusted Mueller-Hinton broth; 7H9: Middlebrook 7H9 broth with 10% (v/v) Middlebrook oleic acid-albumin-dextrose-catalase supplement; AA, amino acid; nd, not determined. Tween 80 was not included in either assay medium.

**Table S2.** CFU data from samples presented in Fig. 1. This table also includes data from replicate assays that are not shown in Fig. 1. Assay medium was CAMHB with 0.05% Tween 80. Red shading indicates bedaquiline (BDQ) samples, and blue shading indicates imipenem (IPM) or IMP with avibactam (AVI) samples.

| Strain, condition,<br>Day 0 bacterial burden,<br>figure number | Drug(s) and<br>concentration(s)<br>in µg/mL | Log<sub>10</sub> CFU/mL at the<br>following time points: |  |
| --- | --- | --- | --- |
|  |  | Day 3 | Day 7 |
| **ATCC 19977 WT,**<br>CAMHB + Tween,<br>Day 0 = 5.96 log<sub>10</sub> CFU/mL,<br>**Fig. 1A** (Day 3 data); Day 7<br>data not in a figure | No drug | Clump | Clump |
|  | BDQ 0.015625 | Clump | Clump |
|  | BDQ 0.03125 | Clump | Clump |
|  | BDQ 0.0625 | 6.58 | 6.56 |
|  | BDQ 0.125 | 6.27 | 6.41 |
|  | BDQ 0.25 | 6.02 | 6.08 |
|  | BDQ 0.5 | 6.04 | 5.78 |
|  | BDQ 1 | 5.93 | 5.91 |
|  | BDQ 2 | 5.95 | 5.30 |
|  | BDQ 4 | 5.92 | 5.53 |
|  | BDQ 8 | 6.09 | 5.53 |
|  | BDQ 16 | 6.03 | 5.30 |
| **OM7 mutant,**<br>CAMHB + Tween,<br>Day 0 = 6.11 log<sub>10</sub> CFU/mL,<br>**Fig. 1B** (Day 3 data); Day 7<br>data not in a figure | No drug | Clump | Clump |
|  | BDQ 0.015625 | Clump | Clump |
|  | BDQ 0.03125 | Clump | Clump |
|  | BDQ 0.0625 | Clump | Clump |
|  | BDQ 0.125 | Clump | Clump |
|  | BDQ 0.25 | Clump | Clump |
|  | BDQ 0.5 | Clump | Clump |
|  | BDQ 1 | Clump | Clump |
|  | BDQ 2 | 5.90 | 5.95 |
|  | BDQ 4 | 6.10 | 6.11 |
|  | BDQ 8 | 6.15 | 5.72 |
|  | BDQ 16 | 5.96 | 5.72 |
| **ATCC 19977 WT,**<br>CAMHB + Tween,<br>Day 0 = 5.81 log<sub>10</sub> CFU/mL,<br>**Fig. 1C** | No drug | Clump | nd |
|  | IPM 0.5 | Clump | nd |
|  | IPM 1 | 6.45 | nd |
|  | IPM 2 | 4.62 | nd |
|  | IPM 4 | 3.17 | nd |
|  | IPM 8 | 3.09 | nd |
|  | IPM 16 | 3.02 | nd |
|  | IPM 32 | 2.81 | nd |
|  | IPM 64 | 2.82 | nd |
|  | IPM 128 | 2.86 | nd |
| **ATCC 19977 WT,**<br>CAMHB + Tween,<br>Day 0 = 6.20 log<sub>10</sub> CFU/mL,<br>Replicate not in a figure | No drug | Clump | Clump |
|  | IPM 1 | 6.64 | Clump |
|  | IPM 2 | 6.48 | Clump |
|  | IPM 4 | 6.00 | Clump |
|  | IPM 8 | 4.90 | Clump |
|  | IPM 16 | 4.41 | 6.85 |
|  | IPM 32 | 3.10 | 6.03 |
|  | IPM 64 | 3.32 | 5.41 |
|  | IPM 128 | 3.48 | 2.51 |
|  | IPM 256 | 3.87 | 1.54 |

**Table S2, continued.**
| Strain, condition,<br>Day 0 bacterial burden,<br>figure number | Drug(s) and<br>concentration(s)<br>in µg/mL | Log<sub>10</sub> CFU/mL at the<br>following time points: |  |
| --- | --- | --- | --- |
|  |  | Day 3 | Day 7 |
| **OM7 mutant,**<br>CAMHB + Tween,<br>Day 0 = 5.85 log<sub>10</sub> CFU/mL,<br>**Fig. 1D** | No drug | Clump | nd |
|  | IPM 0.5 | Clump | nd |
|  | IPM 1 | 6.16 | nd |
|  | IPM 2 | 2.33 | nd |
|  | IPM 4 | 1.23 | nd |
|  | IPM 8 | 1.49 | nd |
|  | IPM 16 | 2.17 | nd |
|  | IPM 32 | 1.11 | nd |
|  | IPM 64 | 0.85 | nd |
|  | IPM 128 | 0* | nd |
| **OM7 mutant,**<br>CAMHB + Tween,<br>Day 0 = 5.96 log<sub>10</sub> CFU/mL,<br>Replicate not in a figure | No drug | Clump | Clump |
|  | IPM 1 | 5.60 | Clump |
|  | IPM 2 | 3.00 | Clump |
|  | IPM 4 | 3.00 | 6.31 |
|  | IPM 8 | 2.30 | 2.79 |
|  | IPM 16 | 3.58 | 2.42 |
|  | IPM 32 | 3.14 | 2.66 |
|  | IPM 64 | 3.35 | 2.68 |
|  | IPM 128 | 4.15 | 2.75 |
|  | IPM 256 | 0.85 | 2.41 |
| **ATCC 19977 WT,**<br>CAMHB + Tween,<br>Day 0 = 5.81 log<sub>10</sub> CFU/mL,<br>**Fig. 1E** | No drug | Clump | nd |
|  | AVI 2 | Cump | nd |
|  | IPM 0.5 + AVI 2 | 6.37 | nd |
|  | IPM 1 + AVI 2 | 5.45 | nd |
|  | IPM 2 + AVI 2 | 2.83 | nd |
|  | IPM 4 + AVI 2 | 2.90 | nd |
|  | IPM 8 + AVI 2 | 2.81 | nd |
|  | IPM 16 + AVI 2 | 2.81 | nd |
|  | IPM 32 + AVI 2 | 2.89 | nd |
|  | IPM 64 + AVI 2 | 2.72 | nd |
|  | IPM 128 + AVI 2 | 2.66 | nd |
| **OM7 mutant,**<br>CAMHB + Tween,<br>Day 0 = 5.85 log<sub>10</sub> CFU/mL,<br>**Fig. 1F** | No drug | Clump | nd |
|  | AVI 2 | Cump | nd |
|  | IPM 0.5 + AVI 2 | 6.22 | nd |
|  | IPM 1 + AVI 2 | 5.45 | nd |
|  | IPM 2 + AVI 2 | 2.87 | nd |
|  | IPM 4 + AVI 2 | 3.72 | nd |
|  | IPM 8 + AVI 2 | 3.15 | nd |
|  | IPM 16 + AVI 2 | 3.10 | nd |
|  | IPM 32 + AVI 2 | 3.04 | nd |
|  | IPM 64 + AVI 2 | 3.02 | nd |
|  | IPM 128 + AVI 2 | 2.78 | nd |

"Clump" indicates that the bacteria had overgrown and formed clumps, precluding CFU quantification. nd, not determined.
\*Lower limit of detection: 0.48 log<sub>10</sub> CFU/mL

**Table S3.** CFU data from samples presented in Fig. 2. Assay medium was CAMHB with 0.05% Tween 80. Red shading indicates bedaquiline (BDQ) samples, blue shading indicates imipenem (IPM) or IMP with avibactam (AVI) samples, and gray shading indicates IPM + AVI + BDQ samples.

| Strain, condition,<br>Day 0 bacterial burden,<br>figure number | Drug(s) and<br>concentration(s)<br>in µg/mL | Log<sub>10</sub> CFU/mL at the<br>following time points: |  |
| --- | --- | --- | --- |
|  |  | Day 3 | Day 5 |
| **ATCC 19977 WT**,<br>CAMHB + Tween,<br>Day 0 = 5.57 log<sub>10</sub> CFU/mL,<br>**Fig. 2A** (Day 3 data);<br>Day 5 data and<br>data from additional<br>combinations not included<br>in a figure | No drug | Clump | Clump |
|  | BDQ 0.5 | 5.45 | Clump |
|  | BDQ 1 | 5.30 | Clump |
|  | AVI 2 | Clump | Clump |
|  | IPM 2 | Clump | Clump |
|  | IPM 4 | 5.26 | Clump |
|  | IPM 2 + AVI 2 | 3.78 | Clump |
|  | IPM 4 + AVI 2 | 3.30 | Clump |
|  | IPM 2 + AVI 2 + BDQ 0.00390625 | 3.60 | <3.30* |
|  | IPM 2 + AVI 2 + BDQ 0.015625 | 5.14 | 5.56 |
|  | IPM 2 + AVI 2 + BDQ 0.0625 | 5.26 | 5.03 |
|  | IPM 2 + AVI 2 + BDQ 0.25 | 5.82 | 5.45 |
|  | IPM 2 + AVI 2 + BDQ 0.5 | 5.88 | 5.73 |
|  | IPM 2 + AVI 2 + BDQ 1 | 5.99 | 5.81 |
|  | IPM 4 + AVI 2 + BDQ 0.00390625 | 3.26 | Clump |
|  | IPM 4 + AVI 2 + BDQ 0.015625 | 4.13 | Clump |
|  | IPM 4 + AVI 2 + BDQ 0.0625 | 5.26 | Clump |
|  | IPM 4 + AVI 2 + BDQ 0.25 | 5.66 | Clump |
|  | IPM 4 + AVI 2 + BDQ 0.5 | 5.72 | Clump |
|  | IPM 4 + AVI 2 + BDQ 1 | 5.81 | Clump |
| **OM7 mutant**,<br>CAMHB + Tween,<br>Day 0 = 5.76 log<sub>10</sub> CFU/mL,<br>**Fig. 2B** (Day 3 data);<br>Day 5 data and<br>data from additional<br>combinations not included<br>in a figure | No drug | Clump | Clump |
|  | BDQ 0.5 | 7.06 | Clump |
|  | BDQ 1 | 6.12 | Clump |
|  | AVI 2 | Clump | Clump |
|  | IPM 2 | 6.18 | Clump |
|  | IPM 4 | 5.08 | Clump |
|  | IPM 2 + AVI 2 | 3.60 | Clump |
|  | IPM 4 + AVI 2 | 4.00 | Clump |
|  | IPM 2 + AVI 2 + BDQ 0.00390625 | 2.78 | 5.58 |
|  | IPM 2 + AVI 2 + BDQ 0.015625 | 3.20 | 5.56 |
|  | IPM 2 + AVI 2 + BDQ 0.0625 | 4.89 | Clump |
|  | IPM 2 + AVI 2 + BDQ 0.25 | 5.45 | Clump |
|  | IPM 2 + AVI 2 + BDQ 0.5 | 5.90 | Clump |
|  | IPM 2 + AVI 2 + BDQ 1 | 5.75 | Clump |
|  | IPM 4 + AVI 2 + BDQ 0.00390625 | 4.87 | 4.82 |
|  | IPM 4 + AVI 2 + BDQ 0.015625 | 2.78 | 4.99 |
|  | IPM 4 + AVI 2 + BDQ 0.0625 | 4.25 | 6.24 |
|  | IPM 4 + AVI 2 + BDQ 0.25 | 5.30 | Clump |
|  | IPM 4 + AVI 2 + BDQ 0.5 | 5.15 | 5.64 |
|  | IPM 4 + AVI 2 + BDQ 1 | 5.48 | 6.66 |

"Clump" indicates that the bacteria had overgrown and formed clumps, precluding CFU quantification.
\*The lower limit of detection for this sample was 3.30 log<sub>10</sub> CFU/mL.

**Table S4.** CFU data from samples presented in Fig. S2. Assay medium was CAMHB with 0.05% Tween 80. Gray shading indicates imipenem (IPM) + avibactam (AVI) + bedaquiline (BDQ) samples.

| Strain, condition,<br>Day 0 bacterial burden,<br>figure number | Drug(s) and<br>concentration(s)<br>in µg/mL | Log<sub>10</sub> CFU/mL at the<br>following time points: |  |
| --- | --- | --- | --- |
|  |  | Day 3 | Day 5 |
| **ATCC 19977 WT,**<br>CAMHB + Tween,<br>Day 0 = 6.13 log<sub>10</sub> CFU/mL,<br>**Fig. S2A** (Day 3 data);<br>Day 5 data not included<br>in a figure | No drug | Clump | Clump |
|  | IPM 2 + AVI 2 + BDQ 0.00390625 | 5.20 | 7.15 |
|  | IPM 2 + AVI 2 + BDQ 0.0078125 | 5.03 | 6.45 |
|  | IPM 2 + AVI 2 + BDQ 0.015625 | 4.34 | 6.79 |
|  | IPM 2 + AVI 2 + BDQ 0.03125 | 5.70 | 6.56 |
|  | IPM 2 + AVI 2 + BDQ 0.0625 | 5.79 | 6.30 |
|  | IPM 2 + AVI 2 + BDQ 0.125 | 6.01 | 6.20 |
|  | IPM 2 + AVI 2 + BDQ 0.25 | 5.89 | 6.58 |
|  | IPM 2 + AVI 2 + BDQ 0.5 | 5.87 | 5.76 |
|  | IPM 2 + AVI 2 + BDQ 1 | 5.81 | 6.34 |
|  | IPM 4 + AVI 2 + BDQ 0.00390625 | 5.16 | 7.20 |
|  | IPM 4 + AVI 2 + BDQ 0.0078125 | <3.30* | 5.82 |
|  | IPM 4 + AVI 2 + BDQ 0.015625 | 5.11 | 6.51 |
|  | IPM 4 + AVI 2 + BDQ 0.03125 | 5.23 | 6.45 |
|  | IPM 4 + AVI 2 + BDQ 0.0625 | 5.62 | 6.29 |
|  | IPM 4 + AVI 2 + BDQ 0.125 | 5.79 | 6.87 |
|  | IPM 4 + AVI 2 + BDQ 0.25 | 5.86 | 5.82 |
|  | IPM 4 + AVI 2 + BDQ 0.5 | 5.81 | 5.87 |
|  | IPM 4 + AVI 2 + BDQ 1 | 5.89 | 5.97 |
| **OM7 mutant,**<br>CAMHB + Tween,<br>Day 0 = 6.22 log<sub>10</sub> CFU/mL,<br>**Fig. S2B** (Day 3 data);<br>Day 5 data not included<br>in a figure | No drug | Clump | Clump |
|  | IPM 2 + AVI 2 + BDQ 0.00390625 | 4.60 | 6.72 |
|  | IPM 2 + AVI 2 + BDQ 0.0078125 | 4.48 | 6.56 |
|  | IPM 2 + AVI 2 + BDQ 0.015625 | 4.76 | 6.56 |
|  | IPM 2 + AVI 2 + BDQ 0.03125 | 4.70 | 6.64 |
|  | IPM 2 + AVI 2 + BDQ 0.0625 | 4.97 | 6.53 |
|  | IPM 2 + AVI 2 + BDQ 0.125 | 5.51 | 6.70 |
|  | IPM 2 + AVI 2 + BDQ 0.25 | 5.64 | 5.94 |
|  | IPM 2 + AVI 2 + BDQ 0.5 | 5.76 | 6.15 |
|  | IPM 2 + AVI 2 + BDQ 1 | 5.89 | 5.72 |
|  | IPM 4 + AVI 2 + BDQ 0.00390625 | 3.38 | 6.70 |
|  | IPM 4 + AVI 2 + BDQ 0.0078125 | 3.15 | 5.30 |
|  | IPM 4 + AVI 2 + BDQ 0.015625 | 3.00 | 5.09 |
|  | IPM 4 + AVI 2 + BDQ 0.03125 | 3.70 | 6.19 |
|  | IPM 4 + AVI 2 + BDQ 0.0625 | 4.41 | 5.03 |
|  | IPM 4 + AVI 2 + BDQ 0.125 | 5.18 | 5.27 |
|  | IPM 4 + AVI 2 + BDQ 0.25 | 5.48 | 5.96 |
|  | IPM 4 + AVI 2 + BDQ 0.5 | 5.53 | 6.45 |
|  | IPM 4 + AVI 2 + BDQ 1 | 5.72 | 5.57 |

"Clump" indicates that the bacteria had overgrown and formed clumps, precluding CFU quantification.
\*The lower limit of detection for this sample was 3.30 log<sub>10</sub> CFU/mL. The value of 3.30 log<sub>10</sub> CFU/mL was used to calculate the change in CFU from Day 0, presented in **Fig. S2**.

**Table S5.** CFU data from samples presented in Fig. 3. Assay medium was CAMHB with 0.05% Tween 80. Red shading indicates bedaquiline (BDQ) samples; blue shading indicates meropenem (MEM), MmpL3 inhibitor (MPL), or clarithromycin (CLR) samples; and gray shading indicates BDQ combination samples. MPL is N-4S-methylcyclohexyl-4,6-dimethyl-1H-indole-2-carboxamide.

| Strain, condition,<br>Day 0 bacterial burden,<br>figure number | Drug(s)/compound(s) and<br>concentration(s)<br>in µg/mL | Log<sub>10</sub> CFU/mL at the following time points: |  |  |
| --- | --- | --- | --- | --- |
|  |  | Day 3 | Day 5 | Day 8 |
| ATCC 19977 WT,<br>CAMHB + Tween,<br>Day 0 = 6.40 log<sub>10</sub> CFU/mL,<br>Fig. 3 | No drug | 7.59 | Clump | Clump |
|  | BDQ 0.00390625 | 6.77 | 7.20 | 6.43 |
|  | BDQ 0.015625 | 7.15 | 7.67 | 6.59 |
|  | BDQ 0.125 | 6.41 | 6.48 | 5.40 |
|  | BDQ 1 | 5.34 | 5.36 | 5.04 |
|  | BDQ 4 | 5.20 | 5.26 | 4.85 |
|  | MEM 16 | 3.23 | 2.48 | 2.73 |
|  | MEM 16 + BDQ 0.00390625 | 3.98 | 2.66 | 4.78 |
|  | MEM 16 + BDQ 0.015625 | 3.76 | 2.32 | 4.85 |
|  | MEM 16 + BDQ 0.125 | 5.34 | 4.95 | 4.20 |
|  | MEM 16 + BDQ 1 | 3.76 | 5.15 | 4.85 |
|  | MEM 16 + BDQ 4 | 5.20 | 5.00 | 4.78 |
|  | MPL 2 | 4.11 | 2.30 | 1.04 |
|  | MPL 2 + BDQ 0.00390625 | 5.26 | 2.85 | 2.38 |
|  | MPL 2 + BDQ 0.015625 | 6.34 | 5.70 | 6.48 |
|  | MPL 2 + BDQ 0.125 | 6.15 | 6.56 | 6.53 |
|  | MPL 2 + BDQ 1 | 5.40 | 5.34 | 5.39 |
|  | MPL 2 + BDQ 4 | 5.43 | 5.26 | 5.05 |
|  | CLR 4 | 4.78 | 3.52 | 3.27 |
|  | CLR 4 + BDQ 0.015625 | 4.65 | 3.51 | 3.20 |
|  | CLR 4 + BDQ 0.125 | 4.90 | 3.43 | 2.23 |
|  | CLR 4 + BDQ 1 | 4.91 | 3.45 | 1.61 |
|  | CLR 4 + BDQ 4 | 4.79 | 3.40 | 1.32 |

"Clump" indicates that the bacteria had overgrown and formed clumps, precluding CFU quantification.

**Table S6.** CFU and relative light unit (RLU) data from samples presented in Fig. 4 and Fig. S3. Assay medium was CAMHB with 0.05% Tween 80. Red shading indicates bedaquiline (BDQ) samples, blue shading indicates imipenem (IPM) samples, and gray shading indicates IPM + BDQ samples.

| Strain, condition,<br>Day 0 bacterial burden,<br>figure number | Drug(s) and<br>concentration(s)<br>in µg/mL | Log<sub>10</sub> CFU/mL at the<br>following time points: |  | Log<sub>10</sub> RLU/mL at the<br>following time points: |  | MegaRLU/CFU at the<br>following time points: |  |
| --- | --- | --- | --- | --- | --- | --- | --- |
|  |  | Day 1 | Day 3 | Day 1 | Day 3 | Day 1 | Day 3 |
| **ATCC 19977 WT,**<br>CAMHB + Tween,<br>Day 0 = 6.25 log<sub>10</sub> CFU/mL,<br>**Fig. 4A,C; Fig. S3A,C**<br>(Replicate 1) | No drug | 6.73 | 7.31 | 3.28 | 3.67 | 356 | 228 |
|  | BDQ 0.0625 | 6.72 | 7.24 | 3.27 | 3.58 | 355 | 217 |
|  | BDQ 1 | 6.68 | 6.70 | 2.95 | 3.17 | 185 | 296 |
|  | IPM 4 | 6.12 | 5.66 | 3.78 | 3.61 | 4576 | 8945 |
|  | IPM 16 | 4.34 | 4.51 | 3.32 | 3.32 | 95273 | 64547 |
|  | IPM 4 + BDQ 0.00390625 | 6.06 | 5.16 | 3.84 | 3.60 | 5966 | 27435 |
|  | IPM 4 + BDQ 0.0078125 | 6.32 | 4.53 | 3.88 | 3.63 | 3648 | 124618 |
|  | IPM 4 + BDQ 0.0625 | 6.62 | 6.51 | 3.74 | 3.77 | 1311 | 1830 |
|  | IPM 4 + BDQ 0.25 | 6.58 | 6.58 | 3.24 | 3.57 | 461 | 971 |
|  | IPM 4 + BDQ 1 | 6.78 | 6.70 | 2.97 | 3.24 | 156 | 351 |
|  | IPM 4 + BDQ 4 | 6.76 | 6.75 | 3.13 | 3.25 | 231 | 319 |
|  | IPM 16 + BDQ 0.00390625 | 4.56 | 3.48 | 3.33 | 3.33 | 59389 | 712667 |
|  | IPM 16 + BDQ 0.0625 | 6.11 | 5.78 | 3.41 | 3.41 | 2026 | 4322 |
|  | IPM 16 + BDQ 0.25 | 6.66 | 6.34 | 3.51 | 3.51 | 697 | 1457 |
|  | IPM 16 + BDQ 1 | 6.62 | 6.53 | 3.25 | 3.25 | 421 | 520 |
| **ATCC 19977 WT,**<br>CAMHB + Tween,<br>Day 0 = 7.03 log<sub>10</sub> CFU/mL,<br>**Fig. 4B,D; Fig. S3B,D**<br>(Replicate 2) | No drug | 7.06 | 7.42 | 3.44 | 3.60 | 242 | 152 |
|  | BDQ 0.0625 | 6.79 | 7.23 | 3.23 | 3.44 | 273 | 162 |
|  | BDQ 1 | 7.09 | 7.01 | 2.52 | 2.82 | 27 | 65 |
|  | IPM 4 | 6.40 | 5.38 | 3.69 | 3.44 | 1910 | 11632 |
|  | IPM 16 | 6.21 | 3.87 | 3.28 | 2.97 | 1169 | 125338 |
|  | IPM 4 + BDQ 0.00390625 | 6.60 | 4.83 | 3.67 | 3.34 | 1170 | 32228 |
|  | IPM 4 + BDQ 0.0078125 | 6.26 | 5.61 | 3.64 | 3.45 | 2433 | 6934 |
|  | IPM 4 + BDQ 0.0625 | 6.89 | 6.79 | 3.58 | 3.54 | 485 | 556 |
|  | IPM 4 + BDQ 0.25 | 6.89 | 6.89 | 3.26 | 3.27 | 236 | 241 |
|  | IPM 4 + BDQ 1 | 7.02 | 6.91 | 2.66 | 2.90 | 44 | 96 |
|  | IPM 4 + BDQ 4 | 6.86 | 6.86 | 2.53 | 2.79 | 47 | 85 |
|  | IPM 16 + BDQ 0.00390625 | 6.26 | 4.00 | 3.32 | 3.00 | 1125 | 100050 |
|  | IPM 16 + BDQ 0.0625 | 6.82 | 6.45 | 3.30 | 3.25 | 304 | 630 |
|  | IPM 16 + BDQ 0.25 | 6.97 | 6.87 | 3.06 | 3.18 | 124 | 205 |
|  | IPM 16 + BDQ 1 | 7.00 | 6.83 | 2.73 | 2.95 | 54 | 131 |
|  | IPM 16 + BDQ 4 | 6.92 | 6.91 | 2.59 | 2.87 | 46 | 91 |

**Table S7.** CFU data from samples presented in Fig. 5. This table also includes data from replicate assays that are not shown in Fig. 5. Assay medium was CAMHB without Tween 80. Red shading indicates bedaquiline (BDQ) samples, and blue shading indicates imipenem (IPM) samples.

| Strain, condition,<br>Day 0 bacterial burden,<br>figure number | Drug(s) and<br>concentration(s)<br>in µg/mL | Log<sub>10</sub> CFU/mL<br>at Day 3 |
| --- | --- | --- |
| ATCC 19977 WT,<br>CAMHB (no Tween),<br>Day 0 = 5.90 log<sub>10</sub> CFU/mL,<br>**Fig. 5A**, with additional<br>concentrations not in the<br>figure | No drug | Clump |
|  | BDQ 0.0078125 | Clump |
|  | BDQ 0.015625 | 6.44 |
|  | BDQ 0.03125 | 5.83 |
|  | BDQ 0.0625 | 5.53 |
|  | BDQ 0.125 | 5.23 |
|  | BDQ 0.25 | 5.25 |
|  | BDQ 0.5 | 5.11 |
|  | BDQ 1 | 5.05 |
|  | BDQ 2 | 5.00 |
|  | BDQ 4 | 5.08 |
|  | BDQ 8 | 5.29 |
|  | BDQ 16 | 5.23 |
| OM7 mutant,<br>CAMHB (no Tween),<br>Day 0 = 5.08 log<sub>10</sub> CFU/mL,<br>**Fig. 5B**, with additional<br>concentrations not in the<br>figure | No drug | Clump |
|  | BDQ 0.0078125 | Clump |
|  | BDQ 0.015625 | Clump |
|  | BDQ 0.03125 | Clump |
|  | BDQ 0.0625 | Clump |
|  | BDQ 0.125 | Clump |
|  | BDQ 0.25 | Clump |
|  | BDQ 0.5 | 5.70 |
|  | BDQ 1 | 5.66 |
|  | BDQ 2 | 5.66 |
|  | BDQ 4 | 5.79 |
|  | BDQ 8 | 5.85 |
|  | BDQ 16 | 5.95 |
| ATCC 19977 WT,<br>CAMHB (no Tween),<br>Day 0 = 5.91 log<sub>10</sub> CFU/mL,<br>**Fig. 5C** | No drug | Clump |
|  | IPM 1 | Clump |
|  | IPM 2 | Clump |
|  | IPM 4 | Clump |
|  | IPM 8 | Clump |
|  | IPM 16 | 6.01 |
|  | IPM 32 | 5.79 |
|  | IPM 64 | 5.62 |
|  | IPM 128 | 5.15 |
|  | IPM 256 | 4.73 |
| ATCC 19977 WT,<br>CAMHB (no Tween),<br>Day 0 = 5.28 log<sub>10</sub> CFU/mL,<br>Replicate not in a figure | No drug | Clump |
|  | IPM 1 | Clump |
|  | IPM 2 | Clump |
|  | IPM 4 | Clump |
|  | IPM 8 | Clump |
|  | IPM 16 | 6.06 |
|  | IPM 32 | 6.12 |
|  | IPM 64 | 5.00 |
|  | IPM 128 | 4.78 |
|  | IPM 256 | 4.33 |

**Table S7, continued.**
| Strain, condition,<br>Day 0 bacterial burden,<br>figure number | Drug(s) and<br>concentration(s)<br>in µg/mL | Log<sub>10</sub> CFU/mL<br>at Day 3 |
| --- | --- | --- |
| **OM7 mutant,**<br>CAMHB (no Tween),<br>Day 0 = 6.20 log<sub>10</sub> CFU/mL,<br>**Fig. 5D** | No drug | Clump |
|  | IPM 1 | 6.58 |
|  | IPM 2 | 6.64 |
|  | IPM 4 | 5.91 |
|  | IPM 8 | 5.38 |
|  | IPM 16 | 5.41 |
|  | IPM 32 | 5.45 |
|  | IPM 64 | 5.00 |
|  | IPM 128 | 4.86 |
|  | IPM 256 | 4.62 |
| **OM7 mutant,**<br>CAMHB (no Tween),<br>Day 0 = 5.72 log<sub>10</sub> CFU/mL,<br>Replicate not in a figure | No drug | Clump |
|  | IPM 1 | 5.66 |
|  | IPM 2 | 5.22 |
|  | IPM 4 | 5.13 |
|  | IPM 8 | 5.05 |
|  | IPM 16 | 5.21 |
|  | IPM 32 | 3.94 |
|  | IPM 64 | 4.97 |
|  | IPM 128 | 4.51 |
|  | IPM 256 | 4.40 |

“Clump” indicates that the bacteria had overgrown and formed clumps, precluding CFU quantification.

**Table S8.**
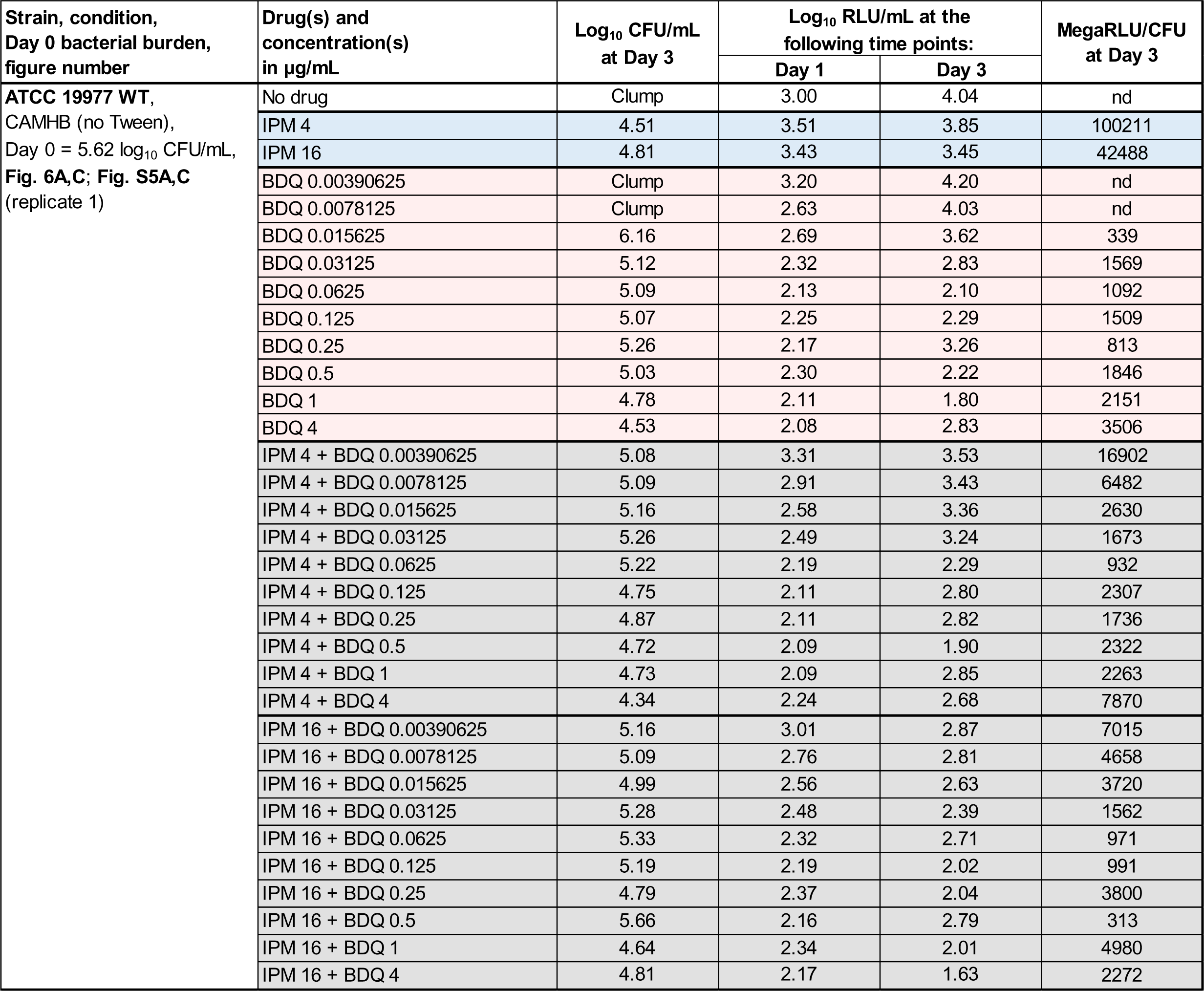
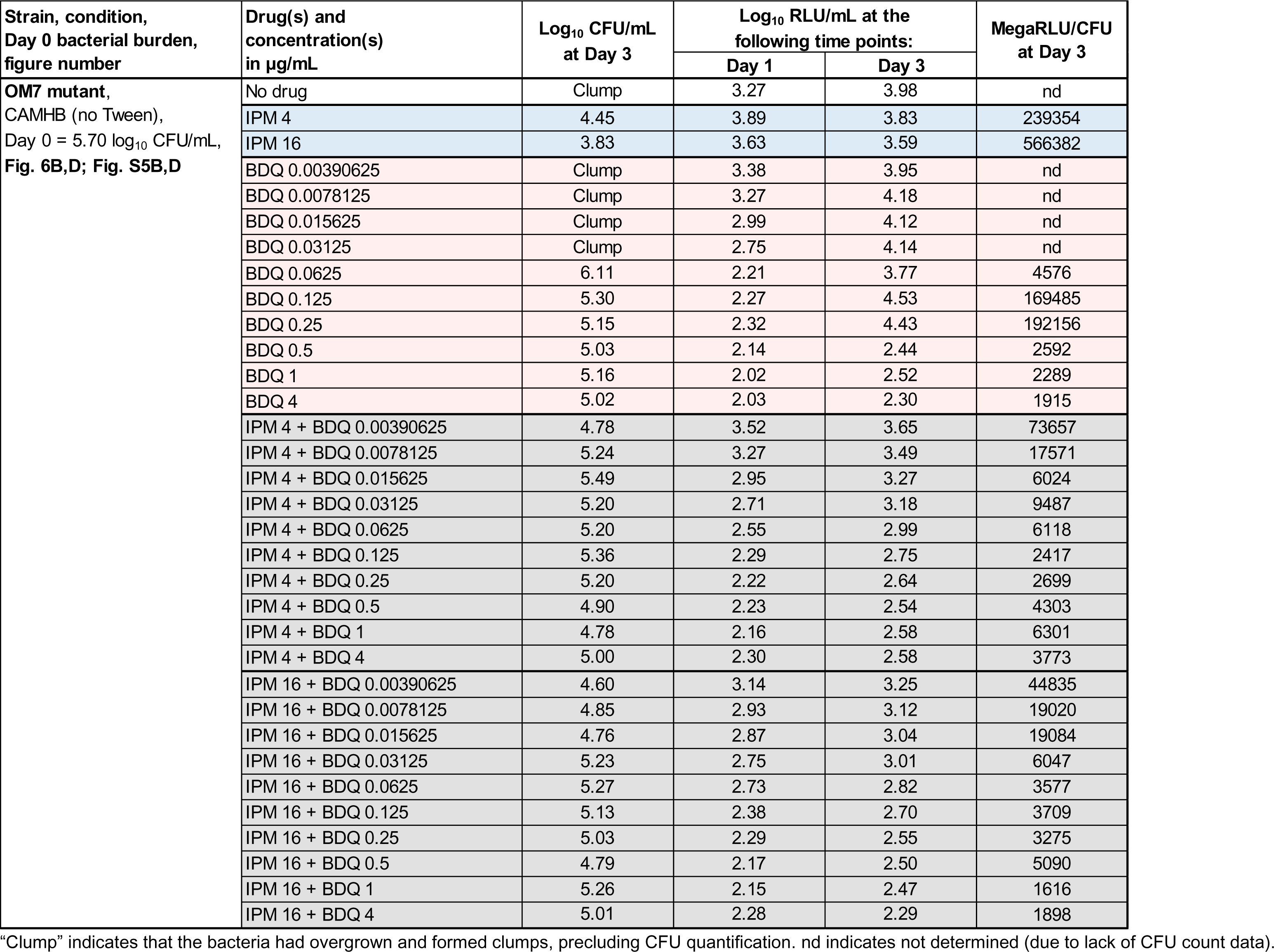
CFU and relative light unit (RLU) data from samples presented in Fig. 6 and Fig. S5. Assay medium was CAMHB without Tween 80. Red shading indicates bedaquiline (BDQ) samples, blue shading indicates imipenem (IPM) samples, and gray shading indicates IPM + BDQ samples.

| Strain, condition,<br>Day 0 bacterial burden,<br>figure number | Drug(s) and<br>concentration(s)<br>in µg/mL | Log<sub>10</sub> CFU/mL<br>at Day 3 | Log<sub>10</sub> RLU/mL at the<br>following time points: |  | MegaRLU/CFU<br>at Day 3 |
| --- | --- | --- | --- | --- | --- |
|  |  |  | Day 1 | Day 3 |  |
| ATCC 19977 WT,<br>CAMHB (no Tween),<br>Day 0 = 5.62 log<sub>10</sub> CFU/mL,<br>Fig. 6A,C; Fig. S5A,C<br>(replicate 1) | No drug | Clump | 3.00 | 4.04 | nd |
|  | IPM 4 | 4.51 | 3.51 | 3.85 | 100211 |
|  | IPM 16 | 4.81 | 3.43 | 3.45 | 42488 |
|  | BDQ 0.00390625 | Clump | 3.20 | 4.20 | nd |
|  | BDQ 0.0078125 | Clump | 2.63 | 4.03 | nd |
|  | BDQ 0.015625 | 6.16 | 2.69 | 3.62 | 339 |
|  | BDQ 0.03125 | 5.12 | 2.32 | 2.83 | 1569 |
|  | BDQ 0.0625 | 5.09 | 2.13 | 2.10 | 1092 |
|  | BDQ 0.125 | 5.07 | 2.25 | 2.29 | 1509 |
|  | BDQ 0.25 | 5.26 | 2.17 | 3.26 | 813 |
|  | BDQ 0.5 | 5.03 | 2.30 | 2.22 | 1846 |
|  | BDQ 1 | 4.78 | 2.11 | 1.80 | 2151 |
|  | BDQ 4 | 4.53 | 2.08 | 2.83 | 3506 |
|  | IPM 4 + BDQ 0.00390625 | 5.08 | 3.31 | 3.53 | 16902 |
|  | IPM 4 + BDQ 0.0078125 | 5.09 | 2.91 | 3.43 | 6482 |
|  | IPM 4 + BDQ 0.015625 | 5.16 | 2.58 | 3.36 | 2630 |
|  | IPM 4 + BDQ 0.03125 | 5.26 | 2.49 | 3.24 | 1673 |
|  | IPM 4 + BDQ 0.0625 | 5.22 | 2.19 | 2.29 | 932 |
|  | IPM 4 + BDQ 0.125 | 4.75 | 2.11 | 2.80 | 2307 |
|  | IPM 4 + BDQ 0.25 | 4.87 | 2.11 | 2.82 | 1736 |
|  | IPM 4 + BDQ 0.5 | 4.72 | 2.09 | 1.90 | 2322 |
|  | IPM 4 + BDQ 1 | 4.73 | 2.09 | 2.85 | 2263 |
|  | IPM 4 + BDQ 4 | 4.34 | 2.24 | 2.68 | 7870 |
|  | IPM 16 + BDQ 0.00390625 | 5.16 | 3.01 | 2.87 | 7015 |
|  | IPM 16 + BDQ 0.0078125 | 5.09 | 2.76 | 2.81 | 4658 |
|  | IPM 16 + BDQ 0.015625 | 4.99 | 2.56 | 2.63 | 3720 |
|  | IPM 16 + BDQ 0.03125 | 5.28 | 2.48 | 2.39 | 1562 |
|  | IPM 16 + BDQ 0.0625 | 5.33 | 2.32 | 2.71 | 971 |
|  | IPM 16 + BDQ 0.125 | 5.19 | 2.19 | 2.02 | 991 |
|  | IPM 16 + BDQ 0.25 | 4.79 | 2.37 | 2.04 | 3800 |
|  | IPM 16 + BDQ 0.5 | 5.66 | 2.16 | 2.79 | 313 |
|  | IPM 16 + BDQ 1 | 4.64 | 2.34 | 2.01 | 4980 |
|  | IPM 16 + BDQ 4 | 4.81 | 2.17 | 1.63 | 2272 |
|  |  |  | Day 1 | Day 3 |  |
| OM7 mutant,<br>CAMHB (no Tween),<br>Day 0 = 5.70 log<sub>10</sub> CFU/mL,<br>Fig. 6B,D; Fig. S5B,D | No drug | Clump | 3.27 | 3.98 | nd |
|  | IPM 4 | 4.45 | 3.89 | 3.83 | 239354 |
|  | IPM 16 | 3.83 | 3.63 | 3.59 | 566382 |
|  | BDQ 0.00390625 | Clump | 3.38 | 3.95 | nd |
|  | BDQ 0.0078125 | Clump | 3.27 | 4.18 | nd |
|  | BDQ 0.015625 | Clump | 2.99 | 4.12 | nd |
|  | BDQ 0.03125 | Clump | 2.75 | 4.14 | nd |
|  | BDQ 0.0625 | 6.11 | 2.21 | 3.77 | 4576 |
|  | BDQ 0.125 | 5.30 | 2.27 | 4.53 | 169485 |
|  | BDQ 0.25 | 5.15 | 2.32 | 4.43 | 192156 |
|  | BDQ 0.5 | 5.03 | 2.14 | 2.44 | 2592 |
|  | BDQ 1 | 5.16 | 2.02 | 2.52 | 2289 |
|  | BDQ 4 | 5.02 | 2.03 | 2.30 | 1915 |
|  | IPM 4 + BDQ 0.00390625 | 4.78 | 3.52 | 3.65 | 73657 |
|  | IPM 4 + BDQ 0.0078125 | 5.24 | 3.27 | 3.49 | 17571 |
|  | IPM 4 + BDQ 0.015625 | 5.49 | 2.95 | 3.27 | 6024 |
|  | IPM 4 + BDQ 0.03125 | 5.20 | 2.71 | 3.18 | 9487 |
|  | IPM 4 + BDQ 0.0625 | 5.20 | 2.55 | 2.99 | 6118 |
|  | IPM 4 + BDQ 0.125 | 5.36 | 2.29 | 2.75 | 2417 |
|  | IPM 4 + BDQ 0.25 | 5.20 | 2.22 | 2.64 | 2699 |
|  | IPM 4 + BDQ 0.5 | 4.90 | 2.23 | 2.54 | 4303 |
|  | IPM 4 + BDQ 1 | 4.78 | 2.16 | 2.58 | 6301 |
|  | IPM 4 + BDQ 4 | 5.00 | 2.30 | 2.58 | 3773 |
|  | IPM 16 + BDQ 0.00390625 | 4.60 | 3.14 | 3.25 | 44835 |
|  | IPM 16 + BDQ 0.0078125 | 4.85 | 2.93 | 3.12 | 19020 |
|  | IPM 16 + BDQ 0.015625 | 4.76 | 2.87 | 3.04 | 19084 |
|  | IPM 16 + BDQ 0.03125 | 5.23 | 2.75 | 3.01 | 6047 |
|  | IPM 16 + BDQ 0.0625 | 5.27 | 2.73 | 2.82 | 3577 |
|  | IPM 16 + BDQ 0.125 | 5.13 | 2.38 | 2.70 | 3709 |
|  | IPM 16 + BDQ 0.25 | 5.03 | 2.29 | 2.55 | 3275 |
|  | IPM 16 + BDQ 0.5 | 4.79 | 2.17 | 2.50 | 5090 |
|  | IPM 16 + BDQ 1 | 5.26 | 2.15 | 2.47 | 1616 |
|  | IPM 16 + BDQ 4 | 5.01 | 2.28 | 2.29 | 1898 |

"Clump" indicates that the bacteria had overgrown and formed clumps, precluding CFU quantification. nd indicates not determined (due to lack of CFU count data).

**Table S9.** CFU and relative light unit (RLU) data from samples presented in Fig. S4. Assay medium was CAMHB without Tween 80. Red shading indicates bedaquiline (BDQ) samples, blue shading indicates imipenem (IPM) samples, and gray shading indicates IPM + BDQ samples.

| Strain, condition,<br>Day 0 bacterial burden,<br>figure number | Drug(s) and<br>concentration(s)<br>in µg/mL | Log<sub>10</sub> CFU/mL at the<br>following time points: |  | Log<sub>10</sub> RLU/mL at the<br>following time points: |  |  | MegaRLU/CFU at the<br>following time points: |  |
| --- | --- | --- | --- | --- | --- | --- | --- | --- |
|  |  | Day 1 | Day 3 | Hour 4 | Day 1 | Day 3 | Day 1 | Day 3 |
| ATCC 19977 WT,<br>CAMHB (no Tween),<br>Day 0 = 5.95 log<sub>10</sub> CFU/mL,<br>Fig. S4 (replicate 2) | No drug | 7.01 | Clump | 1.87 | 2.90 | 4.04 | 77 | nd |
|  | IPM 4 | 5.68 | 5.29 | 2.72 | 3.78 | 4.11 | 12490 | 65650 |
|  | IPM 16 | 5.02 | 4.68 | 3.09 | 3.65 | 3.66 | 43291 | 94145 |
|  | BDQ 0.00390625 | 6.66 | Clump | 2.38 | 3.17 | 3.74 | 323 | nd |
|  | BDQ 0.0078125 | 6.35 | Clump | 2.41 | 3.00 | 3.93 | 443 | nd |
|  | BDQ 0.015625 | 5.98 | 6.41 | 2.17 | 2.51 | 4.05 | 341 | 4341 |
|  | BDQ 0.03125 | 5.81 | 5.78 | 1.28 | 2.19 | 3.38 | 243 | 4007 |
|  | BDQ 0.0625 | 5.76 | 5.76 | 1.50 | 2.24 | 2.42 | 296 | 449 |
|  | BDQ 0.125 | 5.70 | 5.72 | 1.56 | 2.06 | 2.38 | 231 | 459 |
|  | BDQ 0.25 | 5.70 | 5.60 | 1.95 | 1.89 | 2.30 | 154 | 502 |
|  | BDQ 0.5 | 5.66 | 5.34 | not detected | 1.91 | 2.24 | 176 | 789 |
|  | BDQ 1 | 5.75 | 5.29 | not detected | 1.87 | 2.08 | 131 | 613 |
|  | BDQ 4 | 5.58 | 5.32 | not detected | 1.64 | 1.66 | 114 | 217 |
|  | IPM 4 + BDQ 0.00390625 | 5.70 | 5.13 | 2.53 | 3.47 | 3.62 | 5864 | 30496 |
|  | IPM 4 + BDQ 0.0078125 | 5.48 | 5.62 | 2.69 | 3.41 | 3.43 | 8651 | 6400 |
|  | IPM 4 + BDQ 0.015625 | 5.76 | 5.48 | 2.10 | 3.01 | 3.16 | 1781 | 4850 |
|  | IPM 4 + BDQ 0.03125 | 5.92 | 5.68 | 1.64 | 2.59 | 2.94 | 464 | 1813 |
|  | IPM 4 + BDQ 0.0625 | 5.66 | 5.78 | 2.14 | 2.32 | 2.61 | 455 | 682 |
|  | IPM 4 + BDQ 0.125 | 5.70 | 5.60 | 2.10 | 2.20 | 2.44 | 321 | 684 |
|  | IPM 4 + BDQ 0.25 | 5.75 | 5.28 | 2.21 | 2.04 | 2.68 | 194 | 2479 |
|  | IPM 4 + BDQ 0.5 | 5.70 | 5.32 | 1.73 | 2.14 | 2.10 | 276 | 606 |
|  | IPM 4 + BDQ 1 | 5.87 | 5.30 | 1.70 | 1.85 | 2.04 | 96 | 545 |
|  | IPM 4 + BDQ 4 | 5.48 | 5.24 | 1.30 | 1.72 | 1.87 | 173 | 427 |
|  | IPM 16 + BDQ 0.00390625 | 5.68 | 5.53 | 2.87 | 3.27 | 3.30 | 3859 | 5841 |
|  | IPM 16 + BDQ 0.0078125 | 5.75 | 5.51 | 2.85 | 3.20 | 3.14 | 2839 | 4351 |
|  | IPM 16 + BDQ 0.015625 | 5.75 | 5.51 | 2.54 | 2.95 | 2.99 | 1604 | 3050 |
|  | IPM 16 + BDQ 0.03125 | 5.81 | 5.66 | 1.78 | 2.73 | 2.86 | 833 | 1558 |
|  | IPM 16 + BDQ 0.0625 | 5.78 | 5.68 | 2.05 | 2.29 | 2.69 | 322 | 1011 |
|  | IPM 16 + BDQ 0.125 | 5.64 | 5.73 | 1.81 | 2.02 | 2.47 | 240 | 549 |
|  | IPM 16 + BDQ 0.25 | 5.78 | 5.56 | 2.13 | 2.10 | 2.36 | 212 | 642 |
|  | IPM 16 + BDQ 0.5 | 5.68 | 5.64 | 1.73 | 2.10 | 2.18 | 260 | 347 |
|  | IPM 16 + BDQ 1 | 5.82 | 5.14 | 1.61 | 1.98 | 2.15 | 145 | 1020 |
|  | IPM 16 + BDQ 4 | 5.64 | 5.09 | 1.68 | 1.66 | 2.02 | 103 | 838 |

"Clump" indicates that the bacteria had overgrown and formed clumps, precluding CFU quantification. nd indicates not determined (due to lack of CFU count data).

**Table S10.** CFU and relative fluorescence unit (RLU) data from samples presented in Fig. 7, Fig. S6, Fig. S7, and Fig. S8. Bacteria were nutrient-starved in PBS for 14 days (NS-14) prior to drug exposure. Drug activity assays were performed in PBS without Tween 80. Red shading indicates bedaquiline (BDQ) samples, blue shading indicates imipenem (IPM) samples, and gray shading indicates IPM + BDQ samples.

| Strain, condition,<br>Day 0 bacterial burden,<br>figure number | Drug(s) and<br>concentration(s)<br>in µg/mL | Log<sub>10</sub> CFU/mL at the<br>following time points: |  | Log<sub>10</sub> RLU/mL at the<br>following time points: |  |  |  | MegaRLU/CFU at the<br>following time points: |  |
| --- | --- | --- | --- | --- | --- | --- | --- | --- | --- |
|  |  | Day 3 | Day 7 | Hour 4 | Day 1 | Day 3 | Day 7 | Day 3 | Day 7 |
| **ATCC 19977 WT,**<br>NS-14 (no Tween),<br>Day 0 = 6.25 log<sub>10</sub> CFU/mL,<br>**Fig. 7A,C; Fig. S6A,C;**<br>**Fig. S7** | No drug | 6.26 | 6.20 | 2.44 | 2.39 | 2.40 | 2.58 | 135 | 241 |
|  | IPM 4 | 6.12 | 6.68 | 2.45 | 2.47 | 2.60 | 2.91 | 302 | 170 |
|  | IPM 16 | 6.09 | 5.95 | 2.49 | 2.52 | 2.65 | 2.92 | 357 | 918 |
|  | BDQ 0.00390625 | 6.23 | 6.07 | 2.52 | 2.44 | 2.47 | 2.62 | 178 | 355 |
|  | BDQ 0.0078125 | 6.13 | 6.20 | 2.45 | 2.41 | 2.45 | 2.63 | 208 | 264 |
|  | BDQ 0.015625 | 6.18 | 6.23 | 2.48 | 2.46 | 2.43 | 2.62 | 178 | 250 |
|  | BDQ 0.03125 | 6.17 | 6.18 | 2.37 | 2.25 | 2.31 | 2.48 | 138 | 200 |
|  | BDQ 0.0625 | 6.09 | 6.18 | 2.34 | 2.24 | 2.25 | 2.46 | 145 | 188 |
|  | BDQ 0.125 | 5.90 | 6.53 | 2.32 | 2.18 | 2.23 | 2.42 | 213 | 78 |
|  | BDQ 0.25 | 5.53 | 5.36 | 2.49 | 2.18 | 2.20 | 2.34 | 471 | 953 |
|  | BDQ 0.5 | 5.13 | 4.48 | 2.34 | 2.18 | 2.21 | 2.31 | 1182 | 6752 |
|  | BDQ 1 | 4.16 | 3.89 | 2.34 | 2.20 | 2.21 | 2.33 | 11142 | 27340 |
|  | BDQ 4 | 2.21 | 0.95 | 2.29 | 2.17 | 2.18 | 2.30 | 938750 | 25137500 |
|  | IPM 4 + BDQ 0.00390625 | 5.93 | 6.48 | 2.44 | 2.50 | 2.63 | 2.84 | 499 | 229 |
|  | IPM 4 + BDQ 0.0078125 | 6.28 | 6.56 | 2.42 | 2.47 | 2.61 | 2.86 | 213 | 203 |
|  | IPM 4 + BDQ 0.015625 | 5.83 | 6.48 | 2.38 | 2.43 | 2.58 | 2.88 | 565 | 255 |
|  | IPM 4 + BDQ 0.03125 | 6.07 | 6.38 | 2.31 | 2.26 | 2.33 | 2.87 | 183 | 307 |
|  | IPM 4 + BDQ 0.0625 | 6.00 | 6.51 | 2.34 | 2.25 | 2.36 | 2.77 | 230 | 186 |
|  | IPM 4 + BDQ 0.125 | 5.93 | 6.17 | 2.27 | 2.21 | 2.25 | 2.60 | 206 | 270 |
|  | IPM 4 + BDQ 0.25 | 5.64 | 5.22 | 2.28 | 2.18 | 2.23 | 2.46 | 382 | 1730 |
|  | IPM 4 + BDQ 0.5 | 5.06 | 4.37 | 2.33 | 2.18 | 2.23 | 2.34 | 1450 | 9322 |
|  | IPM 4 + BDQ 1 | 3.03 | 4.27 | 2.35 | 2.20 | 2.20 | 2.22 | 147685 | 8750 |
|  | IPM 4 + BDQ 4 | 2.15 | 2.38 | 2.32 | 2.18 | 2.21 | 2.21 | 1158214 | 675208 |
|  | IPM 16 + BDQ 0.00390625 | 6.24 | 5.89 | 2.41 | 2.50 | 2.58 | 2.79 | 223 | 782 |
|  | IPM 16 + BDQ 0.0078125 | 6.33 | 5.76 | 2.52 | 2.48 | 2.54 | 2.78 | 163 | 1051 |
|  | IPM 16 + BDQ 0.015625 | 6.18 | 5.75 | 2.33 | 2.38 | 2.45 | 2.66 | 186 | 808 |
|  | IPM 16 + BDQ 0.03125 | 5.91 | 5.93 | 2.47 | 2.29 | 2.41 | 2.64 | 311 | 503 |
|  | IPM 16 + BDQ 0.0625 | 5.91 | 5.28 | 2.33 | 2.22 | 2.27 | 2.43 | 228 | 1412 |
|  | IPM 16 + BDQ 0.125 | 5.68 | 5.07 | 2.27 | 2.19 | 2.21 | 2.39 | 336 | 2059 |
|  | IPM 16 + BDQ 0.25 | 5.41 | 4.86 | 2.31 | 2.18 | 2.25 | 2.38 | 679 | 3347 |
|  | IPM 16 + BDQ 0.5 | 4.90 | 4.48 | 2.32 | 2.18 | 2.22 | 2.33 | 2096 | 7125 |
|  | IPM 16 + BDQ 1 | 3.82 | 3.70 | 2.33 | 2.19 | 2.17 | 2.33 | 22515 | 42660 |
|  | IPM 16 + BDQ 4 | 1.79 | 0.95 | 2.36 | 2.15 | 2.19 | 2.32 | 2594167 | 25987500 |

**Table S10, continued.**
| Strain, condition,<br>Day 0 bacterial burden,<br>figure number | Drug(s) and<br>concentration(s)<br>in µg/mL | Log<sub>10</sub> CFU/mL at the<br>following time points: |  | Log<sub>10</sub> RLU/mL at the<br>following time points: |  |  |  | MegaRLU/CFU at the<br>following time points: |  |
| --- | --- | --- | --- | --- | --- | --- | --- | --- | --- |
|  |  | Day 3 | Day 7 | Hour 4 | Day 1 | Day 3 | Day 7 | Day 3 | Day 7 |
| **OM7 mutant,**<br>NS-14 (no Tween),<br>Day 0 = 6.29 log<sub>10</sub> CFU/mL,<br>**Fig. 7B,D; Fig. S6B,D;**<br>**Fig. S8** | No drug | 6.42 | 6.82 | 2.30 | 2.34 | 2.31 | 2.52 | 78 | 50 |
|  | IPM 4 | 6.43 | 6.58 | 2.36 | 2.47 | 2.61 | 2.91 | 150 | 215 |
|  | IPM 16 | 6.26 | 6.12 | 2.41 | 2.46 | 2.51 | 2.85 | 174 | 536 |
|  | BDQ 0.00390625 | 6.31 | 6.32 | 2.40 | 2.45 | 2.36 | 2.53 | 113 | 162 |
|  | BDQ 0.0078125 | 6.23 | 6.27 | 2.35 | 2.29 | 2.29 | 2.48 | 115 | 159 |
|  | BDQ 0.015625 | 6.30 | 5.91 | 2.48 | 2.41 | 2.41 | 2.58 | 128 | 468 |
|  | BDQ 0.03125 | 6.26 | 6.19 | 2.35 | 2.35 | 2.32 | 2.52 | 113 | 213 |
|  | BDQ 0.0625 | 6.20 | 6.26 | 2.30 | 2.22 | 2.25 | 2.43 | 112 | 148 |
|  | BDQ 0.125 | 6.15 | 6.21 | 2.28 | 2.14 | 2.18 | 2.42 | 107 | 161 |
|  | BDQ 0.25 | 6.11 | 6.03 | 2.22 | 2.10 | 2.09 | 2.31 | 95 | 192 |
|  | BDQ 0.5 | 6.00 | 6.02 | 2.21 | 2.13 | 2.14 | 2.35 | 138 | 215 |
|  | BDQ 1 | 5.83 | 5.51 | 2.22 | 2.12 | 2.09 | 2.24 | 182 | 544 |
|  | BDQ 4 | 4.51 | 3.43 | 2.23 | 2.12 | 2.09 | 2.19 | 3838 | 57854 |
|  | IPM 4 + BDQ 0.00390625 | 6.25 | 6.25 | 2.39 | 2.50 | 2.56 | 2.84 | 206 | 390 |
|  | IPM 4 + BDQ 0.0078125 | 6.25 | 6.08 | 2.37 | 2.43 | 2.49 | 2.87 | 175 | 612 |
|  | IPM 4 + BDQ 0.015625 | 6.26 | 6.56 | 2.39 | 2.50 | 2.53 | 2.88 | 188 | 212 |
|  | IPM 4 + BDQ 0.03125 | 6.25 | 5.94 | 2.29 | 2.42 | 2.45 | 2.87 | 161 | 838 |
|  | IPM 4 + BDQ 0.0625 | 6.26 | 6.19 | 2.29 | 2.34 | 2.37 | 2.77 | 131 | 386 |
|  | IPM 4 + BDQ 0.125 | 6.11 | 6.30 | 2.25 | 2.21 | 2.26 | 2.60 | 143 | 200 |
|  | IPM 4 + BDQ 0.25 | 5.89 | 6.47 | 2.21 | 2.19 | 2.19 | 2.46 | 200 | 98 |
|  | IPM 4 + BDQ 0.5 | 6.06 | 6.20 | 2.20 | 2.14 | 2.09 | 2.34 | 109 | 138 |
|  | IPM 4 + BDQ 1 | 5.53 | 5.26 | 2.24 | 2.10 | 2.09 | 2.22 | 359 | 894 |
|  | IPM 4 + BDQ 4 | 4.86 | 3.68 | 2.25 | 2.11 | 2.07 | 2.21 | 1644 | 33760 |
|  | IPM 16 + BDQ 0.00390625 | 6.33 | 6.05 | 2.38 | 2.48 | 2.49 | 2.88 | 143 | 673 |
|  | IPM 16 + BDQ 0.0078125 | 6.12 | 5.87 | 2.34 | 2.44 | 2.46 | 2.84 | 219 | 936 |
|  | IPM 16 + BDQ 0.015625 | 6.11 | 5.97 | 2.39 | 2.43 | 2.45 | 2.76 | 220 | 606 |
|  | IPM 16 + BDQ 0.03125 | 6.20 | 5.79 | 2.37 | 2.33 | 2.38 | 2.69 | 153 | 799 |
|  | IPM 16 + BDQ 0.0625 | 6.09 | 5.68 | 2.31 | 2.26 | 2.32 | 2.63 | 170 | 880 |
|  | IPM 16 + BDQ 0.125 | 5.81 | 5.60 | 2.23 | 2.19 | 2.26 | 2.44 | 281 | 693 |
|  | IPM 16 + BDQ 0.25 | 5.86 | 5.45 | 2.26 | 2.21 | 2.17 | 2.38 | 205 | 851 |
|  | IPM 16 + BDQ 0.5 | 5.87 | 5.16 | 2.26 | 2.15 | 2.18 | 2.31 | 202 | 1401 |
|  | IPM 16 + BDQ 1 | 5.58 | 4.66 | 2.21 | 2.10 | 2.09 | 2.23 | 327 | 3682 |
|  | IPM 16 + BDQ 4 | 4.89 | 3.79 | 2.25 | 2.19 | 2.13 | 2.22 | 1748 | 26782 |

**Table S11.** CFU data from samples presented in Fig. 8 and Fig. S9. Bacteria were nutrient-starved in PBS for 20 days (NS-20) prior to drug exposure. Drug activity assays were performed in PBS with 0.05% Tween 80. Red shading indicates bedaquiline (BDQ) samples, blue shading indicates imipenem (IPM) samples, and gray shading indicates IPM + BDQ samples.

| Strain, condition,<br>Day 0 bacterial burden,<br>figure number | Drug(s) and<br>concentration(s)<br>in µg/mL | Log<sub>10</sub> CFU/mL at the<br>following time points: |  |
| --- | --- | --- | --- |
|  |  | Day 1 | Day 4 |
| ATCC 19977 WT,<br>NS-20 + Tween,<br>Day 0 = 6.07 log<sub>10</sub> CFU/mL,<br>**Fig. 8; Fig. S9A,C**<br>(replicate 1) | No drug | 5.68 | 5.48 |
|  | IPM 4 | 5.33 | 2.27 |
|  | IPM 8 | 4.26 | 1.36 |
|  | IPM 16 | 4.60 | 1.18 |
|  | BDQ 0.00390625 | 5.56 | 5.96 |
|  | BDQ 0.0625 | 5.37 | 5.51 |
|  | BDQ 1 | 5.32 | 5.04 |
|  | BDQ 4 | 5.13 | 4.66 |
|  | IPM 4 + BDQ 0.00390625 | 5.62 | 4.26 |
|  | IPM 4 + BDQ 0.0625 | 5.33 | 4.90 |
|  | IPM 4 + BDQ 0.25 | 5.35 | 5.23 |
|  | IPM 4 + BDQ 1 | 5.15 | 4.95 |
|  | IPM 4 + BDQ 4 | 5.08 | 4.56 |
| ATCC 19977 WT,<br>NS-20 + Tween,<br>Day 0 = 6.38 log<sub>10</sub> CFU/mL,<br>**Fig. S9B,D** (replicate 2) | No drug | 6.38 | 6.23 |
|  | IPM 4 | 5.97 | Contam. |
|  | IPM 8 | 5.60 | 2.08 |
|  | IPM 16 | 4.88 | 0.00 |
|  | BDQ 0.00390625 | 6.37 | 6.22 |
|  | BDQ 0.0625 | 5.94 | 6.21 |
|  | BDQ 1 | 6.04 | 5.89 |
|  | BDQ 4 | 6.05 | 5.37 |
|  | IPM 4 + BDQ 0.00390625 | 6.15 | 5.89 |
|  | IPM 4 + BDQ 0.0625 | 6.03 | 5.94 |
|  | IPM 4 + BDQ 0.25 | 6.11 | 5.56 |
|  | IPM 4 + BDQ 1 | 6.06 | 5.81 |
|  | IPM 4 + BDQ 4 | 6.01 | 5.29 |

"Contam." indicates that bacterial or fungal contamination precluded assessment of this sample.

**Table S12.**
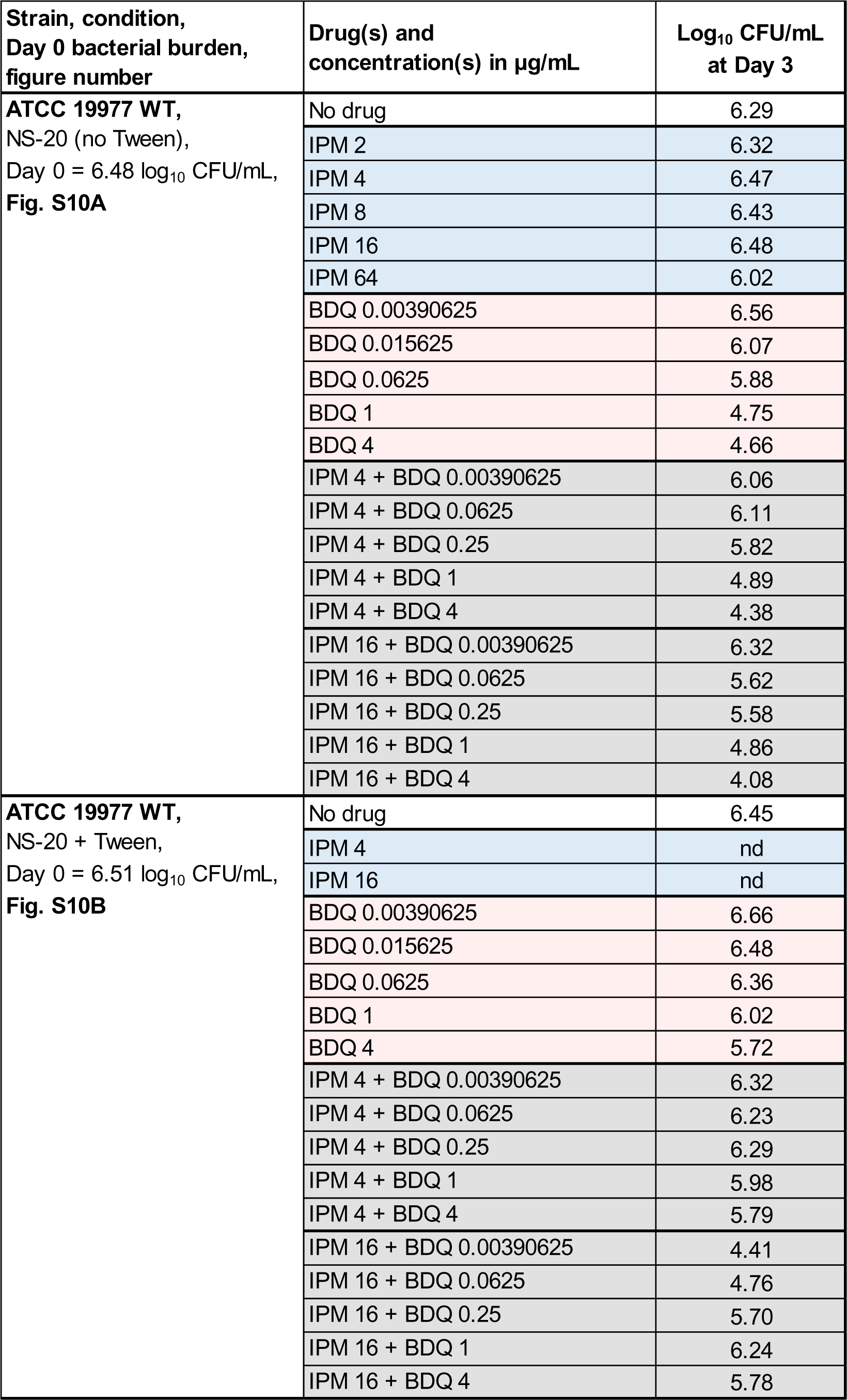
CFU data from samples presented in Fig. S10. Bacteria were nutrient-starved in PBS for 20 days (NS-20) prior to drug exposure. Drug activity assays were performed in PBS with or without 0.05% Tween 80. Red shading indicates bedaquiline (BDQ) samples; blue shading indicates imipenem (IPM) samples; gray shading indicates IPM + BDQ samples.

**Table S13.**
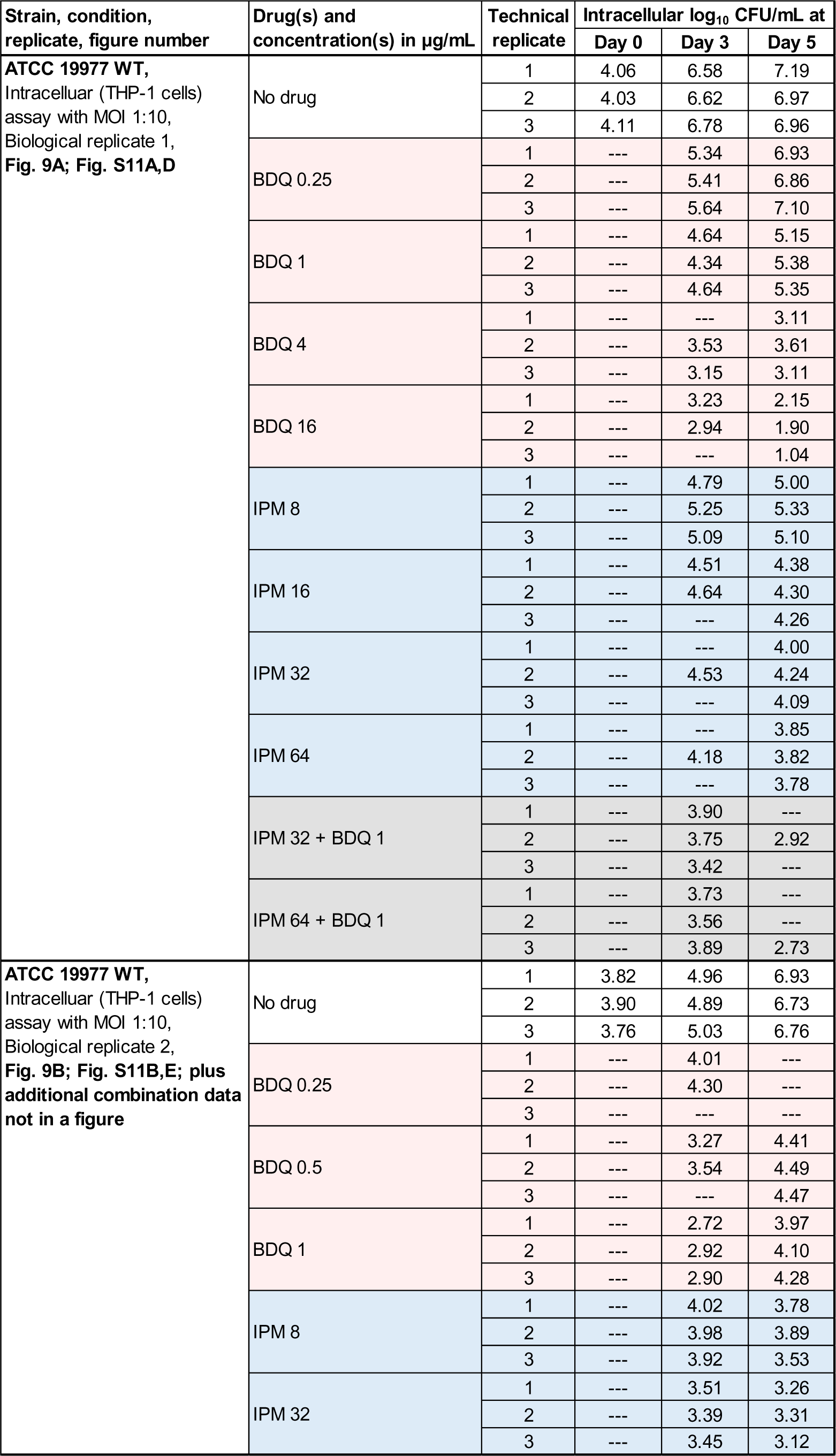
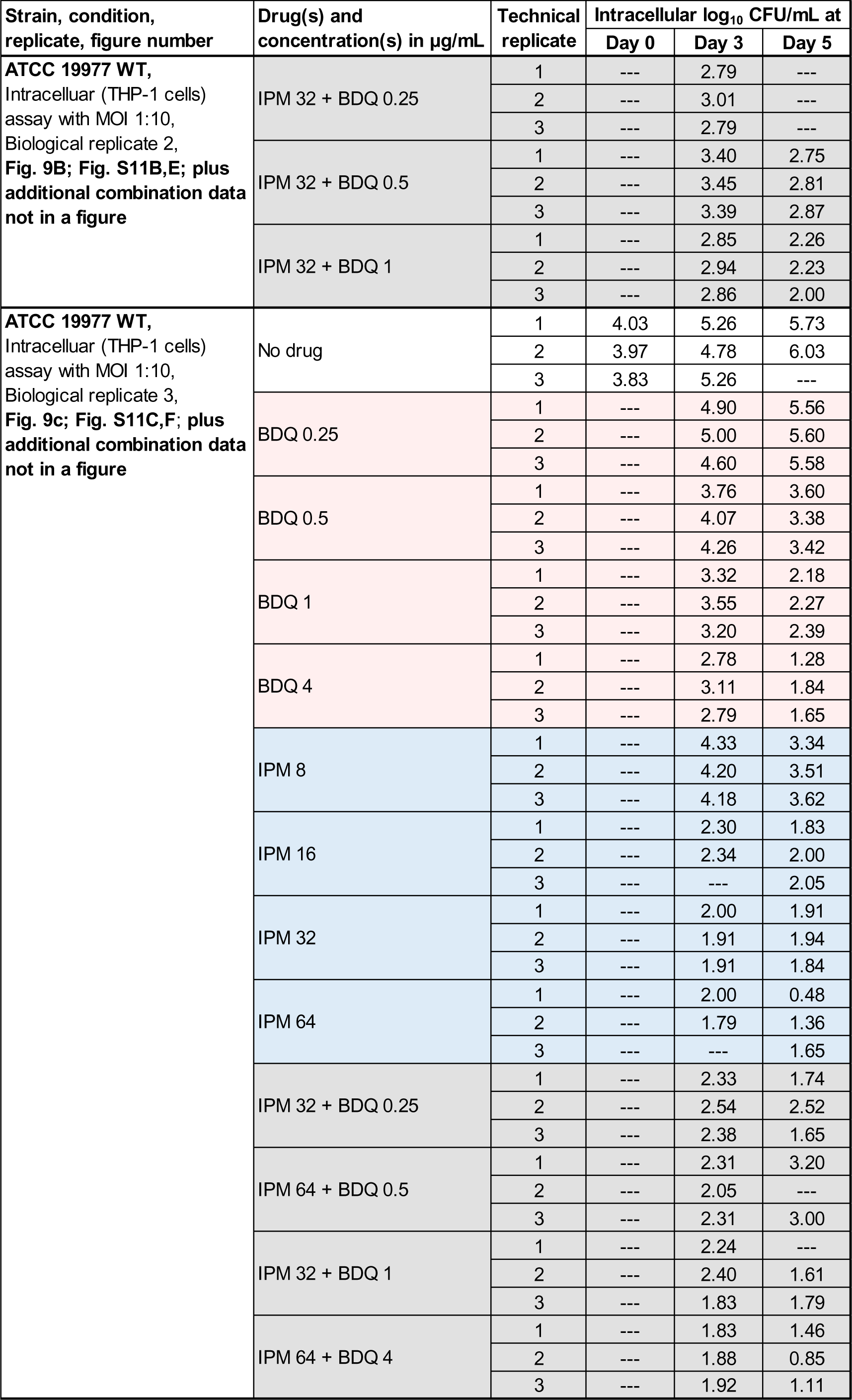
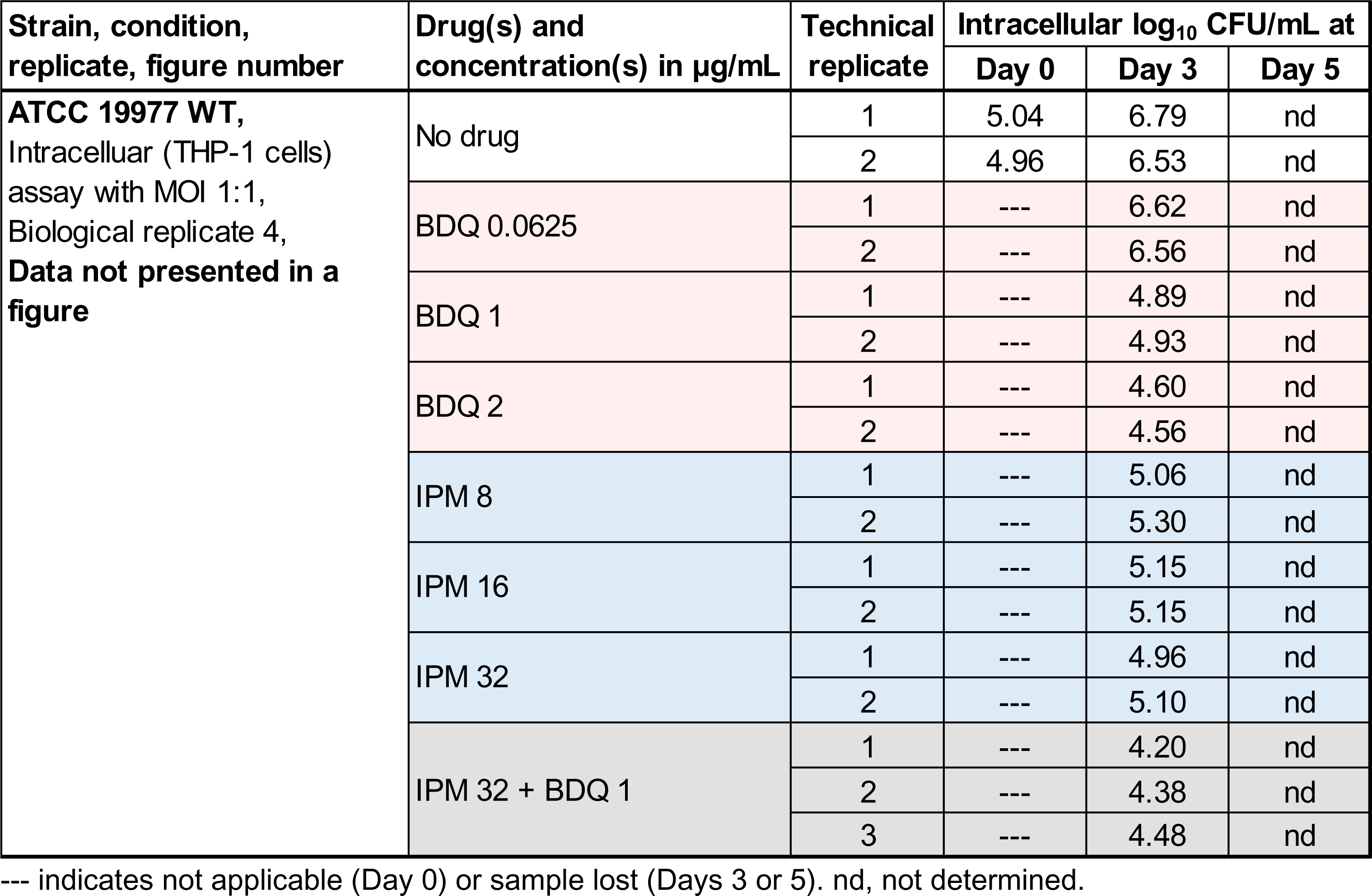
CFU data from intracellular samples presented in Fig. 9 and Fig. S11. Within each biological replicate, data from up to 3 technical replicates were acquired for each group at each time point. Red shading indicates bedaquiline (BDQ) samples; blue shading indicates imipenem (IPM) samples; gray shading indicates IPM + BDQ samples.

| Strain, condition, replicate, figure number | Drug(s) and concentration(s) in µg/mL | Technical replicate | Intracellular log<sub>10</sub> CFU/mL at |  |  |
| --- | --- | --- | --- | --- | --- |
|  |  |  | Day 0 | Day 3 | Day 5 |
| **ATCC 19977 WT,**<br>Intracellular (THP-1 cells)<br>assay with MOI 1:10,<br>Biological replicate 1,<br>**Fig. 9A; Fig. S11A,D** | No drug | 1 | 4.06 | 6.58 | 7.19 |
|  |  | 2 | 4.03 | 6.62 | 6.97 |
|  |  | 3 | 4.11 | 6.78 | 6.96 |
|  | BDQ 0.25 | 1 | --- | 5.34 | 6.93 |
|  |  | 2 | --- | 5.41 | 6.86 |
|  |  | 3 | --- | 5.64 | 7.10 |
|  | BDQ 1 | 1 | --- | 4.64 | 5.15 |
|  |  | 2 | --- | 4.34 | 5.38 |
|  |  | 3 | --- | 4.64 | 5.35 |
|  | BDQ 4 | 1 | --- | --- | 3.11 |
|  |  | 2 | --- | 3.53 | 3.61 |
|  |  | 3 | --- | 3.15 | 3.11 |
|  | BDQ 16 | 1 | --- | 3.23 | 2.15 |
|  |  | 2 | --- | 2.94 | 1.90 |
|  |  | 3 | --- | --- | 1.04 |
|  | IPM 8 | 1 | --- | 4.79 | 5.00 |
|  |  | 2 | --- | 5.25 | 5.33 |
|  |  | 3 | --- | 5.09 | 5.10 |
|  | IPM 16 | 1 | --- | 4.51 | 4.38 |
|  |  | 2 | --- | 4.64 | 4.30 |
|  |  | 3 | --- | --- | 4.26 |
|  | IPM 32 | 1 | --- | --- | 4.00 |
|  |  | 2 | --- | 4.53 | 4.24 |
|  |  | 3 | --- | --- | 4.09 |
|  | IPM 64 | 1 | --- | --- | 3.85 |
|  |  | 2 | --- | 4.18 | 3.82 |
|  |  | 3 | --- | --- | 3.78 |
|  | IPM 32 + BDQ 1 | 1 | --- | 3.90 | --- |
|  |  | 2 | --- | 3.75 | 2.92 |
|  |  | 3 | --- | 3.42 | --- |
|  | IPM 64 + BDQ 1 | 1 | --- | 3.73 | --- |
|  |  | 2 | --- | 3.56 | --- |
|  |  | 3 | --- | 3.89 | 2.73 |
| **ATCC 19977 WT,**<br>Intracellular (THP-1 cells)<br>assay with MOI 1:10,<br>Biological replicate 2,<br>**Fig. 9B; Fig. S11B,E; plus**<br>**additional combination data**<br>**not in a figure** | No drug | 1 | 3.82 | 4.96 | 6.93 |
|  |  | 2 | 3.90 | 4.89 | 6.73 |
|  |  | 3 | 3.76 | 5.03 | 6.76 |
|  | BDQ 0.25 | 1 | --- | 4.01 | --- |
|  |  | 2 | --- | 4.30 | --- |
|  |  | 3 | --- | --- | --- |
|  | BDQ 0.5 | 1 | --- | 3.27 | 4.41 |
|  |  | 2 | --- | 3.54 | 4.49 |
|  |  | 3 | --- | --- | 4.47 |
|  | BDQ 1 | 1 | --- | 2.72 | 3.97 |
|  |  | 2 | --- | 2.92 | 4.10 |
|  |  | 3 | --- | 2.90 | 4.28 |
|  | IPM 8 | 1 | --- | 4.02 | 3.78 |
|  |  | 2 | --- | 3.98 | 3.89 |
|  |  | 3 | --- | 3.92 | 3.53 |
|  | IPM 32 | 1 | --- | 3.51 | 3.26 |
|  |  | 2 | --- | 3.39 | 3.31 |
|  |  | 3 | --- | 3.45 | 3.12 |
|  |  |  | Day 0 | Day 3 | Day 5 |
| ATCC 19977 WT,<br>Intracellular (THP-1 cells)<br>assay with MOI 1:10,<br>Biological replicate 2,<br>**Fig. 9B; Fig. S11B,E; plus**<br>**additional combination data**<br>**not in a figure** | IPM 32 + BDQ 0.25 | 1 | --- | 2.79 | --- |
|  |  | 2 | --- | 3.01 | --- |
|  |  | 3 | --- | 2.79 | --- |
|  | IPM 32 + BDQ 0.5 | 1 | --- | 3.40 | 2.75 |
|  |  | 2 | --- | 3.45 | 2.81 |
|  |  | 3 | --- | 3.39 | 2.87 |
|  | IPM 32 + BDQ 1 | 1 | --- | 2.85 | 2.26 |
|  |  | 2 | --- | 2.94 | 2.23 |
|  |  | 3 | --- | 2.86 | 2.00 |
| ATCC 19977 WT,<br>Intracellular (THP-1 cells)<br>assay with MOI 1:10,<br>Biological replicate 3,<br>**Fig. 9c; Fig. S11C,F; plus**<br>**additional combination data**<br>**not in a figure** | No drug | 1 | 4.03 | 5.26 | 5.73 |
|  |  | 2 | 3.97 | 4.78 | 6.03 |
|  |  | 3 | 3.83 | 5.26 | --- |
|  | BDQ 0.25 | 1 | --- | 4.90 | 5.56 |
|  |  | 2 | --- | 5.00 | 5.60 |
|  |  | 3 | --- | 4.60 | 5.58 |
|  | BDQ 0.5 | 1 | --- | 3.76 | 3.60 |
|  |  | 2 | --- | 4.07 | 3.38 |
|  |  | 3 | --- | 4.26 | 3.42 |
|  | BDQ 1 | 1 | --- | 3.32 | 2.18 |
|  |  | 2 | --- | 3.55 | 2.27 |
|  |  | 3 | --- | 3.20 | 2.39 |
|  | BDQ 4 | 1 | --- | 2.78 | 1.28 |
|  |  | 2 | --- | 3.11 | 1.84 |
|  |  | 3 | --- | 2.79 | 1.65 |
|  | IPM 8 | 1 | --- | 4.33 | 3.34 |
|  |  | 2 | --- | 4.20 | 3.51 |
|  |  | 3 | --- | 4.18 | 3.62 |
|  | IPM 16 | 1 | --- | 2.30 | 1.83 |
|  |  | 2 | --- | 2.34 | 2.00 |
|  |  | 3 | --- | --- | 2.05 |
|  | IPM 32 | 1 | --- | 2.00 | 1.91 |
|  |  | 2 | --- | 1.91 | 1.94 |
|  |  | 3 | --- | 1.91 | 1.84 |
|  | IPM 64 | 1 | --- | 2.00 | 0.48 |
|  |  | 2 | --- | 1.79 | 1.36 |
|  |  | 3 | --- | --- | 1.65 |
|  | IPM 32 + BDQ 0.25 | 1 | --- | 2.33 | 1.74 |
|  |  | 2 | --- | 2.54 | 2.52 |
|  |  | 3 | --- | 2.38 | 1.65 |
|  | IPM 64 + BDQ 0.5 | 1 | --- | 2.31 | 3.20 |
|  |  | 2 | --- | 2.05 | --- |
|  |  | 3 | --- | 2.31 | 3.00 |
|  | IPM 32 + BDQ 1 | 1 | --- | 2.24 | --- |
|  |  | 2 | --- | 2.40 | 1.61 |
|  |  | 3 | --- | 1.83 | 1.79 |
|  | IPM 64 + BDQ 4 | 1 | --- | 1.83 | 1.46 |
|  |  | 2 | --- | 1.88 | 0.85 |
|  |  | 3 | --- | 1.92 | 1.11 |
|  |  |  | Day 0 | Day 3 | Day 5 |
| ATCC 19977 WT,<br>Intracellular (THP-1 cells)<br>assay with MOI 1:1,<br>Biological replicate 4,<br>Data not presented in a figure | No drug | 1 | 5.04 | 6.79 | nd |
|  |  | 2 | 4.96 | 6.53 | nd |
|  | BDQ 0.0625 | 1 | --- | 6.62 | nd |
|  |  | 2 | --- | 6.56 | nd |
|  | BDQ 1 | 1 | --- | 4.89 | nd |
|  |  | 2 | --- | 4.93 | nd |
|  | BDQ 2 | 1 | --- | 4.60 | nd |
|  |  | 2 | --- | 4.56 | nd |
|  | IPM 8 | 1 | --- | 5.06 | nd |
|  |  | 2 | --- | 5.30 | nd |
|  | IPM 16 | 1 | --- | 5.15 | nd |
|  |  | 2 | --- | 5.15 | nd |
|  | IPM 32 | 1 | --- | 4.96 | nd |
|  |  | 2 | --- | 5.10 | nd |
|  | IPM 32 + BDQ 1 | 1 | --- | 4.20 | nd |
|  |  | 2 | --- | 4.38 | nd |
|  |  | 3 | --- | 4.48 | nd |

--- indicates not applicable (Day 0) or sample lost (Days 3 or 5). nd, not determined.

